# P53-independent restoration of p53 pathway in tumors with mutated p53 through ATF4 transcriptional modulation by ERK1/2 and CDK9

**DOI:** 10.1101/2020.10.20.347401

**Authors:** Xiaobing Tian, Nagib Ahsan, Amriti Lulla, Avital Lev, Philip Abbosh, David T. Dicker, Shengliang Zhang, Wafik S. El-Deiry

## Abstract

A long-term goal in the cancer-field has been to develop strategies for treating p53-mutated tumors. A novel small-molecule, PG3-Oc, restores p53 pathway-signaling in tumor cells with mutant-p53, independently of p53/p73. PG3-Oc partially upregulates the p53-transcriptome (13.7% of public p53 target-gene dataset; 15.2% of in-house dataset) and p53-proteome (18%, HT29; 16%, HCT116-p53^-/-^). Bioinformatic analysis indicates critical p53-effectors of growth-arrest (p21), apoptosis (PUMA, DR5, Noxa), autophagy (DRAM1), and metastasis-suppression (NDRG1) are induced by PG3-Oc. ERK1/2- and CDK9-kinases are required to upregulate ATF4 by PG3-Oc which restores p53 transcriptomic-targets in cells without functional-p53. PG3-Oc represses MYC (ATF4-independent), and upregulates PUMA (ATF4-dependent) in mediating cell death. With largely nonoverlapping transcriptomes, induced-ATF4 restores p53 transcriptomic targets in drug-treated cells including functionally important mediators such as PUMA and DR5. Our results demonstrate novel p53-independent drug-induced molecular reprogramming involving ERK1/2, CDK9, and ATF4 to restore upregulation of p53 effector genes required for cell death and tumor suppression.

## Introduction

The p53 transcription factor is activated by various cellular stresses such as DNA damage, oncogene activation, nutrient depletion, oxidative stress and endoplasmic reticulum (ER) stress (Bykov et al., 2018; Lin et al., 2012; Vousden and Prives, 2009). The tumor suppressor p53 regulates complicated transcription programs that respond to particular stress signals in maintaining homeostasis and guarding the genome. There are three main outcomes after the activation of p53: cell-cycle arrest, senescence and apoptosis. Cell-cycle arrest allows cell repair and recovery from the stress, so that cell survival occurs. Senescence and apoptosis are terminal and irreversible. It has been proposed that the nature of the stress signal, the nature of the compound, the duration of the stress signal and the cell type determine the transcriptional program of p53, and the phenotypic outcome in the stressed cell (Andrysik et al., 2017; Riley et al., 2008). Thus, when p53 is activated, a specific set of p53 target genes is regulated, instead of all of the p53 target genes, with tissue specificity (Fei et al., 2002). In addition, specific gene sets of p53-activated changes occur over time in cells following a stressor. For example, HCT116 and HCT116 p53^-/-^ cells treated with the MDM2 inhibitor nutlin-3 for 1 hour, followed by global run-on sequencing (GRO-Seq), identified 198 possible direct targets of p53 (Allen et al., 2014). Both CHIP-seq and RNA-seq analysis identified 432 direct p53 target genes in mouse MEF cells treated with the DNA damage-inducing drug doxorubicin (Kenzelmann Broz et al., 2013). Menendez *et al*. employed CHIP-seq and microarray analysis and identified 205 p53 target genes in U2OS cells. The authors reported that U2OS cells had strikingly different p53-binding patterns and transcriptional responses following exposure to the DNA-damaging agent doxorubicin versus nutlin-3 for 24 hours, with nutlin-3 considered a non-genotoxic activator of p53. Genome-wide analysis of the ChIP-seq identified 3087 p53-binding sites after doxorubicin treatment and nearly 6-fold more sites (18,159) in cells treated with Nutlin-3 (Menendez et al., 2013). Meta-analysis of four publications (using Chip-seq assays) indicated that p53 may directly activate > 1200 genes. However, only 26 of these genes were commonly activated in all four studies (Allen et al., 2014). This lack of overlap is possibly due to methodological differences and cell type-specific differences (Allen et al., 2014). Fischer identified 346 possible direct p53 target genes through searching the literature and performing meta-analysis of data from 319 studies (Fischer, 2017). Because this p53 target gene data-base is not generated from a specific drug in a specific cell line, we selected Fischer’s p53 target genes as a database for evaluating the effectiveness of p53 pathway restoration by a candidate therapeutic compound. We also developed an in-house reference p53 target gene data set by RNA-Seq and p53-dependent protein data set by proteomic analysis, using HCT116 and HCT116 p53^-/-^ cell lines treated with the known p53 activator 5-Fluorouracil (5-FU) as a positive control.

p53 is inactivated in almost all human cancers, either by mutation, deletion, MDM2 overexpression, or inactivation by viral proteins (Mantovani et al., 2019; Vousden and Prives, 2009). Over 50% of human cancers harbor cancer-promoting mutations in p53 (Mantovani et al., 2019). p53 mutations not only abrogate its tumor-suppressor function, but also confer gain-of-function (GOF) properties that contribute to tumorigenesis, proliferation, genomic instability, metabolic remodeling, invasion, metastasis, resistance to apoptosis and cancer therapy resistance (Mantovani et al., 2019; Zhu et al., 2015). Hence, restoration of the p53 pathway represents an important strategy for achieving anti-cancer therapy in mutant p53-bearing tumors. Restoring p53 function has been tried using different approaches, which include restoration of wild-type p53 function through a small molecule that binds to mutant p53 (Bykov et al., 2018; Yu et al., 2012), compound-induced degradation of mutant p53 (Alexandrova et al., 2015; Zhang et al., 2015), disruption of protein-protein interaction between mutant p53 and other transcription factors (Chowdhury et al., 2014). Strategies employing genome-wide restoration of the p53 pathway by small molecules *via* p53-independent mechanisms are considered promising, and may involve the p53 homolog p73 (Hong et al., 2014; Zhang et al., 2015).

About 50% of cancer cells lack wild-type p53 due to mutation or deletion of the TP53 gene, we hypothesized that other transcription factors may compensate for p53 loss and play an important role in coping with extrinsic and intrinsic stresses, and regulate cell fate, such as survival, senescence and apoptosis. Moreover, such factors may be possible to modulate with candidate anti-cancer therapeutics. Like p53, ATF4 (activating transcription factor 4), plays an important role in communicating pro-survival and pro-apoptotic signals. The ER stress kinase PERK (PKR-like endoplasmic reticulum kinase) senses various stresses and catalyzes the phosphorylation of the α-subunit of eIF2α. Phosphorylation of eIF2α at serine 51 attenuates global protein synthesis temporarily while selectively enhancing translation of ATF4 mRNA. Once activated, ATF4 regulates a transcriptional program involved in cell survival (antioxidant response, amino acid biosynthesis and autophagy), senescence and apoptosis. The final outcome of ATF4 activation is dependent on the cell type, nature of stressors and duration of the stresses (Ojha et al., 2019; Tameire et al., 2019; Wang et al., 2015; Wortel et al., 2017).

Prodigiosin is a member of a family of naturally occurring red pigments produced by microorganisms including *Streptomyces* and *Serratia* (Darshan and Manonmani, 2015). Most of the members of this family contain a common 4-methoxy-pyrrolylpyrromethene pharmacophore (Figure 1A, red highlight). Our laboratory reported that prodigiosin shows potent anti-cancer activity against human tumors with mutated p53 through restoring the p53 pathway, in part, via p73 (Hong et al., 2014; Prabhu et al., 2016). Based on the structure of the pharmacophore of the prodigiosin family, we synthesized drug analogs, including a novel compound PG3-Oc whose synthesis is described in an issued composition of matter patent (El-Deiry et al., 2017). We report the effects on cell signaling and anti-cancer activity of PG3-Oc. Importantly, we use PG3-Oc as a chemical probe of the cell signaling mechanisms that underlie p53 pathway restoration in a p53-independent manner. PG3-Oc induces non-canonical ER stress and partially restores the p53 pathway globally through transcription factor ATF4 leading to induction of pro-apoptotic gene targets. Insights into an ERK1/2- and CDK9-dependent ATF4-activation by PG3-Oc leading to cell death and anti-tumor efficacy through PUMA and DR5 are provided. Our results shed light upon the mechanisms used by a novel compound to induce p53-independent p53 pathway restoration in tumors with mutated p53, and demonstrate feasibility of the approach to partially restore the p53 transcriptome globally to achieve tumor suppression.

**Figure 1.**
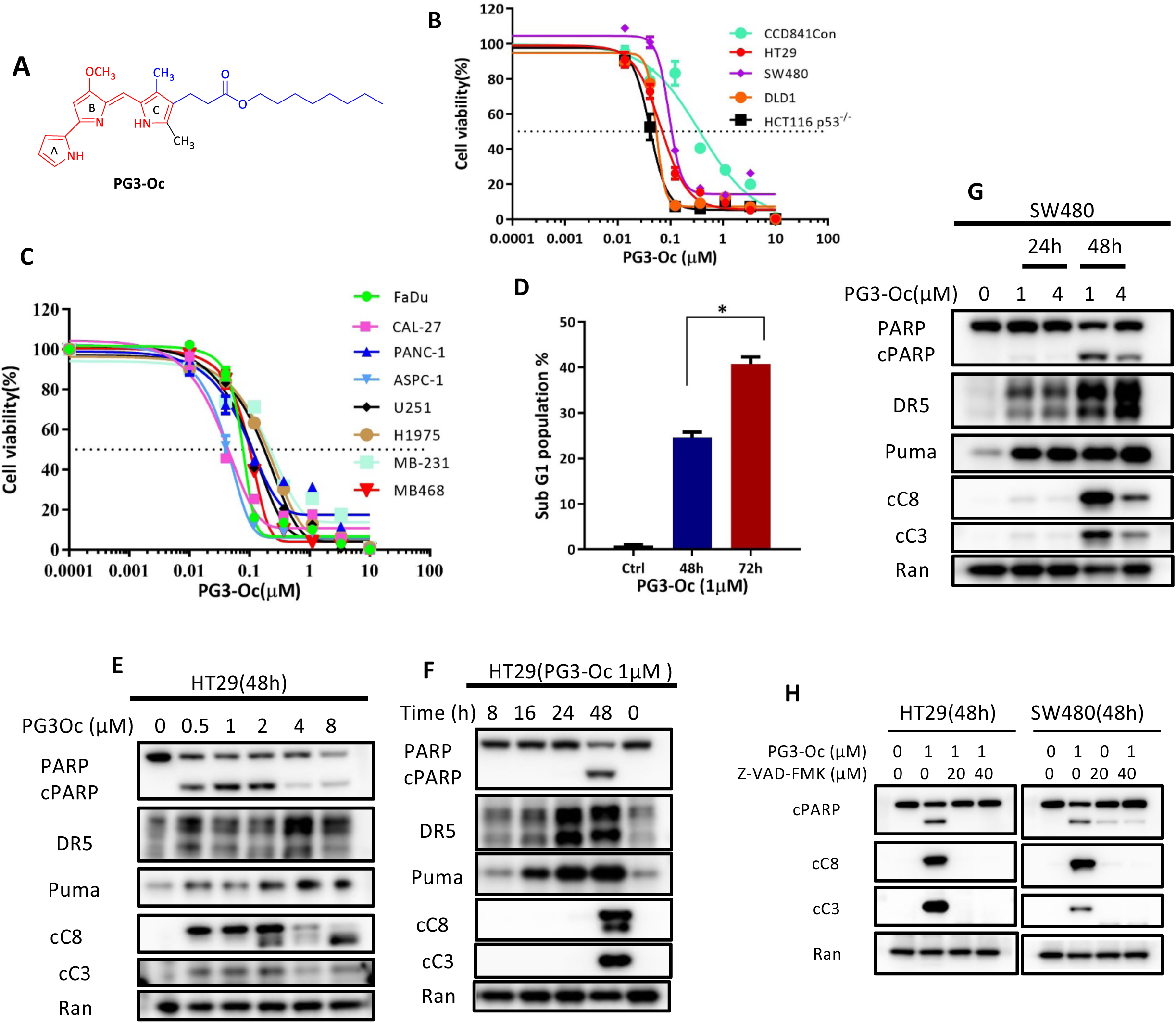
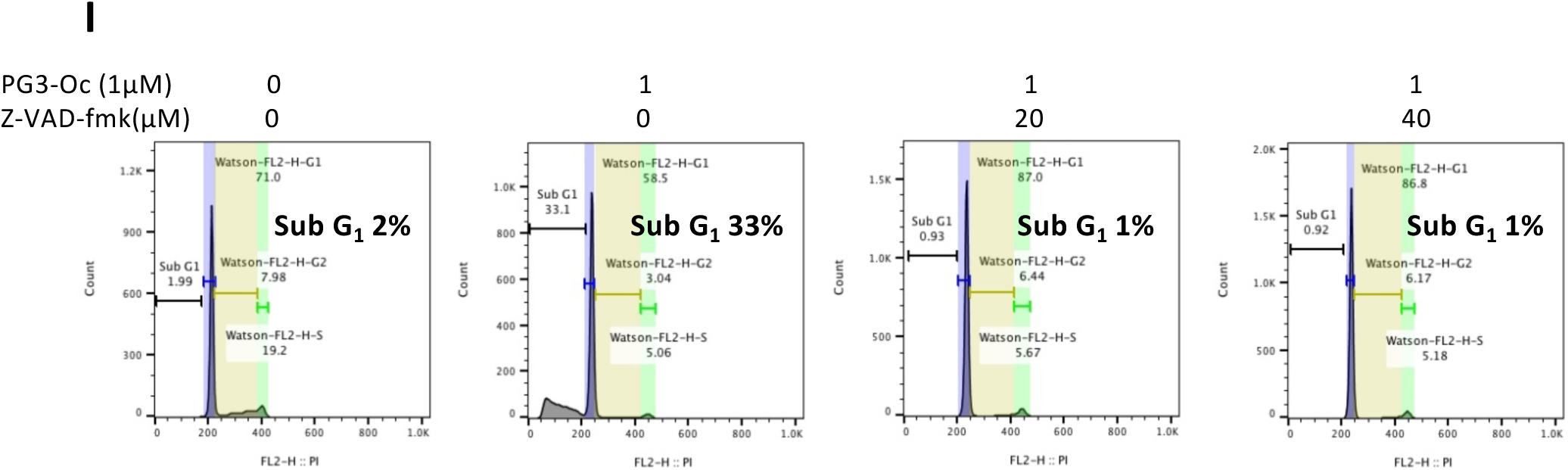
PG3-Oc inhibits cell proliferation and induces apoptosis in mutant p53-expressing cancer cell lines. **(A)** Structure of PG3-Oc. (**B) and (C)** Cell viability assay, dose response curves and IC_50_ value measurement of PG3-Oc in a panel of cancer cell lines. Cells were treated with different concentrations of PG3-Oc (Oc), or DMSO for 72 hr. Luciferase activity was imaged by the IVIS Imaging System after treatment. Cell viability data were normalized to those of DMSO treatment control in each cell line and data analyses were performed using PRISM4 software. IC_50_ data are expressed as the mean ± SD (normal; *n* = 3). (**D)** Cell-cycle profiles after PG3-Oc treatment and apoptosis were analyzed by nuclear PI-staining using flow cytometry. HT29 cells were treated with PG3-Oc at the indicated concentrations for 48 hr or 72 hr respectively. **(E)**, **(F)** and **(G)** Dose-response and time-course analysis of cleavage of caspase-3, −8, −9, cleaved PARP (cPARP), PUMA, and DR5 in PG3-Oc-treated HT29 cells or SW480 cells by western blot using the indicated antibodies. **(H)** Western blot analysis of active caspase-8, active caspase-3 and cleaved PARP in HT29 and SW480 cells. (**I)** HT29 cells were co-treated with 1 μM PG3-Oc and the pan-caspase inhibitor Z-VAD-fmk for 48 hr. Sub G1 populations were analyzed by nuclear PI-staining using flow cytometry.

## Results

### PG3-Oc inhibits cell proliferation and induces apoptosis in mutant p53-expressing cancer cell lines

As a candidate p53-pathway restoring compound with an undefined mechanism of action, PG3-Oc (Figure 1A) is a potent inhibitor of cell proliferation and is efficacious in a broad spectrum of human cancer cells with mutant p53, with IC_50_ values within the nano-molar range (Table 1). PG3-Oc has a 4- to 9-fold therapeutic index in colorectal cancer (CRC) cell lines as compared to normal colon cells CCD 841 Con (Figure 1B and Table 1). In addition, PG3-Oc has anti-proliferative effects on other tumor cell types, including head and neck squamous cell cancer cell lines, pancreatic cancer, breast cancer, glioblastoma multiforme and non-small cell lung cancer (NSCLC) cell lines (Figure 1C and Table 1). Similar to CRC, the IC_50_ in additional tumor types is also in the sub-micromolar range (Table 1). Over 90% inhibition in long-term cell proliferation is also observed in a panel of CRC cell lines treated with low dose PG3-Oc (Figure S1 A and B). Treatment with PG3-Oc induces a 2-fold increase in caspase 3/7 activity as compared to untreated cells using mutant p53-expressing and p73-null cancer cells (Figure S1 C). PG3-Oc’s apoptotic activity is p73-independent as evident by the comparable caspase 3/7 activity in both p53 mutant DLD-1 and DLD1-p73^-/-^ cells post PG3-Oc treatment (Figure S1 C). Treatment of colorectal cancer cell lines HT29 and SW480 with PG3-Oc induces cancer cell death in a dose-and-time dependent manner as demonstrated by sub-G1 analysis (Figure 1D, Figure S1 D and E).

**Table 1.** IC_50_ values for different cancer cell lines with various mutant p53 status.

| **Tumor/Tissue Type** | **Cell Line** | **IC<sub>50</sub>(nM)** | **P53 status** |
| --- | --- | --- | --- |
| **Colorectal Cancer** | HT29 | 66.3 | R273H |
|  | SW480 | 95.3 (4-fold) | R273, P309S |
|  | DLD1 | 54 | S241F |
|  | HCT116 p53<sup>-/-</sup> | 41.1(9 fold) | Null |
| **Non-transformed colorectal epithelial cells** | CCD 841 Con | 375.2 | WT |
| **Head & neck squamous cell carcinoma** | FaDu | 66 | R248L |
|  | CAL-27 | 33.9 | H193L |
| **Pancreatic Cancer** | PANC-1 | 135.5 | R273H |
|  | ASPC-1 | 39.2 | Frameshift |
| **Breast Cancer** | MDA-MB-231 | 242.3 | R280K |
|  | MDA-MB-468 | 97.6 | R273H |
| **Glioblastoma Multiforme** | U251 | 100.2 | R273H |
| **NSCLC** | H1975 | 190.4 | R273H |

To evaluate if the cell death is caspase-dependent, apoptosis markers were analyzed by western blot. As seen in Figure 1E, as low as 0.5 μM PG3-Oc is sufficient to activate cleaved caspase-8 and −3 and cleaved-PARP in HT29 cells. Time-course experiments indicate that PUMA protein is first induced at 16 hours post PG3-Oc treatment and this induction is sustained even at 48 hrs. At 48 hours, we note that induction of cleaved PARP, as well as cleaved caspase-8 and −3 occur in both HT29 and SW480 CRC cell lines (Figure 1F and G). These data also clearly indicate that PG3-Oc induces upregulation of PUMA and DR5 in a dose- and time-dependent manner.

Caspase-dependent induction of apoptosis was further confirmed by the pan-caspase inhibitor (Z-VAD-FMK) co-treatment experiments with PG3-Oc. Western blot analysis show that Z-VAD-FMK completely inhibits the cleavage of caspase-8 and caspase-3 in both HT29 and SW480 cells (Figures 1H). Under the same experimental conditions, 20 μM Z-VAD-FMK completely blocks the formation of a sub-G1 population as compared to the untreated control (Figure 1I). Taken together, these data suggest that PG3-Oc treatment induces caspase-8 and caspase-3 activation in CRC cell lines, and caspase activation is required for PG3-Oc-induced cell death.

### PG3-Oc partially restores global p53 pathway signaling

Having confirmed that PG3-Oc induces apoptosis in multiple mutant p53-expressing cancer cell lines, we investigated whether this small molecule restores the p53 signaling pathway more globally in p53-mutant HT29 and HCT116 p53^-/-^ cells after treatment with 1 µM PG3-Oc for 24 hours. Meta-analysis approaches that enabled comparisons of multiple genome-wide data sets of p53 binding and gene regulation revealed that (1) the transcription factor p53 itself is solely an activator of transcription, (2) gene downregulation by p53 is indirect and requires p21 (Fischer, 2017). Therefore, we focused on upregulated p53 target genes to assess p53 pathway restoration.

Initially, we investigated whether PG3-Oc induces key p53 target genes that regulate cellular apoptosis by qRT-PCR and Western blot. HT29 cells were treated with 1 μM PG3-Oc at different time points followed by qRT-PCR analysis. Time-dependent induction of DR5 (Death Receptor 5), p21 and PUMA (<u>P</u>53-<u>U</u>pregulated <u>M</u>ediator of <u>A</u>poptosis) transcripts is observed in PG3-Oc treated cells (Figure 2 A). Importantly, PG3-Oc very strongly induces upregulation of *PUMA* mRNA in all three cell lines at the 8- or 19-hour time points (Figure 2 A, B and C). Over a 3-fold induction of p21 mRNA is observed at 8 and 19 hours post-treatment with PG3-Oc in HT29 and HCT116 p53^-/-^ cells, but no significant change is observed in SW480 cells. For the DR5 mRNA level, approximately a 2-fold upregulation at 19 hours post-treatment is observed in HT29 and SW480 cells, but not in HCT116-p53^-/-^ cells (Figure 2 A, B and C).

**Figure 2.**
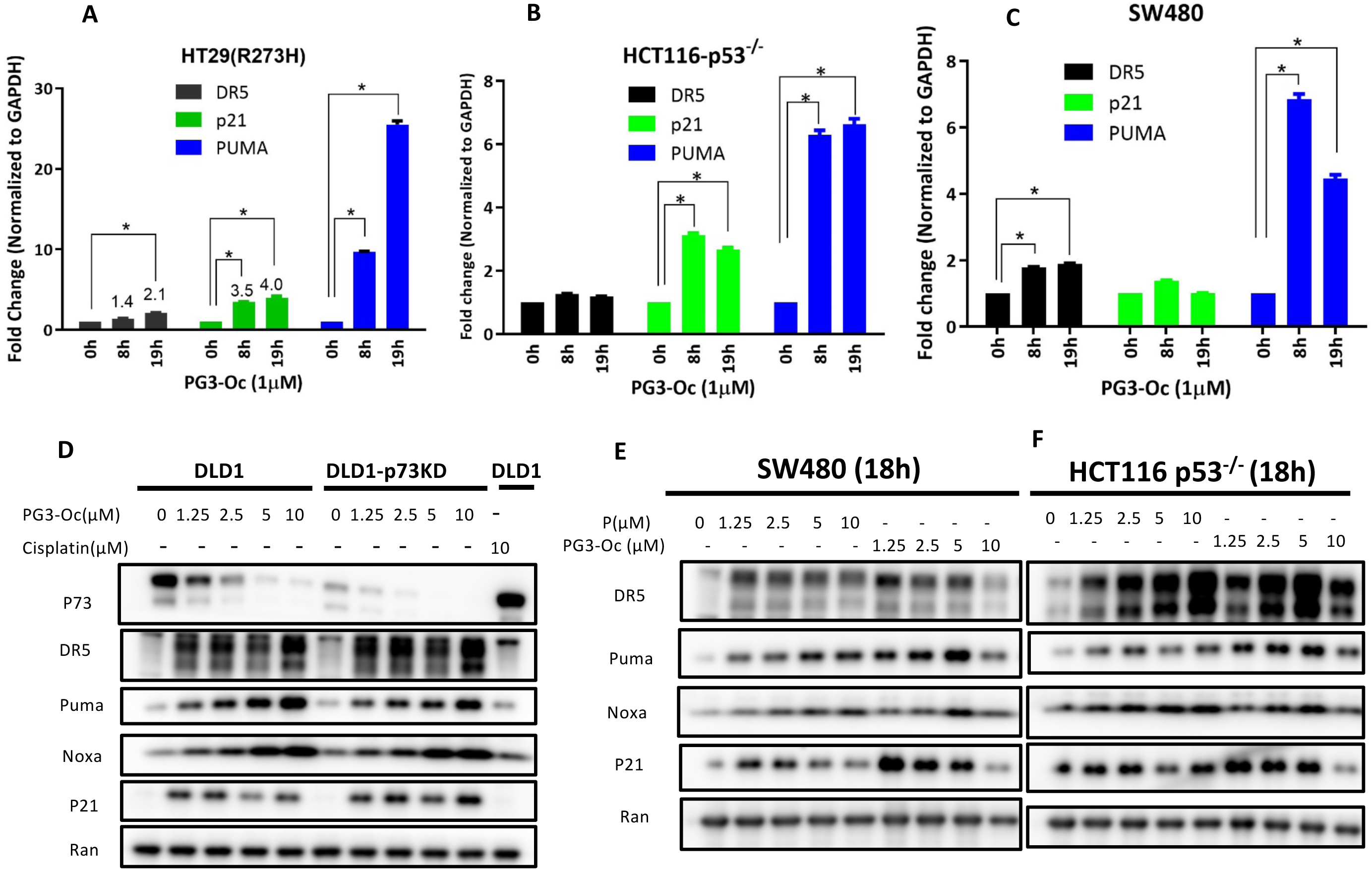
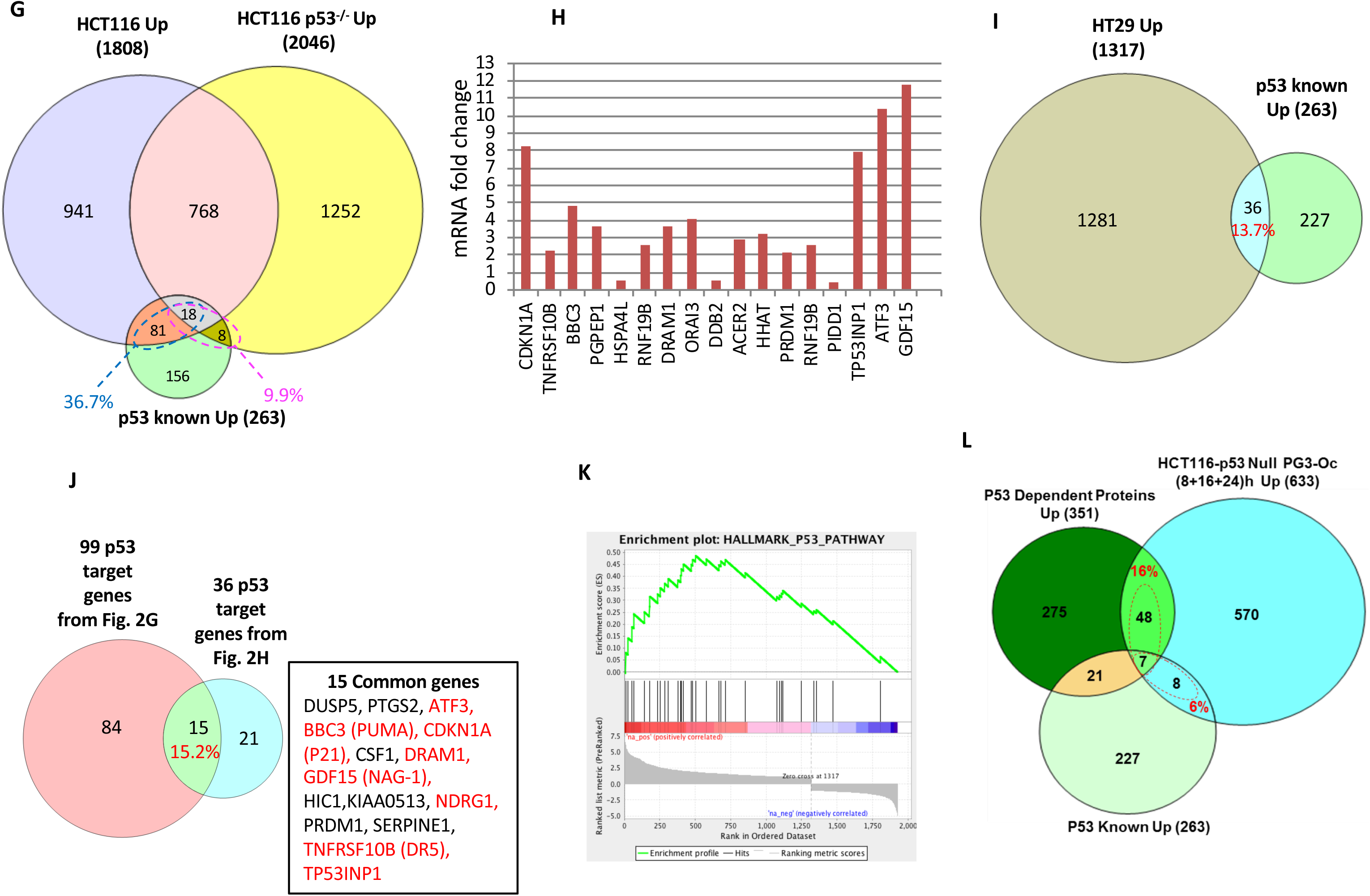
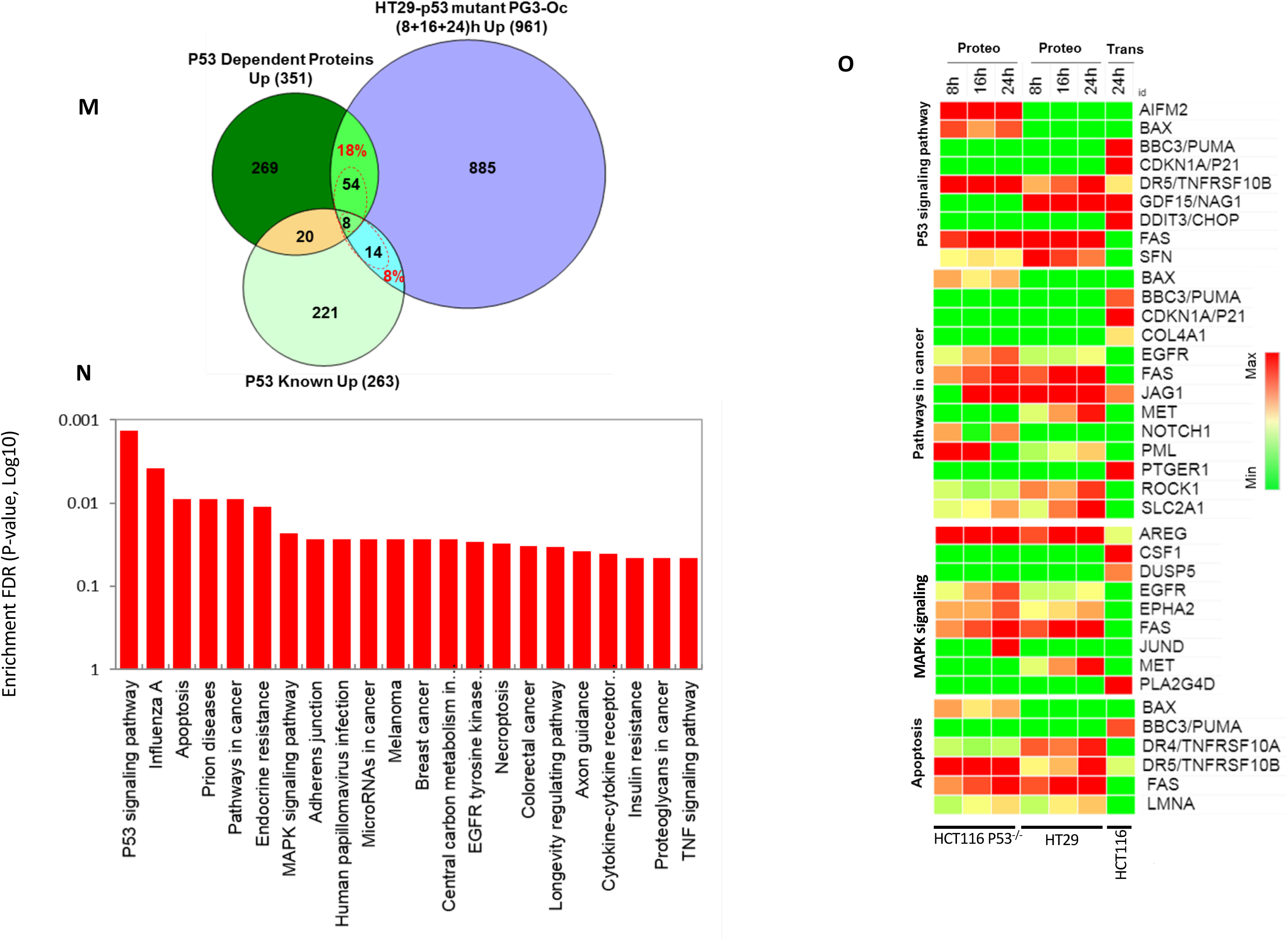
PG3-Oc partially restores the p53 pathway globally. (**A)**, (**B**) and (**C**) Cells were treated with 1 μM PG3-Oc for 8 and 18 hr. qRT-PCR analysis of the change of mRNA level in HT29 and HCT116 p53^-/-^ and SW480 cells. mRNA samples were prepared and RT-PCR was performed to prepare cDNAs as described in Materials and Methods. **(D)**, **(E)** and **(F)** Western blot analysis of p53-target gene expression of DR5, PUMA, Noxa and p21 in p53-mutant and p53-null cancer cells. Cells were treated with the indicated doses of PG3-Oc for 18 hr. **(G)** HCT116 and HCT116 p53^-/-^ cells were treated with/without 50 µM 5-FU for 24 hr in triplicate, and RNA samples were prepared. RNA-seq were performed. Venn diagram shows that 5-FU induced p53 target gene expression. **(H)** A subset of typical p53 target genes were positively regulated by PG3-Oc. (**I**) HT29 cells were treated with or without 1 µM PG3-Oc for 24 hr in triplicate, and RNA samples were prepared. RNA-Seq, IPA and GSEA were performed (see Materials & Methods for details). Venn diagram shows that PG3-Oc restores p53 target gene expression partially. (**J)** Common genes between 99 p53 target genes induced by 5-FU from Figure 2 **G** and 36 p53 target genes induced by PG3-Oc from Figure 2 **H**. (**K**) GSEA plot: Representative gene set from 1867 differential expression genes showing specific responses to the p53 pathway. **(L)** and **(M)**. Proteomic identification of PG3-Oc responsive p53 restored protein in HCT116-p53^-/-^ and HT29 cell lines. Cells were treated with 1 µM PG3-Oc for 8, 16 and 24 hours. Venn diagram analysis were performed in comparison with the in-house build p53 dependent gene (proteome) and known 53 gene data-base. **(N)** KEGG gene enrichment analysis of the overlapped p53 restored proteins identified from proteome analysis (**L** and **M**). As expected the p53 signaling pathway proteins were top hit. In addition, proteins associated with cancer, MAPK signaling, and apoptosis signaling pathway were highly enriched. **(O)** Heat map analysis of p53 restored genes identified by proteomic analysis of HCT116-p53^-/-^ and transcriptome analysis of HT29 in response to PG3-Oc treatment. Proteins were selected from the KEGG gene enrichment pathway analysis in Figure 2 **N**.

Western blot analysis of p53-mutant DLD1, SW480, HT29 cells, and HCT116-p53^-/-^ colon cancer cells show strong upregulation of DR5, p21, PUMA and Noxa (Phorbol-12-Myristate-13-Acetate-Induced Protein 1) in a time- and dose-dependent manner (Figure 1E, F and G; Figure 2 D, E and F), which is consistent with the qRT-PCR results.

In order to develop a reference p53 target/responsive gene collection as a control, HCT116 cells and HCT116 p53^-/-^ cells were treated with 50 µM of the p53 activator 5-Fluorouracil (5-FU) for 24 and 48 hours. Western blot indicates that a set of typical p53 target genes (p21, PUMA, DR5, NAG-1 and Noxa) is significantly upregulated over time in HCT116 cells, whereas their expression is not changed in HCT116 p53^-/-^ (Figure S2 A). Then, isogenic HCT116 and HCT116 p53^-/-^ cells were treated with or without 50 µM 5-FU for 24 hours in triplicate, RNA samples were prepared and RNA-seq was performed (Supplemental Files S1-S3). Principal component analysis (PCA) shows close clustering of total normalized mRNA abundance of the replication in each condition, however, each treatment condition is significantly distinct from other groups (Figure S2 B and C). The global change in transcription across the groups compared is visualized by a volcano plot (Figure S2 D and E). Ingenuity pathway analysis (IPA) revealed that the p53 pathway is a top hit and is activated in HCT116 cells, but not in HCT116 p53^-/-^ cells (Figure S2 F and G). GSEA (gene set enrichment analysis) show that p53 signaling pathway is a top hit and the most enriched pathway in HCT116 cells, but not in HCT116 p53^-/-^ cells (Figures S2 H, I and J). Not surprisingly, the “p53 pathway” is a top hit found in both IPA and GSEA analyses.

Fischer’s p53 target gene set contains a total of 344 including 263 genes positively regulated by p53 and 81 genes negatively regulated by p53 (Fischer, 2017). The total of 344 and 263 genes is found in a spreadsheet in supplementary data (Supplementary Table S1 in the Fischer manuscript) rather than the total of 346 and 246 genes referred to in the text of the manuscript (Fischer, 2017). As Figure 2G shows, compared to known p53 target gene data base (263 positively regulated genes only), 5-FU induced upregulation of 99 (81+18) p53 target genes in p53 wild-type HCT116 (cutoff of fold change = 1.87, FDR 0.05), which covers 37.6% (99/263) of the p53 target gene database. Though p53 is a *bona fide* transcription factor of p53 target genes, this data clearly indicates that 5-FU through DNA damage partially restores the p53 pathway in p53 wild-type HCT116 cells. By contrast, in HCT116 p53^-/-^ cells, 26 p53 target genes were upregulated, which is p53-independent. This represents 9.9% (26/263) restoration of p53 target genes through unknown transcription factors in p53-deficient cells treated with 5-FU. These results are consistent with the notion that transcription programs by p53 are dependent on cell-type, as well as the nature of the stresses and inducers.

HT29 cells were treated with or without 1 µM PG3-Oc for 24 hours. RNA samples were prepared, and then RNA-Seq was performed (Supplemental Files S4). Key p53 target genes *CDKN1A* (p21), *TNFRSF10B* (DR5), *BBC3* (PUMA), *TP53INP1* (Teap) and *GDF15* (NAG-1) are significantly upregulated (Figure 2 H) and identified in IPA canonical p53 pathway analysis (Figure S2 K and L). IPA analysis (cutoff of log2 fold change = 2, FDR 0.05) of 1317 up-regulated genes revealed that among the 263 known p53 target genes, 36 genes are up-regulated (Figure 2I). That is 13.7% (36/263) of 263 total p53 genes, and higher than 9.9% of 5-FU-induced restoration of the p53 target genes in HCT116 p53^-/-^ cells.

There are 15 overlapping p53 target genes between the 99 genes induced by 5-FU from Figure 2G and the 36 genes induced by PG3-Oc from Figure 2I as shown in a Venn diagram analysis (Figure 2J), which covers 15.2% (15/99) of 99 5-FU-induced p53 target genes (Figure 2J). Importantly, the analysis indicates that critical effector p53 target genes that are involved in cell cycle arrest (p21), apoptosis (PUMA, DR5, Noxa and NAG-1), autophagy (DRAM1), and suppression of cancer cell metastasis (NDRG1) are in this overlapping gene set and are potently induced (Figure 2J, highlighted in red). These data suggest that although the percentage of restoration of p53 target genes is dependent on both cell type and nature of inducers, the p53 core gene set that regulates cell proliferation and apoptosis could be induced regardless of cell type and properties of inducers in tested cell lines.

GSEA analysis indicates that PG3-Oc-induced differential expression of genes is enriched in the p53 pathway (Figure 2 K), suggesting PG3-Oc has a significant impact on upregulation of p53 target genes in p53-mutant cells. Also, the apoptosis signaling pathway is highly enriched in PG3-Oc treated tumor cells (Figure S2 M).

Because very limited information is available on the proteomic changes in p53 mutant and p53-null cell lines, we investigated the proteomic response of HCT116 and HCT116 p53^-/-^ cell lines treated with or without 50 µM 5-FU for 24 hours (Figure S3 A to E). In a comparative analysis, a total of 448 proteins are increased in abundance at least 1.5-fold in response to 5-FU in the HCT116 cell line whereas 455 proteins are increased in the HCT116 p53^-/-^ cell line. A comparison of these two proteome sets showed 283 proteins are unique in the HCT116 cell line whereas 165 proteins are overlapping with the p53-null cell line (Figure S3 F). Enrichment analysis indicates that p53-regulated metabolic signaling was significantly enriched in HCT116 cells, but not in HCT116 p53^-/-^ cells (Figure S3 G and H). Among these overlapping proteins, 68 show increased abundance of at least 1.2-fold in HCT116 cells as compared with the p53-null cell line. Thus, a total of 351 (283 + 68) proteins are considered as our p53-responsive proteome dataset (Figure 2L) which we further use as a positive p53 control proteome for comparison with PG3-Oc treatment of the HT29 and HCT116-p53^-/-^ cell lines (Supplementary Table 2).

To evaluate the restoration of p53-dependent proteins in HCT116 p53^-/-^ and mutant p53-expressing HT29 cell lines in response to PG3-Oc treatment, the cell lines were exposed to 1 µM PG3-Oc treatment for 8, 16 and 24 hours and subjected to proteomic analysis, respectively (Figure S3 I to M). Because expression of p53 target genes is also time-dependent, genes upregulated at any of the three timepoints (8, 16 and 24 hours) were included. A total of 633 (Figure 2 L) and 961 (Figure 2 M) proteins were increased at least 1.5-fold in response to PG3-Oc treatment in HCT116 p53^-/-^ and HT29 cells, respectively (Figure 2 L, M and Supplementary Tables 3 and 4). These protein sets were further compared with our reference p53-responsive proteome dataset (Figure 2 L, M and Supplementary Table 2). Venn diagram analysis shows that 16% (55/351) of the p53-responsive proteins are restored in the HCT116-p53 ^-/-^ cell line in response to 1 µM PG3-Oc treatment as compared with our reference p53-responsive proteome dataset. An additional 6% (15/263) of upregulated proteins are overlapping with the known p53 target gene database (Figure 2L). Similarly, 18% (62/351) proteins are restored in the mutant p53-expressing HT29 cell line when compared with our reference p53-responsive proteome dataset and an additional 8% (22/263) are overlapping with the known p53 target gene dataset (Figure 2M). This analysis suggests that proteomic analysis of p53 pathway restoration is feasible and the percentages of p53 target genes restored by PG3-Oc in the HT29 cell line are similar: 13.7% in the transcriptome and 18% in the proteome analysis (Figure 2 I and M).

KEGG gene enrichment analysis of the overlapping p53 restored proteins identified from Figure 2 L and M proteome analysis found that the p53 signaling pathway is a top hit (Figure 2 N), suggesting PG3-Oc is able to restore proteins of p53 target genes partially, which is consistent with the transcriptome analysis.

We performed a heat map analysis of restored p53 pathway genes identified by proteomic analysis of HCT116 p53^-/-^ and HT29 cell lines in response to PG3-Oc treatment (Figure 2 O). Proteins were selected from the KEGG gene enrichment pathway analysis in Figure 2 N (Supplementary Table 6). Proteins associated with cancer, MAPK signaling, and apoptosis signaling pathways are highly enriched (Figure 2 O).

Taken together, these data suggest that PG3-Oc can partially restore the p53 pathway in mutant p53-expressing HT29 and p53-null HCT116 cancer cell lines at both the transcriptional and protein levels.

### PUMA is required for PG3-Oc mediated cell death

PUMA is a BH-3-only Bcl-2 family member that binds and inactivates anti-apoptotic proteins like Bcl-2, Bcl-X_L_, and Mcl-1. This facilitates induction of the caspase-9 mediated intrinsic apoptosis pathway (Carneiro and El-Deiry, 2020; Lomonosova and Chinnadurai, 2008). DR5 activation results in recruitment of the adaptor protein FADD (Fas-associated death domain) and caspase-8 to form the DISC (death inducing signaling complex), leading to the cleavage and activation of caspase-8. Activated caspase-8 can directly activate effector caspase-3/7 *via* the extrinsic pathway (Type I) or cleave Bid (BH3 interacting-domain death agonist) and activate the intrinsic pathway (Type II), leading to cell apoptosis (Carneiro and El-Deiry, 2020; LeBlanc and Ashkenazi, 2003). Since PUMA and DR5 are important proapoptotic proteins, we evaluated if PUMA and DR5 are dispensable for PG3-Oc mediated cell death in mutant p53-expressing cells. As shown in Figure 3A, when PUMA was knocked down, alone or together with DR5, using siRNA, there was complete blunting of PARP cleavage and cleavage of caspases after PG3-Oc treatment. Similar results were observed when knockdown of PUMA by siRNA reduced the sub-G1 population to 11.1% as compared to 25.8% in siControl, in PG3-Oc treated cells (Figure 3B and Figure S4 A). DR5 knockdown alone had no impact on the same apoptotic markers and Sub-G1 population under the experimental conditions (Figure 3 A, B and Figure S4 A). However, our preliminary data shows that PG3-Oc-induced upregulation of DR5 sensitizes TRAIL-resistant HT29 cells to TRAIL treatment (Figure S4 C, D and E).

**Figure 3.**
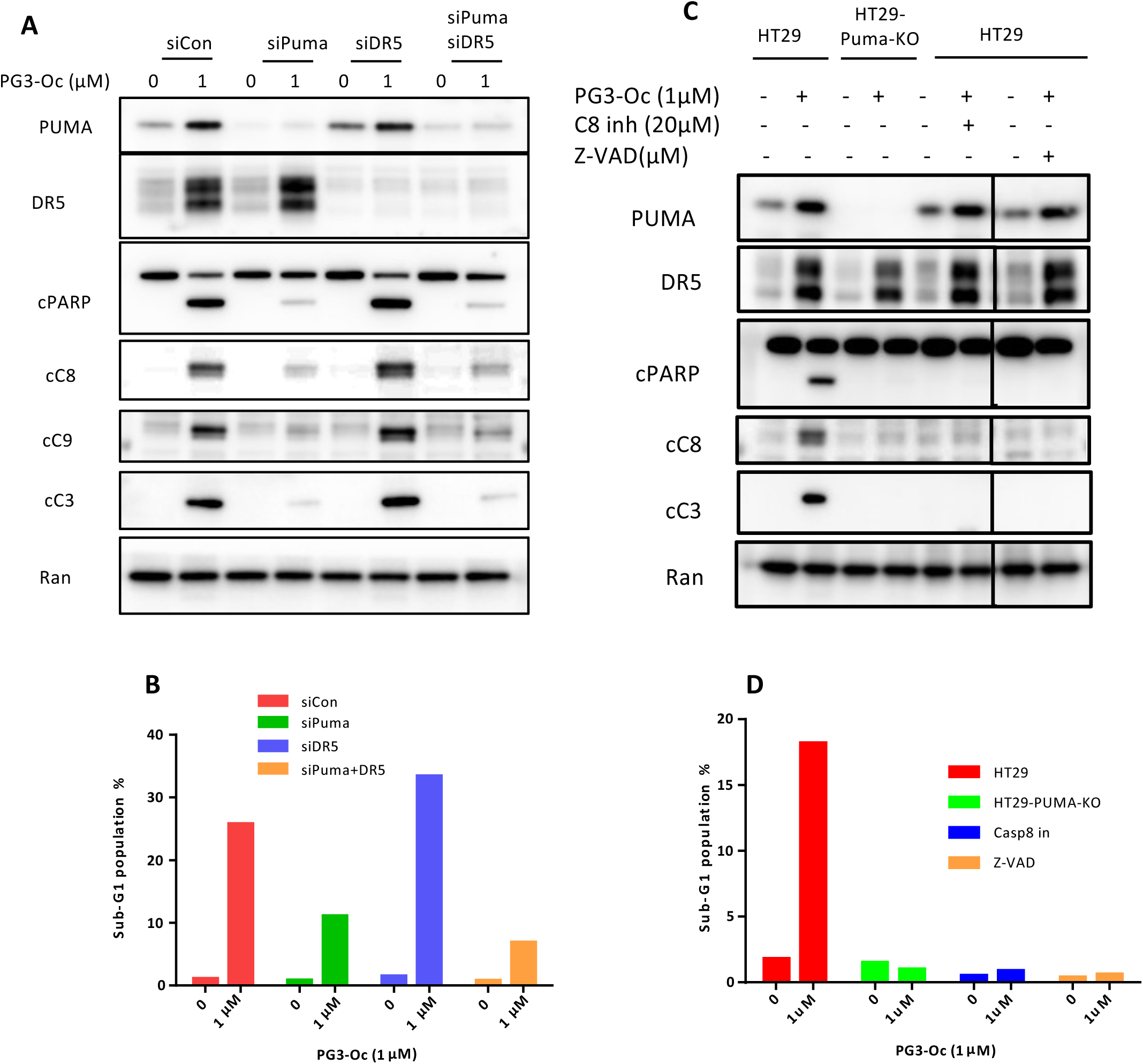
PUMA is required for PG3-Oc-mediated cell death. (**A)** HT29 cells were transfected with Control, PUMA, DR5 and PUMA/DR5 siRNAs, and at 24 hr after transfection, the cells were treated with 1 μM PG3-Oc for 48 h. After treatment, western blot analysis of PUMA, DR5, cleavage of caspase-8, −3, −9 and cleaved PARP was performed using indicated antibodies. (**B)** Cell death was analyzed by nuclear PI-staining using flow cytometry. **(C)** HT29 and HT29-PUMA-KO cells were treated with PG3-Oc or co-treated with the caspase 8 inhibitor (cas8 inh) or pan-caspase (Z-VAD-FMK) inhibitor for 48 hours. Cleavage of caspases and PARP were detected by western blotting using the indicated antibodies. (**D**) Sub G1 populations were analyzed by flow cytometry.

PUMA siRNA studies were validated by creating *PUMA* gene knockout HT29 cells *via* CRISPR/Cas9 gene-editing technology (Figure S5, for details see Materials and Methods). The gRNA was designed to target the DNA sequence that encodes amino-acid residues for the BH3-domain of PUMA (Figure S5 A). Knockout of the *PUMA* gene abolishes PG3-Oc-induced cleavage of PARP and caspase-8, −3 and the sub-G1 population was the same as the positive control caspase-8 inhibitor Z-IETD-fmk and the pan-caspase inhibitor Z-VAD (Figure 3C and 3D and Figure S4 B). Taken together, these data suggest PUMA protein is required and is a key mediator in cell death induced by PG3-Oc treatment in HT29 cancer cells.

Of note, both knockdown and knockout of the *PUMA* gene abolishes caspase-8 and caspase-3 cleavage/activation and PARP cleavage after PG3-Oc treatment (Figure 3A and 3C). Furthermore, the caspase-8 inhibitor Z-IETD-fmk not only inhibits caspase-8 cleavage, but also results in inhibition of caspase-3 and PARP cleavage. These data suggest that the induced PUMA is able to feedback to mediate the activation of caspase-8 through an unknown mechanism.

### PG3-Oc dependent repression of MYC upregulates PUMA

PG3-Oc-induces significant downregulation of MYC and upregulation of PUMA protein levels is observed in a panel of p53 mutant cell lines, such as HT29, DLD1, FaDu, MDA-MB-231, MDA-MB-468, SW480 and CAL27 (Figure 4A). Inhibition of MYC leads to downregulation of expression of its target gene E2F1, and inhibition of expression of E2F1 target gene p73 (Figure 4B). Transcriptome data analysis in HT29 cells also shows that PG3-Oc treatment downregulates E2F1 and p73 mRNA levels as compared to untreated control cells (Figure S 2K and 2L). Experiments in isogenic HCT116 cells with wild-type p53 or p53^-/-^ cells show no significant differences in induction of PUMA or downregulation of MYC by PG3-Oc (Figure 4C). Taken together, these data suggest that PG3-Oc-induced downregulation of MYC and upregulation of PUMA is not limited to a specific cell line or p53 mutation, and is independent of p53 status.

**Figure 4.**
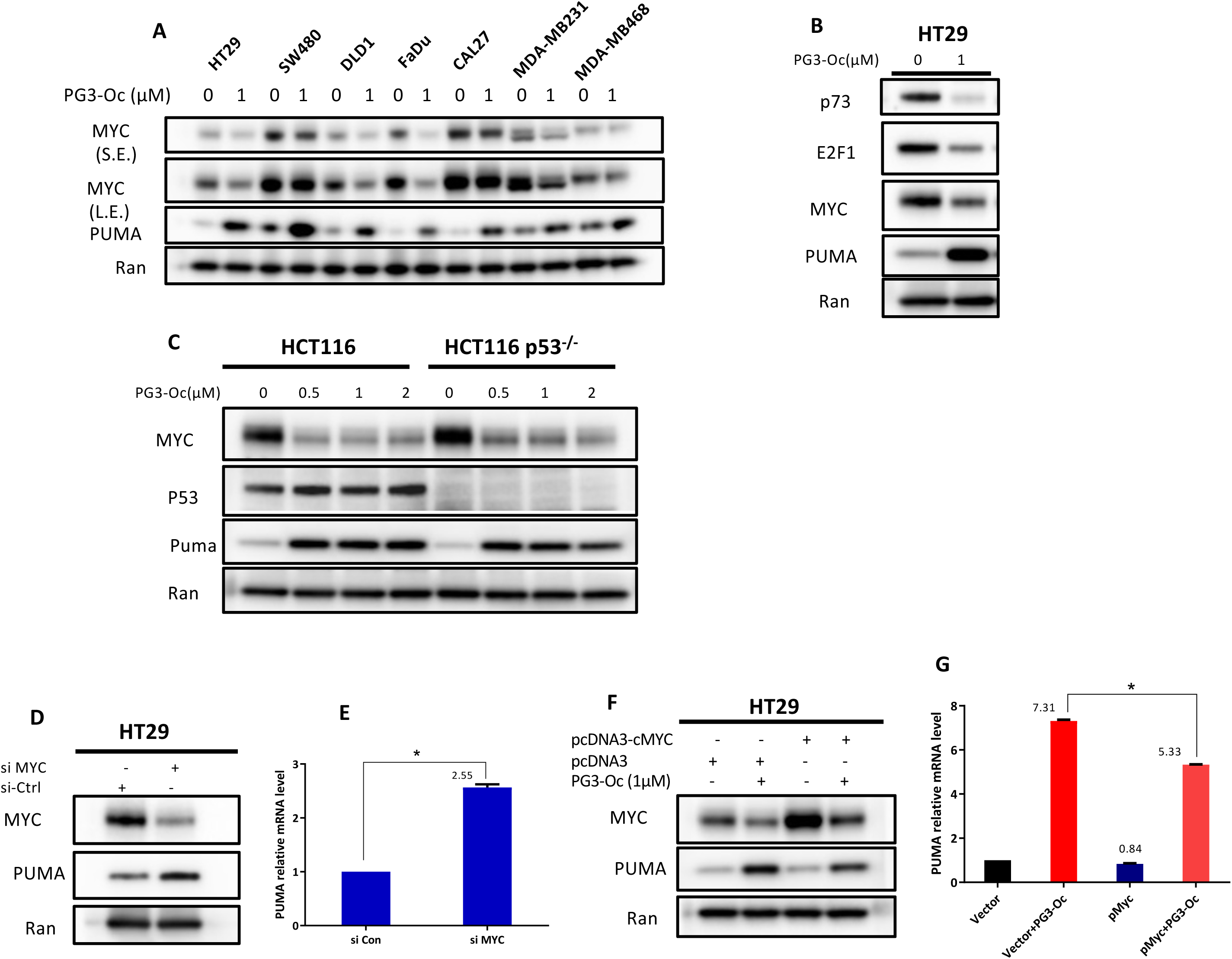
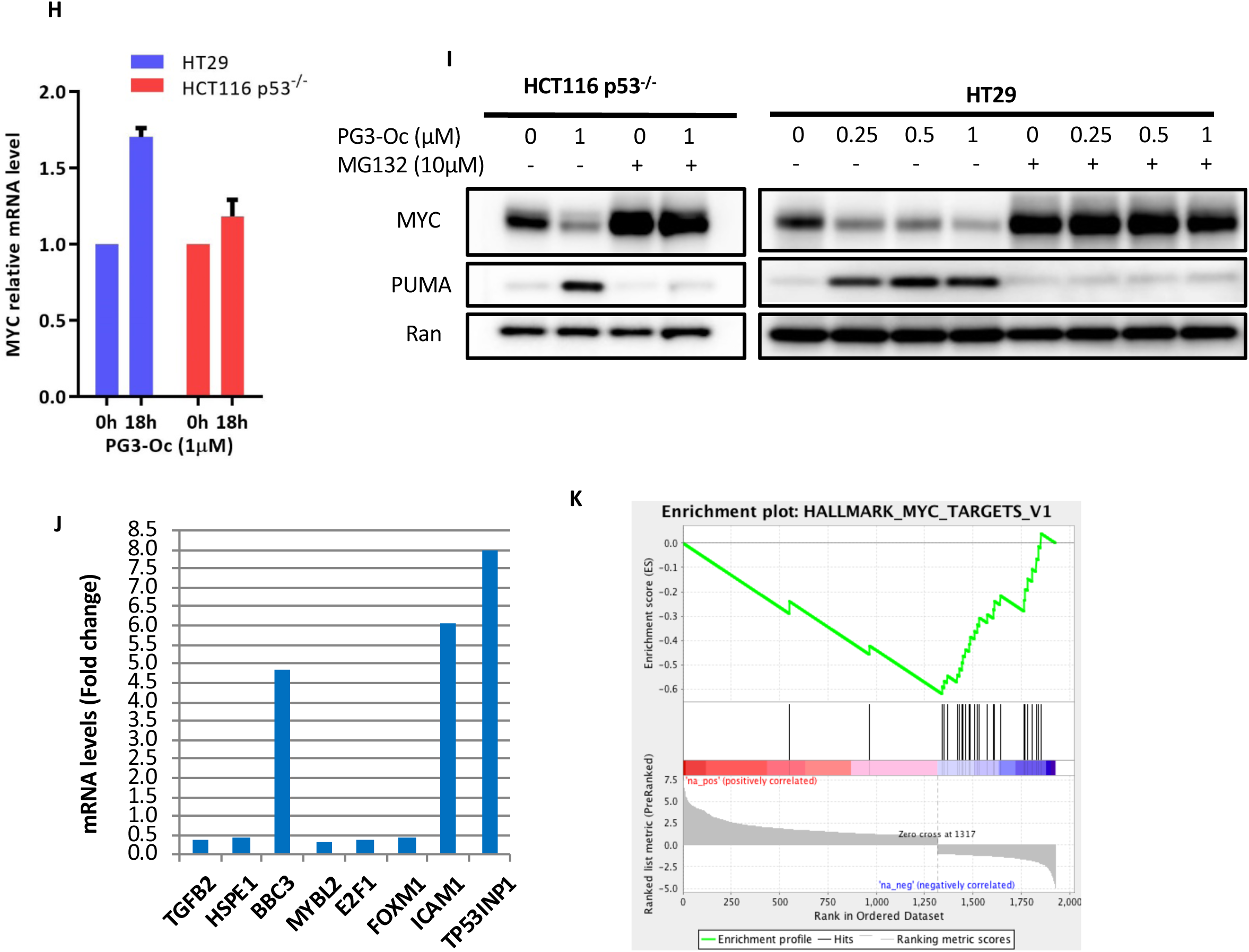
PG3-Oc dependent repression of MYC upregulates PUMA. **(A)** Various mutant Mutant p53-expressing cancer cell lines were treated with PG3-Oc for 24 hours. Cell lysates were prepared for western blots using the indicated antibodies. S.E.: short exposure; L.E.: long exposure. **(B)** HT29 cells were treated with 1 μM PG3-Oc for 24h, and western blots were performed using the indicated antibodies. (**C)** HCT116 and HCT116 p53^-/-^ cells were treated with PG3-Oc at the indicated doses for 24 hr, and western blots were carried out by using the indicated antibodies. (**D)** Cells were transfected with MYC siRNAs and control siRNAs for 48 hr, cell lysates were prepared for western blots, and (**E**) mRNAs was extracted for qRT-PCR analysis to evaluate the change of PUMA mRNA levels after PG3-Oc treatment. **(F)** HT29 cells were transfected with the vector pcDNA3 and pcDNA3-cMYC, and after 24h, cells were treated with 1 μM PG3-Oc for 17 hr. Cell lysates were prepared for western blots. (**G**) mRNA was extracted for qRT-PCR analysis to determine the change of PUMA mRNA levels after PG3-Oc treatment. **(H)** HT29 and HCT115 p53^-/-^ cells were treated with PG3-Oc for 18 hr, and mRNAs was extracted for cDNA synthesis and qRT-PCR analysis to determine the change of MYC mRNA levels after PG3-Oc treatment. **(I)** Cells were co-treated with the proteasome inhibitor MG132 for 24 hr, and western blot was performed with the indicated antibodies. **(J)** Analysis of MYC target genes from RNA-Seq analysis with differential gene expression induced by 1 µM PG3-Oc after 24 hr treatment in HT29 cells. **(K)** GSEA plot: Representative gene set from 1867 differential expression genes with specific response to MYC.

It was reported that MYC binds to the *PUMA* promotor and represses *PUMA* gene transcription (Amente et al., 2011; Fernandez et al., 2003). We explored whether PG3-Oc-induced downregulation of MYC activates PUMA gene transcription. Basal PUMA levels are modestly de-repressed on knockdown of MYC, both at the protein and mRNA levels (Figure 4D and 4E). Over-expression of MYC leads to attenuation of PUMA induction at both the protein and mRNA levels (Figure 4F and 4G) post PG3-Oc treatment. qRT-PCR data indicates that PG3-Oc treatment does not significantly change MYC mRNA levels in either HT29 or HCT116 p53^-/-^cell lines (Figure 4H), suggesting MYC downregulation may be occurring through the proteasome pathway. To study whether endogenous MYC can inhibit PG3-Oc-induced upregulation of PUMA or not, HCT116 p53^-/-^ and HT29 cells were co-treated with PG3-Oc and proteasome inhibitor MG132. MG132 blocks MYC degradation and leads to accumulation of endogenous MYC that abolishes the PG3-Oc-induced upregulation of PUMA (Figure 4I). Taken together, these data indicate that degradation of MYC allows PUMA gene expression, which is consistent with MYC negative regulation of PUMA expression (Amente et al., 2011; Yun et al., 2016). MYC protein degradation occurs through the proteasome pathway in the tested cell lines.

RNA-Seq data of PG3-Oc-treated HT29 cells indicates that MYC-activated genes (Figure 4J), *TGFB2* (TGFβ-2), *HSPE1* (Hsp10), *MYBL2* (B-Myb), *E2F1* (E2F1) and *FOXM1* (FOXM1) (Fernandez et al., 2003), are downregulated. By contrast, MYC-repressed genes, *BBC3* (PUMA), *ICAM1* (ICAM1) and *TP53INP1* (TP53INP1) (Amente et al., 2011; Yun et al., 2016), are upregulated (Figure 4J). GSEA analysis indicates that downregulation of genes is enriched in the MYC pathway, suggesting that PG3-Oc has a significant negative impact on the MYC pathway and network (Figure 4K).

In summary, our data demonstrate that PG3-Oc-induced degradation of MYC correlates well with subsequent upregulation of PUMA in the tested cell lines.

### ATF4 is a key regulator that mediates PG3-Oc-induced p53 pathway restoration

We sought to discover upstream regulators that mediate p53 pathway restoration in mutant p53-expressing cancer cells following PG3-Oc treatment. We also questioned which transcription factor may positively regulate *PUMA* gene expression after MYC degradation. Transcription factors p73 and p63 are p53 family members, and the majority of the genome-wide p53 target sites can be bound by p63 and p73 *in vivo* (Smeenk et al., 2008). p73 or p63 bind to p53-responsive elements and regulate PUMA gene expression in response to various stressful stimuli (Melino et al., 2004; Pyati et al., 2011). However, in this case, we observed that PG3-Oc treatment leads to downregulation of p73 both at the protein level (Figure 2D and Figure 4B) and the mRNA level (Figure S2 K and L). Consistent with this, induced upregulation of DR5, p21, PUMA and Noxa between DLD1 and DLD1-p73^-/-^ cell lines show no significant differences (Figure 2D). To further verify these observations, siRNA knockdown of p73 and/or p63 was performed, and was found to not attenuate PG3-Oc-induced upregulation of PUMA in HT29 cells (Figure S6 A). Thus, the upregulation of the p53 pathway by PG3-Oc is independent of p73 and p63.

Transcription factors FOXO3a, NF-κB, and JNK/c-Jun can regulate *PUMA* gene expression in a p53-independent manner depending on cell type and stimuli (Chen et al., 2014; Dudgeon et al., 2012; Dudgeon et al., 2010; Gao et al., 2010; Ghosh et al., 2012; Qing et al., 2012; Zhao et al., 2012). Knockdown of transcription factors FOXO3a and NF-κB (p65) respectively, or inhibition of JNK/c-Jun signaling by JNK inhibitor SP600125 does not blunt PG3-Oc-induced upregulation of PUMA (Figure S6 B and C). These data suggest that NF- κB, FOXO3a, and JNK/c-Jun are not involved in the regulation of PUMA in PG3-Oc-treated cells.

It was reported that PUMA, Noxa, p21 and NAG-1 are ATF4 direct target genes (Inoue et al., 2017; Qing et al., 2012; Wang et al., 2015). We found that knockdown of ATF4 not only blocks PG3-Oc induced upregulation of PUMA, but also DR5, p21, Noxa and NAG-1, as shown in Figure 5 A and B. Also, knockdown of ATF4 does not block PG3-Oc-induced downregulation of MYC, indicating after MYC degradation and de-repression of *PUMA* gene, it is ATF4 that mediates the expression of the *PUMA* gene (Figure 5 A and B). CHOP is an ATF4 direct target gene, and it directly regulates DR5, Noxa and PUMA gene expression (Ghosh et al., 2012; Hu et al., 2018; Teske et al., 2013). We observed that knockdown of ATF4 leads to downregulation of CHOP (Figure 5 A and B), and knockdown of CHOP blocks PG3-Oc-induced upregulation of DR5 and Noxa, but not PUMA, p21 and NAG-1 (Figure 5 C and D). Taken together, these data suggest that ATF4 is an upstream regulator that mediates p53 pathway restoration induced by PG3-Oc in p53-deficient cells.

**Figure 5.**
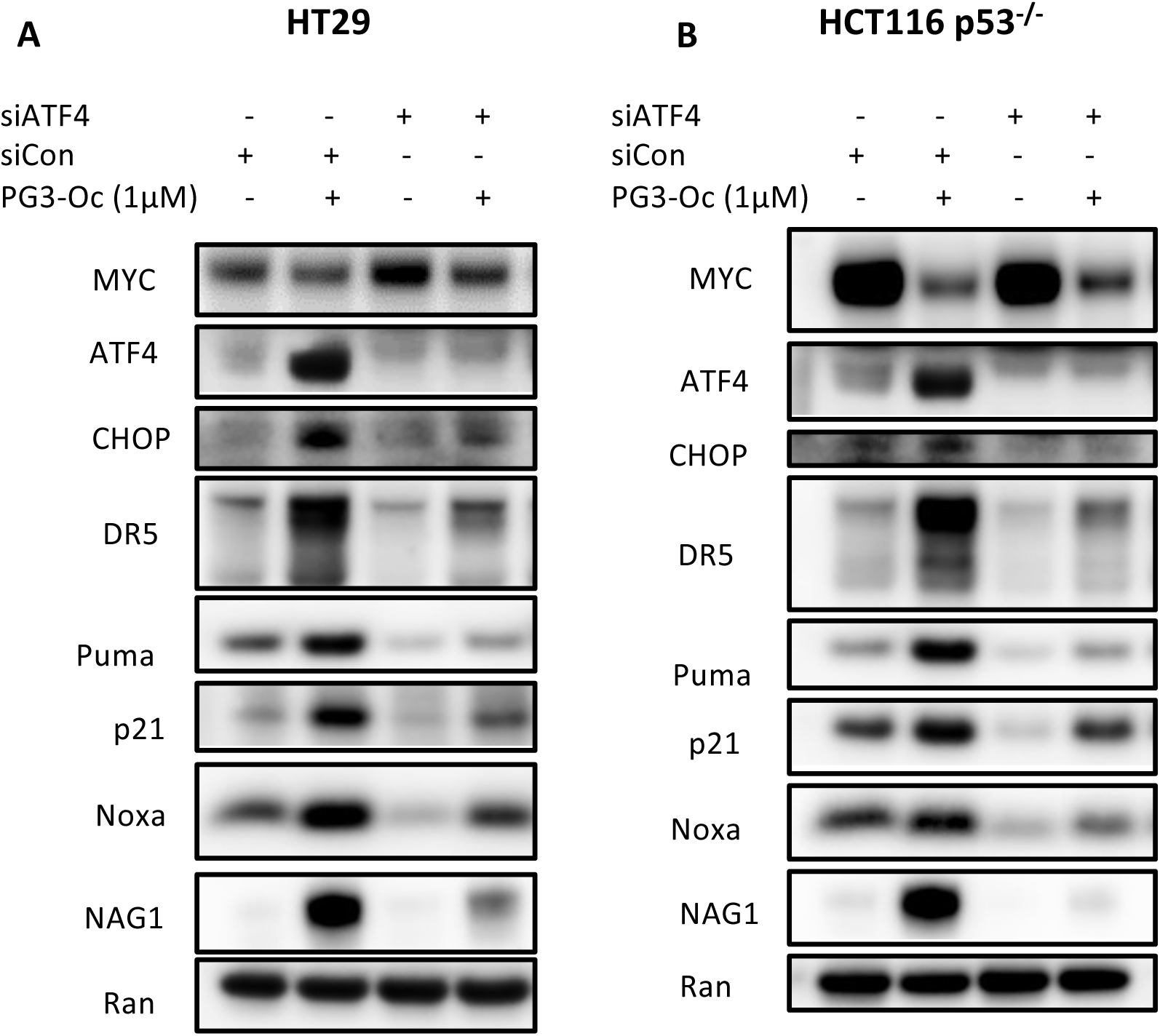
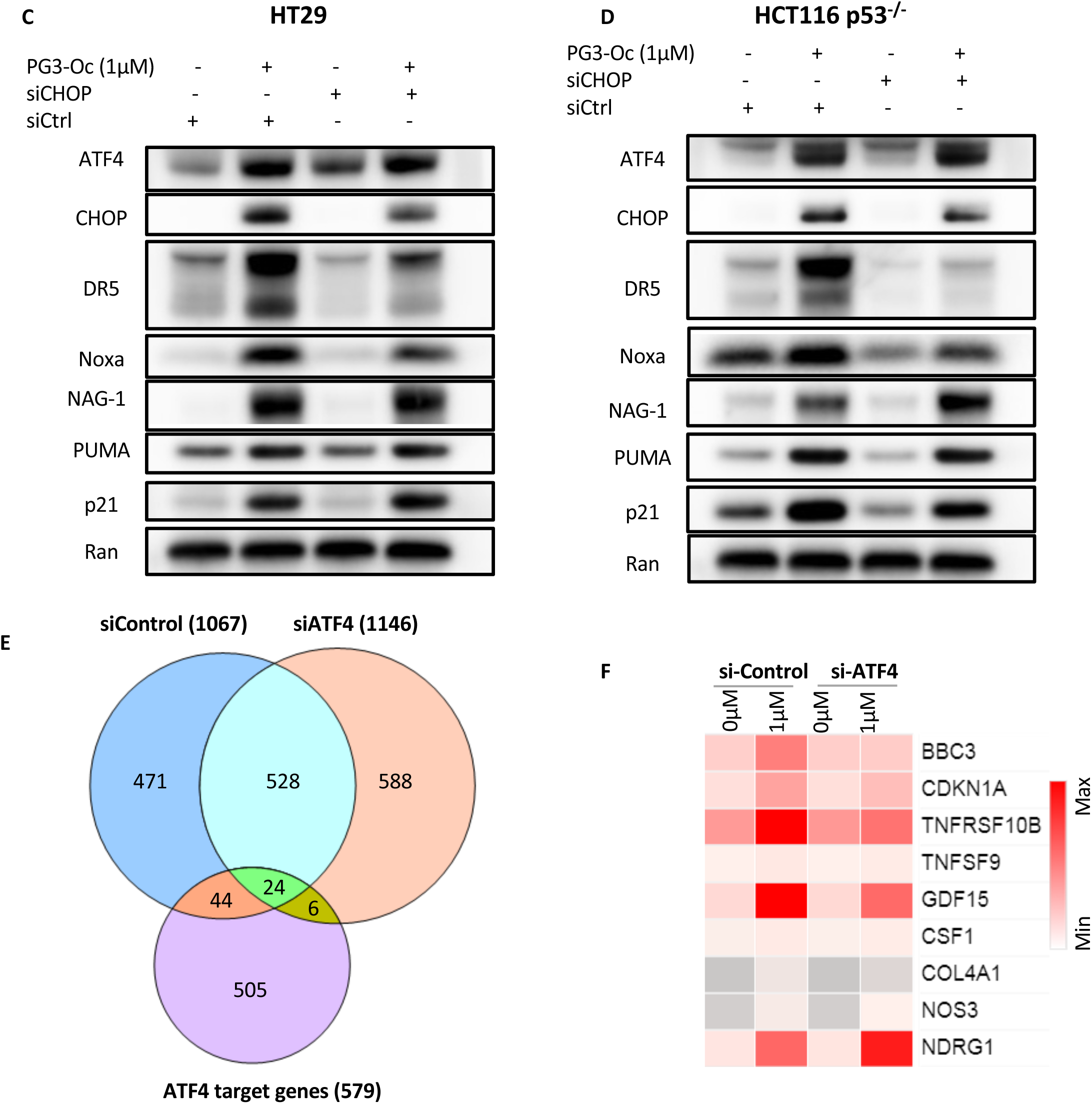
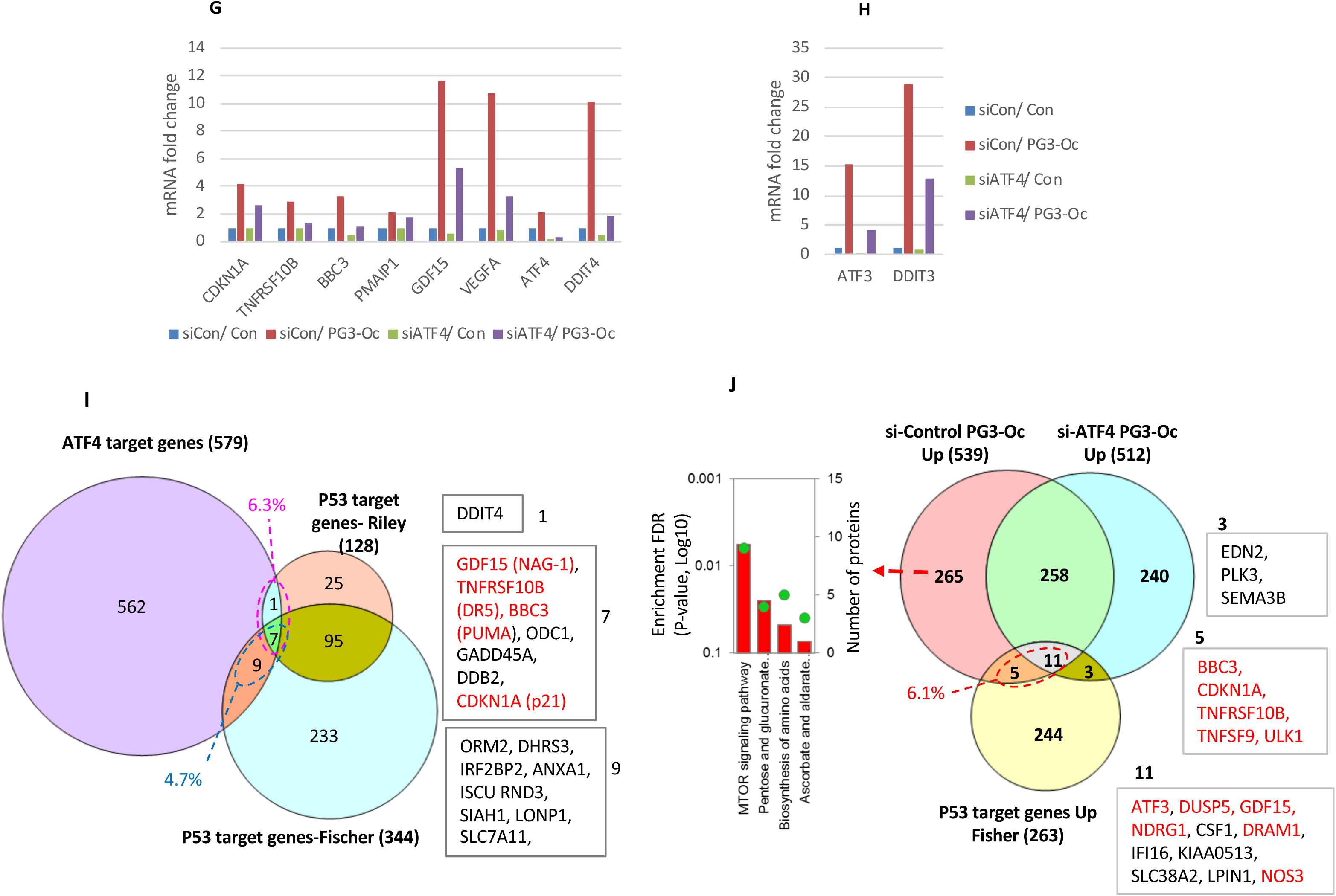
ATF4 is a mediator of PG3-Oc-induced p53 pathway restoration. HT29 **(A)** and HCT116 p53^-/-^ **(B)** cells were transfected with Control and ATF4 siRNAs, and at 24hr after transfection, the cells were treated with 1 µM PG3-Oc for 24h. Then cell lysates were prepared and western blot analysis were performed using indicated antibodies. HT29 **(C)** and HCT116 p53^-/-^ **(D)** cells were transfected with Control and CHOP siRNAs, and at 24hr after transfection, the cells were treated with 1 µM PG3-Oc for 24h. Then cell lysates were prepared and western blot analysis were performed using indicated antibodies. **(E)** HT29 cells were transfected with Control and ATF4 siRNAs, and at 24hr after transfection, cells were treated with/without 1 µM PG3-Oc for 24 hr in triplicate, and RNA samples were prepared. RNA-Seq, and IPA analysis were performed (see Materials & Methods for details). Venn diagram shows that PG3-Oc induces ATF4 target gene expression. **(F)** Heat map and bar charts. (**G** and **H**) Analysis of some key p53 genes expression associated in apoptosis. **(I)** Common genes between known ATF4 target gene database and known p53 target gene data bases (Fischer’s and Riley’s databases). **(J)** Venn diagram analysis showed unique and overlapped genes identified in PG3-Oc treated si-Control and si-ATF4 cell lines in comparison with known p53 gene data-base.

To further confirm our hypothesis, transcriptome analysis of siATF4-knockdown experiment was performed. HT29 cells were transfected with control or ATF4 siRNAs respectively, after 24 hours, cells were treated with or without PG3-Oc for 24 hours. RNA samples were prepared and RNA-Seq and IPA analysis was performed (cutoff of fold change = 2, FDR 0.05) (Supplemental Files S5-S7). IPA canonical pathway analysis identified that the p53 signaling pathway is significantly activated in HT29 cells, but not in siATF4/PG3-Oc-treated HT29 cells (Figure S7 A, B and C). The heatmap of signaling pathway and network analysis clearly indicate that p53 signaling is activated in siControl/PG3-Oc-treated HT29 cells, and inhibited in siATF4/PG3-Oc-treated HT29 cells (Figure S7 D).

YW3-56 is a compound that induces ER stress in the triple negative breast cancer cell line MDA-MB-231. CHIP-exo assay was performed to identify genome-wide ATF4-bindign sites after YW3-56 treatment, and 579 possible ATF4 target genes were identified (Wang et al., 2015). Using this ATF4 target gene data base, Venn diagram (Figure 5E) was generated. 68 (44+24) ATF4 target genes are induced in control siRNA cells by PG3-Oc treatment. After knockdown of ATF4, the number decreased to 24 ATF4 target genes (Figure 5E and Supplementary table 7). Six new ATF4 target genes are induced after ATF4 knockdown, suggesting PG3-Oc also induces upregulation of ATF4 target genes in an ATF4-independent way (Figure 5E). Among the 24 genes, RNA-seq data clearly show the mRNA levels of some genes are significantly decreased due to ATF4 knockdown (Figure 5 F, G and H), such as GDF15 (NAG-1), VEGFA, CDKN1A (p21), DDIT4 (REDD-1), DDIT3 (CHOP) and ATF3. However, the fold changes of these genes remain higher than 2-fold, therefore IPA software still considers them as induced and includes them in the gene list. Based on these observations, the six genes are included as affected by ATF4 knockdown, so a total of 26 (20+6) genes are significantly downregulated, that is a total of 39.4% (26/66) of the ATF4 target genes induced by PG3-Oc are downregulated by ATF4 knockdown.

To find out the percentage of ATF4 target genes that are p53 target genes, a Venn diagram was generated using the ATF4 target gene base and two p53 target gene databases (Figure 5 I and Supplementary table 8)(Fischer, 2017; Riley et al., 2008; Wang et al., 2015). 16 common genes between ATF4 and Fischer’s p53 gene data are found, that is 4.7% (16/344). 8 common genes between ATF4 and Riley’s p53 gene database are found, that is 6.3% (8/128). The common gene set (Figure 5 I box7) contains important target genes that regulate cell apoptosis as indicated in red.

PG3-Oc induces upregulation of 16 p53 target genes, which is 6.1% (16/263) of the 263 p53 target genes. Among the 16 p53 target genes, after knockdown of ATF4, 5 genes are significantly downregulated (Figure 5J box 5 and Supplementary table 5). ATF3 and GDF15 (Figure 5 G and H) are also significantly downregulated, however their mRNA levels are still higher than 2-fold, therefore the two genes are shown in common block 11 (Figure 5J box11). Based on these observations, a total of 7 (5+2) genes among the 16 genes are significantly downregulated after ATF4 knockdown, which indicates that 43.8% (7/16) p53 target genes induced by PG3-Oc are affected by knockdown of ATF4. A set of 265 genes is unique in siControl-PG3-Oc-treated HT29 cells, and KEEG gene enrichment analysis indicates mTOR signaling pathway and biosynthesis of amino acids are significantly affected, which is consistent with ATF4 activation (Figure 5J and Figure S7 E).

In summary, ATF4 regulates expression of a sub-set of key p53 target genes, which are involved in the regulation of the cell cycle and apoptosis.

### PG3-Oc-induced upregulation of ATF4 is not through canonical ER stress

ER stress or the integrated stress response (ISR) activate the UPR (unfolded protein response) signaling pathway. Canonical ER stress activates ATF4 through the PERK/eIF2α/ATF4 pathway, and ISR activates ATF4 through the PKR (or GCN2 or HRI)/eIF2α/ATF4 pathway. Both pathways result in upregulation of ATF4 protein at a post-transcriptional level. To determine which pathway mediates PG3-Oc-induced upregulation of ATF4, time-course studies were performed (Figure 6A and 6B). Time-dependent induction of phosphorylation of Ser51 of eIF2α, upregulation of ATF4, and its target genes CHOP and PUMA is observed in both HT29 and HCT116 p53^-/-^ cell lines (Figure 6A and 6B). Another ER stress marker GRP78 is induced in a time-dependent way in HT29 cells, but not in HCT116 p53^-/-^ cells (Figure 6A and 6B). Thapsigargin (TG) is a known ER stress inducer and used as a positive control. GSK2606414 is a selective and potent inhibitor of PERK. As shown in Figure 6C, HT29 cells were treated for 5 hours. The PERK inhibitor potently inhibits TG-induced phosphorylation of Ser51 of eIF2α, upregulation of ATF4, and its target gene CHOP, but has no effect on PG3-Oc-induced phosphorylation of eIF2α, upregulation of ATF4, and CHOP. These data clearly indicate that PG3-Oc-induced upregulation of ATF4 is not through canonical ER stress. This observation was furthered confirmed by 24 hour treatment of HT29 cells (Figure 7B). The PERK inhibitor does not attenuate PG3-Oc-induced upregulation of ATF4 protein and upregulation of ATF4 downstream gene expression of CHOP, PUMA, p21, NAG-1, DR5 and Noxa. In addition, the PERK inhibitor also does not block phosphorylation of ERK2, suggesting that PG3-Oc-induced upregulation and activation of ATF4 is also not through non-canonical ER stress PERK/ERK2/ATF4 pathway as reported (Ojha et al., 2019).

**Figure 6.**
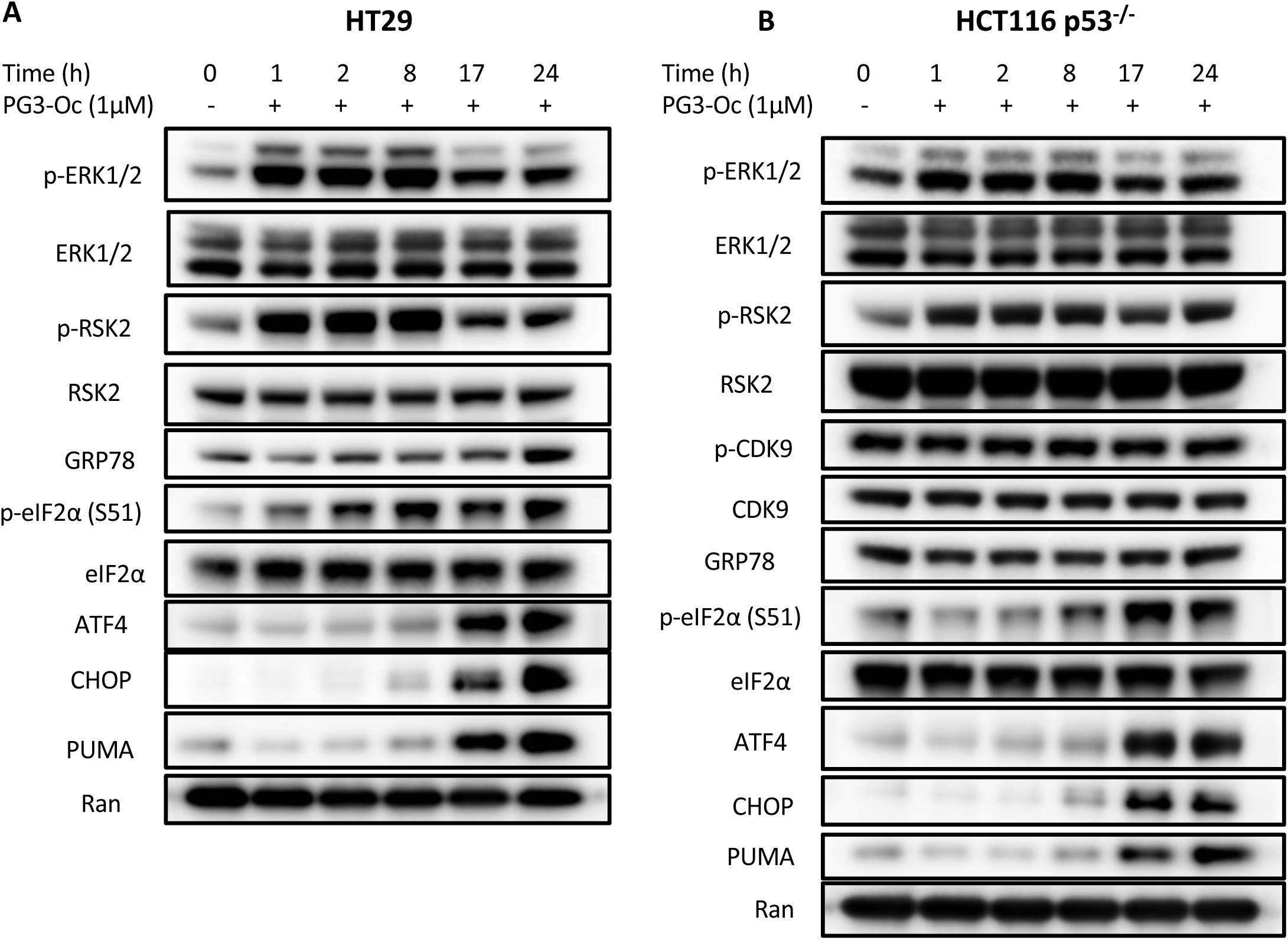
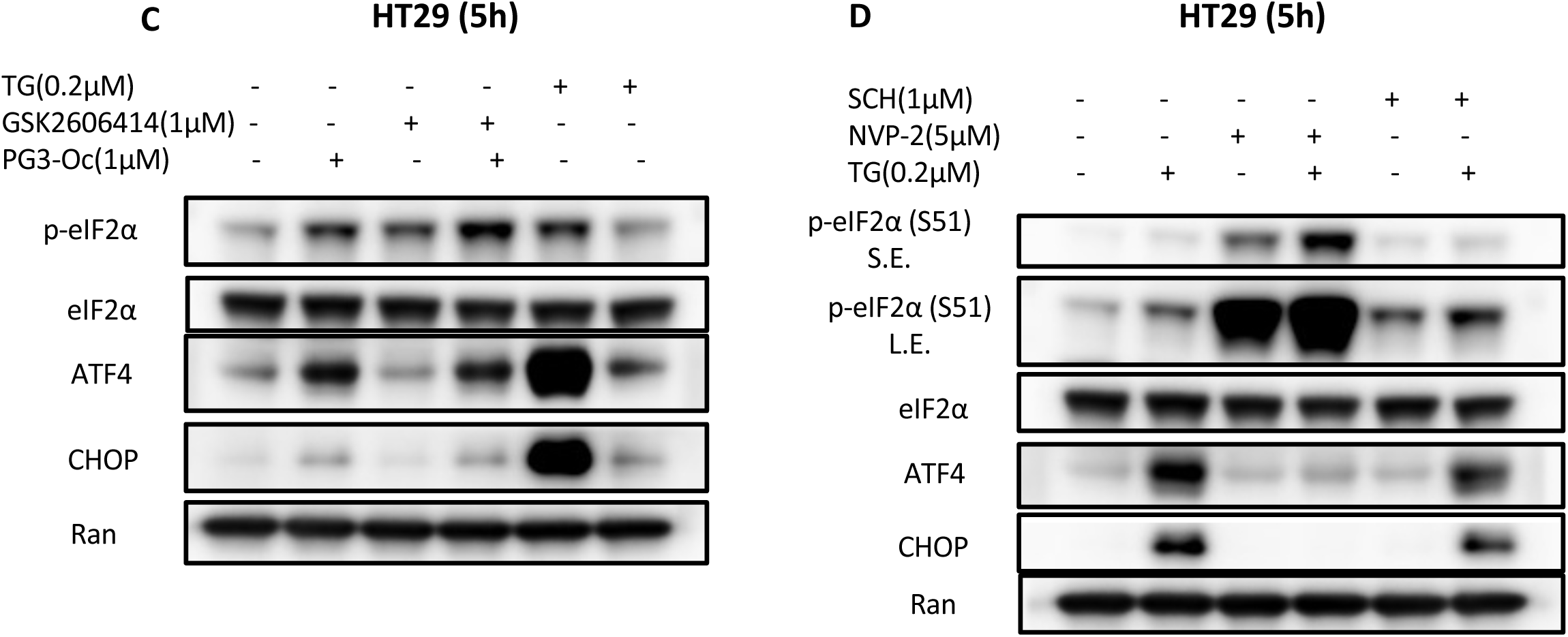
PG3-Oc-induced upregulation of ATF4 is not through ER stress. HT29 **(A)** and HCT116 p53^-/-^ **(B)** cells were treated with PG3-Oc at different time points as indicated in the Figure **A** and **B**. Cell lysate were prepared and western blots were performed using indicated antibodies. **(C)** HT29 cells were treated with GSK2606414, TG and PG3-Oc for 5 hr, and then cell lysates were prepared and western blots were performed using indicated antibodies. **(D)** HT29 cells were treated with thapsigargin (TG), SCH772984 (SCH) and NVP-2, respectively or combined treatments. Cell lysates were prepared and western blots were performed using indicated antibodies.

**Figure 7.**
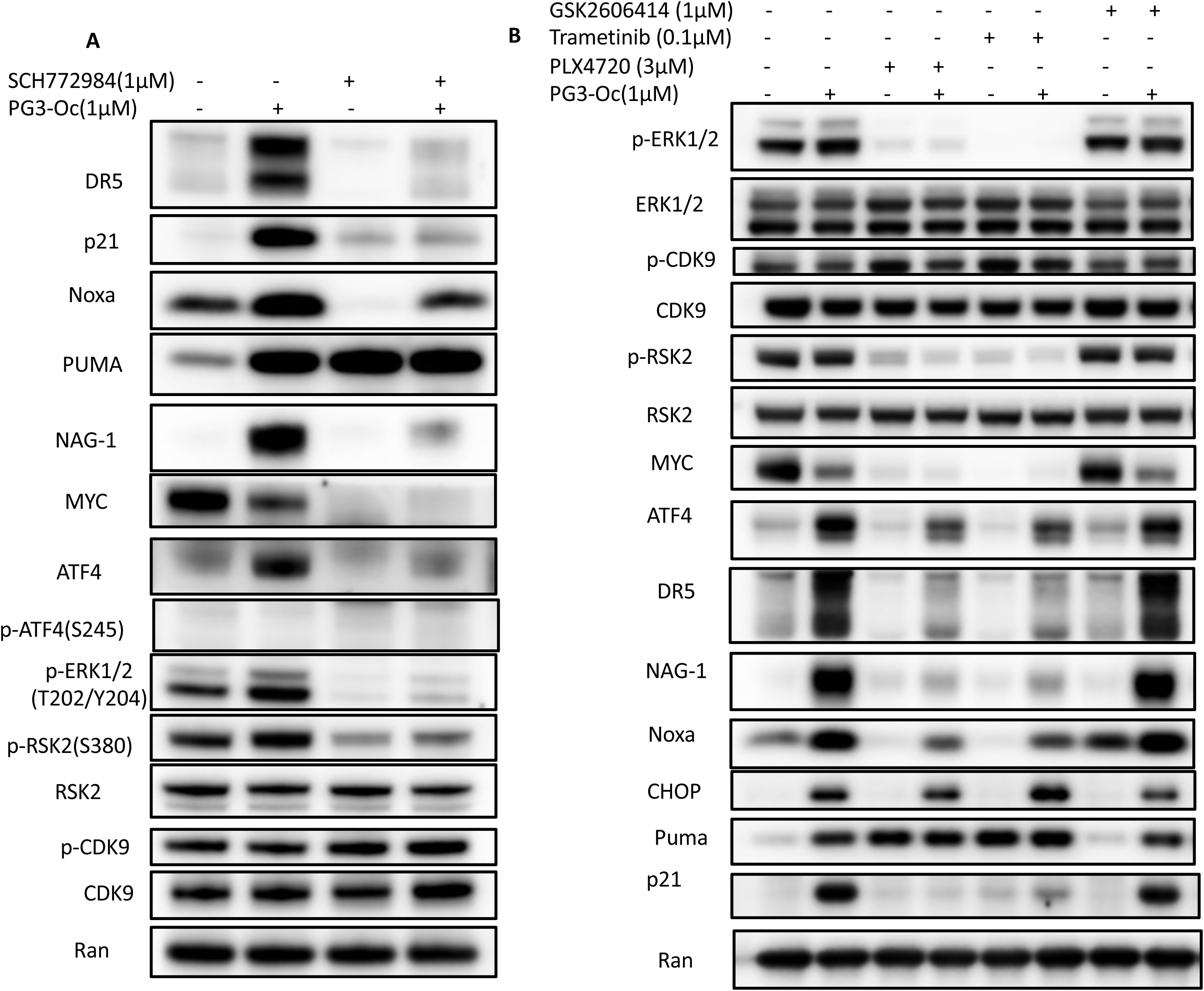
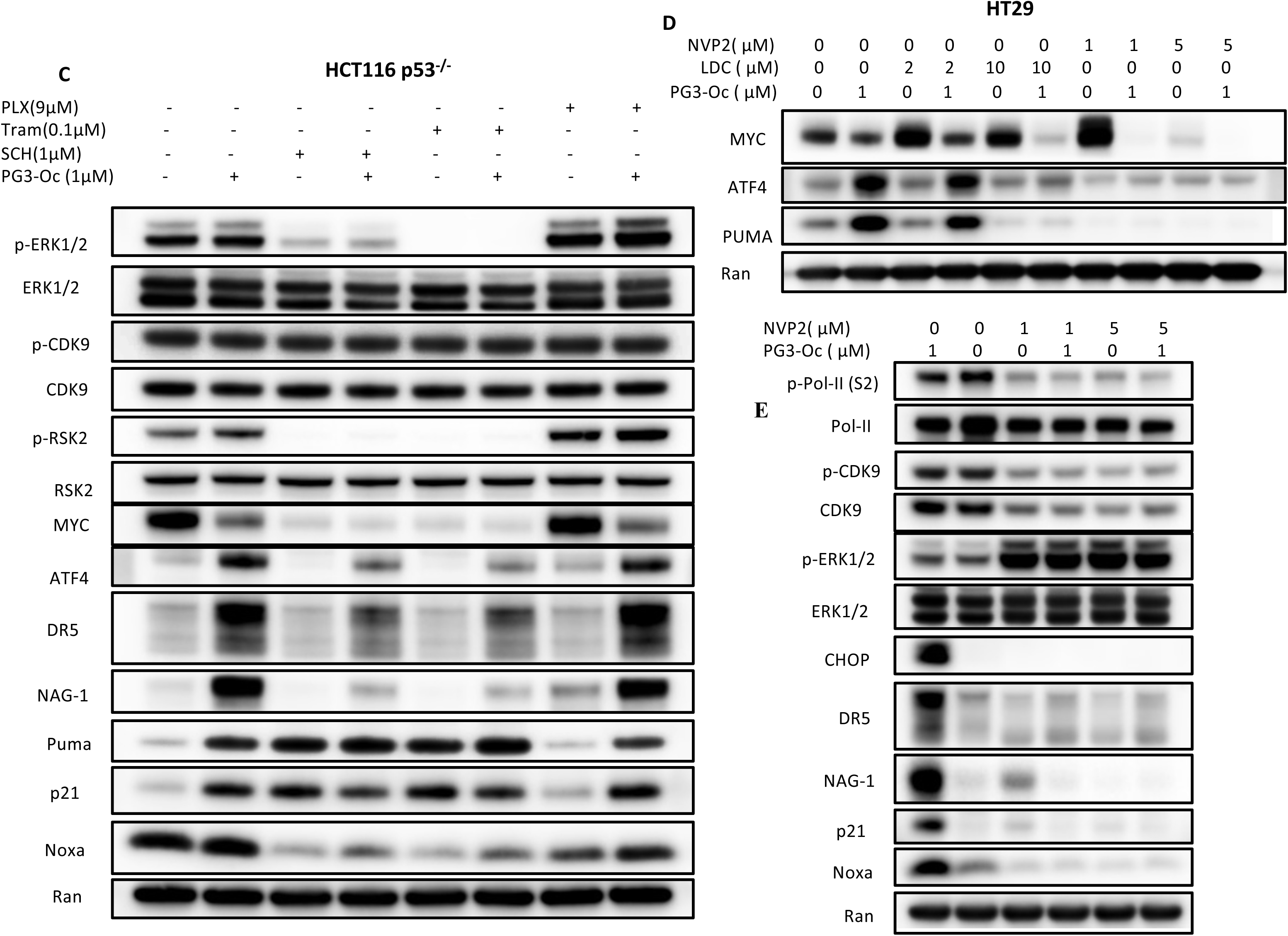
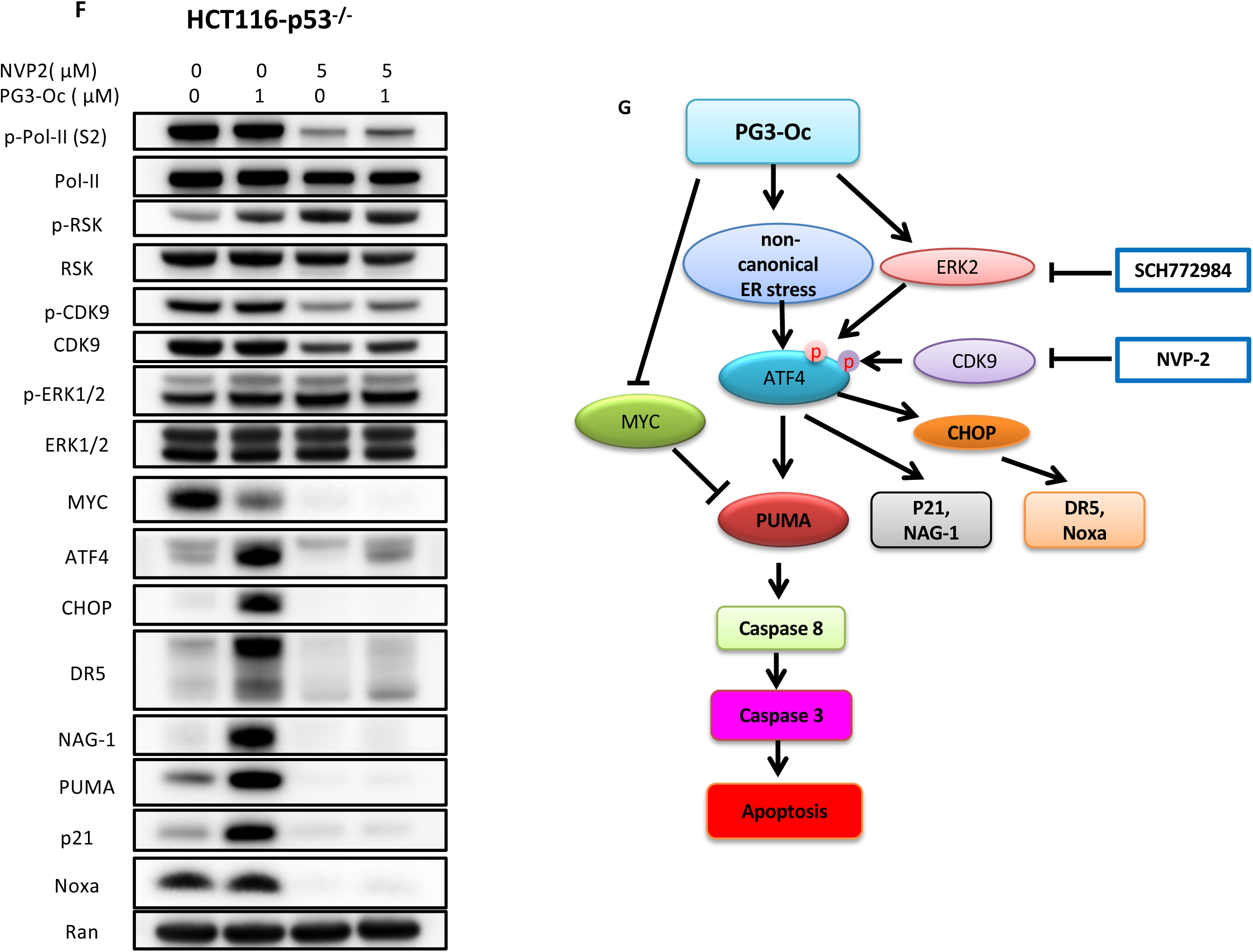
ERK1/2 and CDK9 kinase functions are required for ATF4 transcriptional activity. **(A)** HT29 cells were treated with PG3-Oc or SCH772984 (SCH)respectively, or co-treatment with PG3-Oc/SCH772984 for 28h. **(B)** HT29 cells were treated with PG3-Oc, trametinib (Tram), PLX4720 (PLX) and GSK2606414 respectively, or co-treatment with PG3-Oc/trametinib, PG3-Oc/PLX4720 and PG3-Oc/GSK2606414 for 24h. **(C)** HCT116 p53^-/-^ were treated with PG3-Oc, SCH772984, trametinib and PLX4720 respectively, or co-treatment with PG3-Oc/SCH772984, PG3-Oc/trametinib and PG3-Oc/PLX4720 overnight. **(D)** HT29 cells were treated with PG3-Oc, LDC 000067 (LDC) and NVP-2 respectively, or co-treatment with PG3-Oc/LDC000067 or PG3-Oc/NVP-2 overnight. HT29 **(E)** and HCT116 p53^-/-^ **(F)** cells were treated with PG3-Oc, or NVP-2 respectively, or co-treatment with PG3-Oc/NVP-2 overnight. (**G)** Proposed model of PG3-Oc-induced partial restoration of p53 pathway through ATF4 transcriptional modulation by ERK1/2 and CDK9.

In addition, IPA canonical pathway analysis indicates that the UPR signaling pathway is one of the top hits (second place) in siControl and PG3-Oc treated cells; after ATF4 knockdown, the UPR pathway moved down to 4^th^ place, suggesting suppression of UPR signaling (Figure S7 A and B). Importantly, the color code for UPR signaling pathway is grey, indicating that the pathway has no activity identified by the analysis, which is consistent with our observation that ATF4 is not activated through ER stress. Therefore, PG3-Oc-induced upregulation of ATF4 is through an unknown non-canonical ER stress pathway.

### ERK1/2 and CDK9 kinase functions are required for ATF4 transcriptional activity

Recent papers reported that phosphorylation of ATF4 by ERK2 or CDK9 stabilizes and promotes ATF4 nuclear translocation and ATF4 transcriptional function downstream of canonical ER stress (Ojha et al., 2019; Shiozaki et al., 2018). Phosphorylation sites of ATF4 by ERK2 or CDK9 are different (Ojha et al., 2019; Shiozaki et al., 2018). Our time-course experiments indicate that PG3-Oc treatment increases phosphorylation of ERK2 and its direct substrate RSK2 in both HT29 and HCT116 p53^-/-^ cell lines (Figure 6A and 6B), indicating ERK2 kinase activation. PG3-Oc does not increase the basal level of phosphorylation of CDK9 kinase (Figure 6B, 7B and 7E), suggesting the compound has no effect on CDK9 kinase activity.

We investigated whether ERK2 or CDK9 regulate non-canonical ER stress-induced ATF4 transcriptional activity. To address this question, we used the highly specific and potent ERK1/2 inhibitor SCH772984 and the CDK9 inhibitor NVP-2. SCH772984 significantly and NVP-2 completely blocks thapsigargin-induced upregulation of ATF4 and its target gene CHOP without inhibition of phosphorylation of eIF2α, demonstrating that ERK2 and CDK9 kinases function downstream of canonical ER stress (Figure 6D). That is consistent with the results of the publications mentioned above.

The ERK2 inhibitor blocks PG3-Oc-induced upregulation of ATF4 and CHOP, DR5, p21, Noxa and NAG-1 in both HT29 and HCT116 p53^-/-^ cell lines (Figure 7A and 7C), demonstrating ERK2 kinase activity is required for ATF4 stabilization and transcriptional function. As shown in Figure 7A and 7C, the ERK1/2 inhibitor treatment alone results in potent downregulation of MYC and upregulation of PUMA, therefor, blockage of PG3-Oc-induced PUMA upregulation by SCH772984 is not observed.

The ERK2 inhibitor has no effect on phosphorylation of CDK9 in either HT29 or HCT116 p53^-/-^ cell lines (Figure 7A and 7C), suggesting that CDK9 is not a substrate of ERK2. This is consistent with ERK2 phosphorylation sites of ATF4 being different from CDK9 phosphorylation sites of ATF4.

It is well-known that inhibition of MEK or Raf kinases leads to inhibition of ERK1/2 kinases. To further confirm these observations, HT29 and HCT116 p53^-/-^ cells were treated with MEK-specific inhibitor trametinib or mutant B-Raf (V600E)-specific inhibitor PLX4720 with or without PG3-Oc. The MEK inhibitor potently inhibits ERK1/2 kinases, indicated by potent inhibition of phosphorylation of both ERK1/2 and its substrate RSK2 in both HT29 and HCT116 p53^-/-^ cell lines (Figure 7B and 7C). PLX4720 potently inhibits ERK1/2 kinases in HT29 cells, as both phosphorylation of ERK1/2 and RSK2 are potently inhibited (Figure 7B), but not in HCT116 p53^-/-^ cells (Figure 7C). The HT29 cell line harbors mutant B-Raf (V600E), and HCT116 p53^-/-^ cell line has wild-type B-Raf which is not inhibited by PLX4720. Trametinib or PLX4720 impede PG3-Oc-induced upregulation of ATF4 and DR5, p21, NAG-1 and Noxa in HT29 cells, respectively, thus phenocopying the effects of the ERK1/2 inhibitor. Trametinib also prevents PG3-Oc-induced upregulation of ATF4 and DR5, p21, NAG-1 and Noxa in HCT116 p53^-/-^ cells, which again phenocopies the effects of the ERK1/2 inhibitor. On the other hand, PLX4720 does not inhibit wild-type B-Raf in HCT116 p53^-/-^ cells as shown in Figure 7C, where phosphorylation levels of both ERK1/2 and RSK2 are maintained as the untreated control, PG3-Oc-induces upregulation of ATF4. DR5, p21, NAG-1, Noxa and PUMA are not affected by PLX4720 treatment, which supports the idea that ERK2 kinase activity is required for ATF4 stabilization and transcriptional function. Also, trametinib or PLX4720 treatment alone leads to potent downregulation of MYC and upregulation of PUMA respectively, which phenocopies ERK1/2 inhibitor SCH772984 again (Figure 7B and 7C).

HT29 cells were treated with CDK9 inhibitor NVP-2 or LDC000067. LDC000067 impedes the upregulation of AFT4 and PUMA at 10 µM concentration. NVP-2 abolishes the induction of ATF4 and PUMA at both 1 µM and 5 µM concentrations. Interestingly, LDC000067 alone does not induce MYC downregulation at 10 µM concentration, but enhances PG3-Oc-induced MYC degradation. By contrast, NVP-2 alone and cotreatment with PG3-Oc potently downregulates MYC protein level (Figure 7D).

We noticed that PUMA was not upregulated though MYC was potently downregulated in HT29 (Figure 7D) and HCT116 p53^-/-^ cells (Figure 7F) because PG3-Oc-induced upregulation of ATF4 was abolished by the CDK9 inhibitor. These results support our model that both MYC degradation and ATF4 upregulation are required for PUMA induction by PG3-Oc.

Since NVP-2 is more potent and more specific than LDC000067, HT29 cell lysates treated with NVP-2 and NVP-2/PG3-Oc co-treatment from same experiment (Figure 7D) were used for further western blot analysis shown in Figure 7E.

HT29 (Figure 7E) and HCT116 p53^-/-^ cells were treated with NVP-2 or cotreated with NVP-2/PG3-Oc (Figure 7F). CDK9 directly phosphorylates Ser 2 of RNA polymerase II CTD (pol II) during elongation of the transcripts (Shiozaki et al., 2018). We observed that the phosphorylation of both CDK9 and RNA polymerase II CTD is potently inhibited by NVP-2 alone or combined treatment. Interestingly, NVP-2 does not show any effects on PG3-Oc-induced phosphorylation of ERK1/2, suggesting that ERK1/2 are not a substrate of CDK9. Importantly, inhibition of CDK9 abolishes PG3-Oc-induced upregulation of ATF4 and DR5, NAG-1, PUMA, p21, Noxa and CHOP.

In summary, inhibition of either ERK2 or CDK9 blocks PG3-Oc-induced upregulation and activation of ATF4. ERK2 and CDK9 together regulate ATF4 stability and transcriptional activities downstream of both thapsigargin-induced canonical and PG3-Oc-induced non-canonical ER stress. We propose a model of upregulation and activation of ATF4 by PG3-Oc and ATF4 mediates partial restoration of p53 pathway (Figure 7G).

### Lack of genotoxic stress by PG3-Oc

DNA damage induces the p53 pathway and leads to cell apoptosis. To study whether the p53 pathway restoration by compound PG3-Oc is due to DNA damage, we investigated the uptake and localization of PG3-Oc in cells. PG3-Oc and prodigiosin are red fluorescent compounds, and their localization in live cells can be monitored by fluorescence microscopy. We found that PG3-Oc and prodigiosin rapidly enter cells within 2 hours of incubation and remain in the cytosol at the 8-hour time point in HT29 and SW480 cells (Figure S8 A). Since we already observed that 1 μM PG3-Oc treatment for 8 hours can prominently induce the upregulation of PUMA mRNA (Figure 2 A, B and C) in HT29, SW480 and HCT116 p53^-/-^ cells, we investigated the DNA damage marker γ-H2AX (phospho Ser 139-histone H2AX) expression after the treatment for 8 hours. Western blot analysis shows that PG3-Oc and prodigiosin do not induce γ-H2AX in HT29 and SW480 cells at lower doses required for p53 pathway activation (Figure S8 B). Immunofluorescence staining shows that 1 μM PG3-Oc and prodigiosin does not induce γ-H2AX focus formation after the 8-hour treatment. By comparison, the DNA damaging chemotherapeutic drug CPT-11 used as a positive control significantly induces γ-H2AX foci in HT29 and SW480 cells (Figure S8 C). Both western blot and immunofluorescence staining data are consistent with the cytoplasmic localization of PG3-Oc. Our data indicate that the restoration of the p53 pathway by PG3-Oc at low concentrations does not show genotoxic effects in mutant p53-expressing cancer cells.

### *In vivo* studies of PG3-Oc demonstrate anti-tumor efficacy

To evaluate the anti-tumor effects of PG3-Oc *in vivo*, we established human tumor xenograft models by subcutaneous injection of human colon cancer cells into nude mice. After the tumor volume reached approximately 50 mm^3^, with HT29 xenografts, mice were treated by i.p. injection with vehicle or PG3-Oc at 5 mg/kg 3 times weekly for 2 weeks. The tumor volume in PG3-Oc-treated mice is significantly reduced as compared with vehicle-treated mice (Figure S9 A, B and C). No significant difference in body weight is observed between PG3-Oc and the vehicle treatment groups (Figure S9 D). Ki-67 expression is significantly decreased in PG3-Oc-treated tumors as compared with the vehicle group (Figure S9 E). PUMA is significantly induced in PG3-Oc-treated tumors as compared with controls (Figure S9 E). No *in vivo* toxicity is observed as indicated by H&E staining of tissues (Figure S9 F). These results indicate that PG3-Oc inhibits tumor growth in the HT29 mouse xenograft model.

With HCT116 p53^-/-^ xenografts, mice were treated by i.p. injection with vehicle or PG3-Oc·HCl at 7.6 mg/kg/day for biomarker studies (Figure S9 G, H and I). As shown in Figure S9 G, Western blot indicates that PG3-Oc·HCl induces significant upregulation of ATF4, PUMA and apoptosis markers of cleaved caspase 3 and cleaved PARP. IHC (immunohistochemistry) staining indicates that PG3-Oc·HCl induces upregulation of ATF4, PUMA and cleaved caspase 3 and downregulation of Ki-67 in HCT116 p53^-/-^ xenografts (Figure S9 H and I). However, tumor volume in treated mice appears to be not significantly different compared with vehicle-treated mice (Figure S9 J and K). No significant difference in body weight is observed between treated and the control groups (Figure S9 L).

We note that the conditions of the *in vivo* studies are not optimized, and we plan to perform more experiments in the future, including further studies of pharmacokinetics and toxicity of PG3-Oc, at different doses and treatment schemes, etc.

## Discussion

This is the first report on the compound PG3-Oc, a novel chemical entity (El-Deiry et al., 2017; issued composition of matter patent). Our manuscript describes therapy-induced p53-independent restoration of the p53 transcriptome. The prior literature typically investigated a few p53 targets and claimed p53 pathway restoration. It is necessary to evaluate the transcriptome and describe the extent of p53 pathway restoration by novel therapeutic candidates that act in a manner independent of p53. In this manuscript we have also evaluated the p53-independent p53 pathway-associated proteome by a novel p53 pathway restoring compound. There is little prior literature that has rigorously defined the p53-activated proteome following treatment by any drug and so we created an in-house p53-activated proteome data-base. An aspect of our approach that often gets misunderstood is the expectation that p53 pathway-restoring drugs act by altering mutant p53 protein and that such compounds have correlate activities in cells such as p53-specific genomic DNA-binding and chromatin association through mutant p53. This is not how the drugs that emerge from our functional p53-reporter screens work. Thus, this paper shows how a different transcription factor, ATF4 can “restore” p53 target gene-activation after drug treatment, essentially bypassing the defective mutated-p53 pathway. We evaluated the effect of the drug on the transcriptome and proteome. We show *in vivo* activity of the compound that is significant single-agent efficacy but this has not been optimized, and no maximally-tolerated dose (MTD) has been determined. This manuscript points to the ATF4 pathway with a specific therapeutic agent and claims partial (and significant) global p53-pathway restoration. The drug is also interesting because it targets Myc for degradation, an additional activity that we believe is relevant to its anti-tumor efficacy. Insights are included for non-canonical ER stress-independent activation of ATF4 by ERK1/2 and CDK9.

Restoration of the p53 pathway has been a long-term goal of the field of cancer research as an approach to treat tumors with mutated p53 and aggressive clinical behavior. The current work demonstrates the feasibility of the approach for small molecule drug-induced cellular reprogramming to achieve partial restoration of the global p53 transcriptome and proteome, in a p53-independent manner. Our dissection of the molecular components led us to identify ATF4 as a key transcription factor mediating the expression of p53 target genes in p53 mutant and null cancer cell lines after PG3-Oc treatment. Previously, ATF4 stabilization was thought to be regulated by ER stress (PERK-eIF2α signaling) or IRS (GCN2/PKR/HRI-eIF2α signaling), and phosphorylation of eIF2α at serine 51 is required. Very recently, it was discovered that mitochondrial protease ClpP (caseinolytic protease P) activation by ONC201 results in ATF4 increase without increase in phosphorylated eIF2a in Z138 cells (Ishizawa et al., 2019). In addition, regulation of ATF4 through other mechanisms was reported recently, such as phosphorylation of ATF4 by ERK2, RSK2 and CDK9 stabilizes ATF4 and promotes ATF4 translocation into the nucleus (Oh et al., 2010; Ojha et al., 2019; Shiozaki et al., 2018). We show that ATF4 shares a subset of p53 target genes that are involved in cell cycle arrest and apoptosis. This suggests that ATF4 is a critical node for responding to various intrinsic and extrinsic stresses and regulating cell fate.

Rescue of deficient or lost p53 function is an attractive strategy for cancer therapy. p53-restoring compounds usually act on specific p53 mutations, such as R273H (APR-246) or R175H (ZMC1) (Bykov et al., 2018). Toxicity, off-target effects and limited activity have been roadblocks for these different small molecules to progress to the clinic although progress is being made (Bykov et al., 2018). However, there is still a major unmet medical need to target tumors harboring mutant p53. Since thousands of mutations of p53 have been reported (Freed-Pastor and Prives, 2012; Olivier et al., 2010), these kinds of drugs targeting specific mutations may have a somewhat limited use in clinic. In this regard, functional restoration of the p53 pathway using small molecules regardless of what kind of p53 mutations exist in the tumor cells is an attractive method to target mutant p53-bearing tumors. PG3-Oc potently induces cancer cell death through ATF4-mediated restoration of p53 pathway in various mutant p53-expressing cancer cells, including single, double, multiple, truncated, frameshift mutations or p53-null cancer cells (Table 1, Figure 1, 2 and 3). These results indicate the versatility of a candidate therapeutic such as PG3-Oc to restoring the p53 pathway in cancer cells carrying various p53 mutants.

We propose a model (Figure 7G) in which activation of ATF4 through non-canonical ER stress by PG3-Oc results in upregulation of PUMA, DR5, p21 and NAG-1. Both ERK2 and CDK9 kinase activities are required for the stabilization and transcriptional activity of ATF4. We also identify that both MYC downregulation and ATF4 upregulation are required to induce upregulation of PUMA, and PUMA-mediated activation of caspase 8 causes cell apoptosis. It is clear that MYC downregulation occurs independently of ATF4 in PG3-Oc treated cells, as knockdown of ATF4 does not block MYC downregulation by PG3-Oc (Figure 5A and 5B). It is noteworthy that PG3-Oc targets MYC for degradation and that may contribute to its anti-cancer effect. As monotherapy, PG3-Oc shows significant anti-tumor efficacy with lack of toxicity in our *in vivo* studies.

The activation of ATF4 as a drug-induced mechanism to partially restore the p53 pathway provides an understanding for how treatment of cancer cells by a small molecule can restore critical anti-tumor p53 signaling in cells with mutant p53 and without involvement of p53 family members such as p73 or p63. While largely non-overlapping the ATF4 and p53 transcriptomes appear to overlap at a critical set of effector genes that confer anti-tumor properties such as apoptosis induced by PUMA. Moreover, while each transcription factor regulates a set of genes that mediate its cellular effects, there are some common themes even where the additional gene sets are non-overlapping.

A major effect of PG3-Oc is to activate ATF4 which plays an important role in communicating pro-survival and pro-apoptotic signals. ER stress, the integrated stress response and mitochondrial stress lead to activation of ATF4. As a transcriptional factor, ATF4 regulates a transcriptional program involved in upregulation of p21 for cell cycle arrest and senescence; PUMA, DR5 and Noxa for apoptosis; ATG5 and ATG7 for autophagy, which are similar with p53. On the other hand, ATF4 also positively regulates gene expression involved in antioxidant response to protect cell from ROS (oxidative stress), and ER-associated degradation (ERAD) pathway for degradation of abnormal proteins, and re-establishment of cellular homeostasis. ATF4 and MYC co-regulate 30 MYC-target genes involved in amino acid biosynthesis and protein synthesis. These are different from p53. Final outcome of ATF4 activation is dependent on the cell type, nature of stressors and duration of the stresses (Ojha et al., 2019; Tameire et al., 2019; Wang et al., 2015; Wortel et al., 2017).

Future work can focus in more detail on comparisons of the global and gene-specific regulation between ATF4 and p53, for example through ChIP-seq and single cell analysis. It will be important to unravel whether the PG3-Oc anti-tumor drug effects are primarily due to the partial restoration of the p53-transcriptome and -proteome that includes critical effector genes such as PUMA or whether other genes within the ATF4-transcriptome contribute to the drug efficacy. The insights from our paper could provide the basis for novel drug screens that optimize further the anti-tumor properties of both transcription factors, or investigate anti-cancer cooperativity in tumors that retain wild-type p53.

In the future it would be of interest to investigate similarities and differences in the ATF4-activated transcriptomes and general transcriptomes and proteomes between small molecule compounds such as ONC201 and PG3-Oc as both upregulate ATF4 although through different upstream pathways. ONC201 is not known as a p53-pathway restoring compound although it was discovered as a TRAIL inducing compound (TIC10) and later found to induce DR5 through ATF4 and an integrated stress response involving HRI and PKR (Allen et al., 2013; Kline et al., 2016). Thus, as there is more focus on ATF4 as a therapeutic target in cancer it will be important to understand the downstream drug effects that it mediates through different upstream regulators, as well as ATF4-independent effects, e.g. MYC suppression in the case of PG3-Oc. It will also be of interest to determine whether PG3-Oc relies on mitochondrial mechanisms involving ClpX and ClpP for ATF4 activation (Ishizawa et al., 2019)

In summary, our results demonstrate that a small molecule can stimulate global p53 pathway restoration in tumor cells with mutated p53 or cells that are null for p53. This occurs in a p53 family-independent manner by PG3-Oc which impacts on a relevant transcriptome and proteome leading to tumor cell death. The mechanism of p53 pathway restoration by PG3-Oc involves activation of ATF4, which has a largely non-overlapping transcriptome with p53, but nonetheless activates critical targets required for drug-induced tumor suppression, including PUMA. The involvement of ATF4 in a partial global p53 pathway restoration represents a novel mechanism for therapy-induced molecular reprogramming to achieve an anti-cancer effect that may be translatable to the clinic.

## Materials and Methods

### Cell lines and reagents

P53-mutant cell lines: HT29 (R273H), SW480 (R273H/P309S), DLD-1 (S241F), H1975 (R273H), MDA-MD-231 (R280K), U251 (R273H), FaDu (R248L), CAL-27 (H193L), PANC-1 (R273H), Aspc-1 (frameshift mutation), P53 wild-type cell lines: HCT116, and CCD 841 Con; P53-null cell line: HCT116 p53^-/-^. H1975 and CCD 841 Con cells were purchased from ATCC. HT29, SW480, DLD-1, HCT116, FaDu, CAL-27, MDA-MD-231, PANC-1 and Aspc-1 and Jurkat cell lines were purchased from Fox Chase Cancer Center cell culture facility. HCT116 p53^-/-^ cell lines were from the Vogelstein laboratory at Johns Hopkins. Cells were verified to be MYCoplasma-free at multiple times throughout the study. We routinely checked for MYCoplasma and all cell lines underwent STR authentication. Chemicals: Caspase 8 inhibitor Z-IETD-fmk (BD Bioscience), LDC000067, NVP-2, and thapsigargin (Tocris Bioscience), MG-132, 5-FU, trametinib, PLX-4720, Z-VAD-fmk, GSK2606414, SCH772984, regorafenib, SB216763 and SP600125 (Selleck chemicals).

### Western blotting

After treatment, protein lysates were collected for Western blot analysis. A total of 15 μg of protein was used for SDS-PAGE. After primary and secondary antibody incubations, the signal was detected by a chemiluminescence detection kit, imaged by Syngene (Imgen Technologies). Antibodies for PUMA (for IHC), NAG-1 (GDF15), P53 were from Santa Cruz Biotechnology; for caspase 8, cleaved caspase 8, caspase 9, caspase 3, cleavage PARP, eIF2α, p-eIF2α (Ser51), CHOP, ATF4, DR5, FOXO3a, p-FOXO3a (Ser253), NF-κB p65, p-NF-κB p65 (Ser536), c-Jun, p-c-Jun (Ser63), JNK, p-JNK (Thr183/Tyr185), PUMA (for WB), MYC, phosphor-S62-cMYC, NDRG1, Phospho-CDK9 (Thr86), CDK9, Rpb1 NTD (RNA PII subunit B1), phosphor-(Ser2) Rpb1 CTD (RNA PII subunit B1), RSK and phospho-p90RSK(Ser380) were from Cell Signaling Technology. Noxa and p21 were from Calbiochem. p73 was from Bethyl laboratories Inc., Ran was from BD Biosciences. β-actin was from Sigma.

### Cell viability assay

Cells were seeded in 96-well plates (6 x 10^3^ cells/well). Cells were treated with different concentrations of compounds or dimethyl sulfoxide (DMSO) as a control for 72 hours. The cell viability was assessed by CellTiterGlo bioluminescent cell viability assay (Promega), following the manufacturer’s protocol. Bioluminescence imaging was measured using the IVIS imager. Percentage of cell viability (mean ± SEM) at each dose was calculated against the respective DMSO control. The IC_50_ values were determined from the sigmoidal dose– response curves using GraphPad Prism.

### Caspase activity assay

Cells were seeded in 96-well plate (1 x 10^4^ cells/well). Cells were treated with different concentrations of compounds or dimethyl sulfoxide (DMSO) as a control for 24 hours. Caspase 3/7 activity was assessed by the Caspase-Glo® 3/7 Assay kit (Promega), following the manufacturer’s protocol. Bioluminescence imaging was measured using the IVIS imager. Caspase activity was normalized to cell numbers and compared to those of the DMSO treatment control in each cell line. Data is reported as mean RLU + SEM (n=3).

### Colony formation assays

Five hundred cells were seeded per well in 6-well plates and treated with different concentrations of compounds for 24 hr, then, cells were cultured with drug-free complete medium for 2 weeks with fresh medium changed every 7 days. Cells were fixed with 10% formalin and stained with 0.05% crystal violet at the end of 2 weeks period of cell culture (Franken et al., 2006).

### Cell uptake and localization

A total of 5 × 10^4^ cells was seeded in each well of 8-well chamber slides. Cells were incubated with PG3-Oc for 2 and 8 hours respectively, washed and fixed by 4% paraformaldehyde for 15 minutes at room temperature, washed, stained with DAPI for 10 minutes, mounted, and examined by fluorescence microscopy.

### Immunofluorescence staining

A total of 5 × 10^4^ cells was seeded in each well of 8-well chamber slides. After treatment, cells were fixed and permeabilized by methanol:acetone (1:1) for 20 minutes at −20⁰C. Fixed cells were blocked by 2% BSA for 1 hour, followed by primary antibody incubation for 1 hours and Cy3-conjuated secondary antibody incubation for 1 hour at room temperature. After washing, cells were stained with DAPI for 10 minutes at room temperature. Cells were mounted, and examined by fluorescence microscopy.

### Flow cytometry assay

#### Cell Cycle Analysis

Propidium iodide (PI) staining and flow cytometry were used to determine the degree of cellular apoptosis. Cells were seeded at 3 x 10^5^ cells/well in six-well plates. Cells were treated with PG3-Oc for 48 h. Cells were harvested, fixed by 70% ethanol, and stained by propidium iodide, then flow cytometry was performed as previously described (28). The percentage of hypo-diploid cells (sub-G1) was used to quantify dead cells in apoptosis assays.

### qRT-PCR

Total RNA was isolated from PG3-Oc-treated cells using the Quick-RNA mini prep kit (Zymo Research, Irvine, CA) according to the manufacturer’s protocol. 500 ng of total RNA was used to generate cDNA using SuperScript III first-strand synthesis system with random primers (Invitrogen), following manufacturer’s protocol. Real-time PCR was performed using POWER SYBR GREEN mast mix (Applied Biosystem) for *DR5, p21, PUMA* and *GAPDH*, and TaqMan primer-probes for detection of MYC mRNA levels on 7900HT Sequence Detection System (Applied Biosystem). PUMA primer (forward, 5’-GAC-GAC-CTC-AAC-GCA-CAG-TA-3’; reverse, 5’-AGG-AGT-CCC-ATG-ATG-AGA-TTG-T-3’), DR5 primer (forward, 5-ACAGTTGCAGCCGTAGTCTTG-3’,; 5’-CCAGGTCGTTGTGAGCTTCT-3), GAPDH primer (forward, 5ʹ-TCG ACA GTC AGC CGC ATC TTC TTT-3ʹ; reverse, 5ʹ-ACC AAA TCC GTT GAC TCC GAC CTT-3ʹ), Taq Prob IDs for *MYC* (HS 00153408) and *GAPDH* (HS 99999905). ΔΔCt method was used to analyze and report fold-changes of the indicated genes.

### siRNA knockdown

Knockdown experiments were performed by transfecting either 80 pmoles of indicated siRNA(s), or scramble siRNA using RNAiMAX (Invitrogen). Transfected cells were treated with PG3-Oc, 24 hrs post-transfection. The control scrambled siRNA and siRNA for human ATF4, CHOP, DR5, Puma, NF- κB p65 and MYC were purchased from Santa Cruz Biotechnology. p73 siRNA was from Ambion, and FOXO3a siRNA from Thermo Scientific Dharmacon.

### Transfection of plasmids

Cells were transfected with MYC expression plasmids (Ricci et al., 2004) and vector pcDNA3 (Invitrogen) using Lipofectamine 2000 (Invitrogen) according to the manufacturer’s instruction.

### RNA-Seq analysis

RNA-sequencing was performed by Fox chase Cancer Center genomics facility (333 Cottman Ave, Philadelphia, PA 19111) and Genewiz (115 Corporate Boulevard, South Plainfield, NJ 07080). HT29 cells were treated with or without 1 μM PG3-Oc in triplicate for 24 hours. HCT116 and HCT116 p53^-/-^ were treated with 50 μM 5-FU for 24 hours. Total RNA was isolated using RNeasy Mini kit (Qiagen). RNA concentration and quality was analyzed using a NanoDrop 2000. RNA integrity of each sample was analyzed on a Bioanalyzer (Agilent).

#### Reagents

Truseq stranded mRNA library kit, Hiseq rapid SRcluster kit, HiSeq rapid SBS kit (Illumina,CA). Equipment: HiSeq2500 sequencer (Illumina, CA).

Stranded mRNA-seq library: 1000ng total RNAs from each sample were used to make library according to the product guide. In short, mRNAs were enriched twice via poly-T based RNA purification beads, and subjected to fragmentation at 94 degree for 8 min via divalent cation method. The 1st strand cDNA was synthesized by Superscript II and random primers at 42 degree for 15 mins, followed by 2nd strand synthesis at 16 degree for 1hr. During second strand synthesis, the dUTP was used to replace dTTP, thereby the second strand was quenched during amplification. A single ‘A’ nucleotide is added to the 3’ ends of the blunt fragments at 37 degree for 30 min. Adapters with illuminaP5, P7 sequences as well as indices were ligated to the cDNA fragment at 30 degree for 10 min. After Ampure bead (BD) purification, a 15-cycle PCR reaction was used to enrich the fragments. PCR was set at 98 degree for 10 sec, 60 degree for 30 sec and extended at 72 degree for 30 sec. Libraries were again purified using AmPure beads, had a quality check on bioanalyzer (Agilent) and quantified with Qubit (Invitrogen). Sample libraries were subsequently pooled and loaded to the sequencer. Single end reads at 100bp were generated for the bioinformatic analysis.

#### Bioinformatics analysis

Pathway and network analysis (cut-off is 2-fold and above) by Ingenuity Pathway Analysis (IPA) (Qiagene) was performed to identify key biological processes, canonical pathways, upstream transcriptional regulators and gene networks. Gene Set Enrichment Analysis was performed by ranking genes first by highest to lowest log 2-fold change. The ranked gene list was then queried using GSEA software to known Molecular Signature Database (MsigDB). Known pathways from curated databases and published studies that matched our gene signature were then reported in the analysis.

### Proteomic analysis

#### Sample preparation for LC-MS/MS analysis

Cell pellets (HCT116, HCT116 p53^-/-^ and HT29) were lysed with a lysis buffer (8 M urea, 1 mM sodium orthovanadate, 20 mM HEPES, 2.5 mM sodium pyrophosphate, 1 mM β-glycerophosphate, pH 8.0, 20 min, 4°C) followed by sonication at 40% amplification by using a microtip sonicator (QSonica, LLC, Model no. Q55) and cleared by centrifugation (14 000 × g, 15 min, 15°C). Protein concentration was measured (Pierce BCA Protein Assay, Thermo Fisher Scientific, IL, USA) and a total of 100 µg of protein per sample was subjected for trypsin digestion. Typtic peptides were desalted using C18 Sep-Pak plus cartridges (Waters, Milford, MA) and were lyophilized for 48 hours to dryness. The dried eluted peptides were reconstituted in buffer A (0.1 M acetic acid) at a concentration of 1 µg/µl and 5 µl was injected for each analysis.

The LC-MS/MS was performed on a fully automated proteomic technology platform (Ahsan et al., 2017). that includes an Agilent 1200 Series Quaternary HPLC system (Agilent Technologies, Santa Clara, CA) connected to a Q Exactive Plus mass spectrometer (Thermo Fisher Scientific, Waltham, MA). The LC-MS/MS set up was used as described earlier (Ahsan et al., 2017, J Proteomics 2017, 165: 69-74). Briefly, the peptides were separated through a linear reversed-phase 90 min gradient from 0% to 40% buffer B (0.1 M acetic acid in acetonitrile) at a flow rate of 3 µl /min through a 3 µm 20 cm C18 column (OD/ID 360/75, Tip 8 µm, New objectives, Woburn, MA) for a total of 90 min run time. The electrospray voltage of 2.0 kV was applied in a split-flow configuration, and spectra were collected using a top-9 data-dependent method. Survey full-scan MS spectra (m/z 400-1800) were acquired at a resolution of 70,000 with an AGC target value of 3×10^6^ ions or a maximum ion injection time of 200 ms. The peptide fragmentation was performed via higher-energy collision dissociation with the energy set at 28 normalized collision energy (NCE). The MS/MS spectra were acquired at a resolution of 17,500, with a targeted value of 2×10^4^ ions or maximum integration time of 200 ms. The ion selection abundance threshold was set at 8.0×10^2^ with charge state exclusion of unassigned and z =1, or 6-8 ions and dynamic exclusion time of 30 seconds.

#### Database search and label-free quantitative analysis

Peptide spectrum matching of MS/MS spectra of each file was searched against the human database (UniProt) using the Sequest algorithm within Proteome Discoverer v 2.3 software (Thermo Fisher Scientific, San Jose, CA). The Sequest database search was performed with the following parameters: trypsin enzyme cleavage specificity, 2 possible missed cleavages, 10 ppm mass tolerance for precursor ions, 0.02 Da mass tolerance for fragment ions. Search parameters permitted variable modification of methionine oxidation (+15.9949 Da) and static modification of carbamidomethylation (+57.0215 Da) on cysteine. Peptide assignments from the database search were filtered down to a 1% FDR. The relative label-free quantitative and comparative among the samples were performed using the Minora algorithm and the adjoining bioinformatics tools of the Proteome Discoverer 2.3 software. To select proteins that show a statistically significant change in abundance between two groups, a threshold of 1.5-fold change with p-value (0.05) were selected.

### Knock-out of PUMA by CRISPR/Cas9 gene editing

#### sgRNA design and plasmid construction

sgRNA targeted the exon 3 of PUMA gene, which contains sequence code for BH3 domain of PUMA. Two crRNAs introduced into lentiviral vectors (pLentiCRISPR-E, Addgene #78852) which contains eSpCas9 and puroMYCin cassette.

Guide1 DNA (forward, 5’-CACC <u>GGCGGGCGGTCCCACCCAGG-3’</u>; reverse, 5’-AAAC <u>CCTGGGTGGGACCGCCCGCC</u>-3’) and Guide 2 DNA (forward, 5’-CACC <u>GCCGCTCGTACTGTGCGTTG</u>-3’; reverse, 5’-AAAC <u>CAACGCACAGTACGAGCGGC-3’)</u> were annealed and ligated to the restriction enzyme-cut plasmid by T4 ligase. Stbl3 strain (Invitrogen C7373-03) was transformed by the guides-containing plasmids. LB-amp plates were streaked and incubated on a shaker at 37°C overnight. The bacterial colonies were selected and mixed up with LB (Terrific Broth) and 100 μg/mL ampicillin, and were incubated on a shaker at 37°C overnight. Plasmids from different colonies were isolated and purified using QIAprep Spin Miniprep Kit (Qiagen). Plasmids were digested with BsmBI and BamHI in Cut Smart Buffer (New England BioLabs, Inc.) at 37°C for 1 hour and then analyzed by 1% agarose gel. Sequencing was performed by GENEWIZ (South Plainfield, New Jersey, NJ; Figure S5 A-F).

#### Cell culture, DNA transfection

Lentivirus was generated with psPAX2, pVSV-G and the pLentiCRISPR plasmids that contain the guides and Cas9 in 293T cells. 48 hours later, all supernatant was transferred to a 1.5 mL tube that was centrifuged to remove debris. The supernatant was transferred to a new 1.5 mL tube, and stored at 4°C. HT29 cells were transfected with the lentivirus supernatant and polybrene was added to enhance the transfection. PuroMYCin (final concentration is 1μg/mL) was added to the medium to select positive cells.

#### Mutation screens by Sanger sequencing and TIDE analysis

DNA was extracted and purified from positive HT29 cells using DNeasy Blood & Tissue kit (Qiagen). PCR primers that frank both sides of the exon 3 of PUMA gene were used to amplify the target region (forward, 5’-CACAGTCTCTGGCCTTCTGG-3’; reverse, 5’-AGCTGCCGCACATCTGG-3’). The amplicon is GC-rich region, to improve PCR specificity, we performed temperature gradient PCR to optimize annealing temperature. A hot-start and touch-down PCR with accuPrime™ Pfx DNA Polymerase (ThermoFisher Scientific) and 2.5% DMSO and 1M betaine, was performed to achieve specific amplification of target region. The PCR products were purified by QIAquick PCR purification kit (Qiagen) for Sanger sequencing. TIDE analysis was performed using an online tool (TIDE: Tracking of Indels by DEcomposition, https://tide-calculator.nki.nl/). Sequencing was performed by GENEWIZ (South Plainfield, New Jersey, NJ; Figure S5 C).

#### Single cell colonies

300 positive HT29 cells were placed into a 10 cm dish and incubated at 37°C. After 2 weeks, single cell colonies were selected and expanded. Western blotting using PUMA antibody was performed to screen the colonies (Figure S5 E and F).

### *In vivo* anti-tumor assay and Immunohistochemistry

All animal experiments were approved by the Institutional Animal Care and Use Committee at Fox Chase Cancer Center and Brown University. One million HT29 or HCT116 p53^-/-^ cells were implanted subcutaneously in the flanks in each athymic nude mouse (female, 5–6 weeks old). The mice with HT29 tumor xenografts were divided at random into two groups and treated with the vehicle (10% DMSO, 20% Kollipher EL in PBS) and PG3-Oc (5 mg/kg, 3 times/wk) by intraperitoneal injection when the tumor masses reached a size of 5 to 6 mm (Supplemental Figure S9 A). The mice with HCT116 p53^-/-^ tumor xenografts were divided at random into two groups and treated with the vehicle (10% DMSO, 20% Kollipher EL in PBS) and PG3-Oc·HCl (7.6 mg/kg/day) by intraperitoneal injection when the tumor masses reached a size of 5 to 6 mm (Figure S9 H and J). Subsequently tumor volumes were measured with a caliper and calculated using V=0.5 x Length x Width^2^. Twenty-three days after treatment, the mice were euthanized and tumors were excised. H&E staining and Immunohistochemistry (IHC) of paraffin-embedded tumor and tissue sections were performed at the Fox Chase Cancer Center Histopathology Facility and Brown University Alpert Medical School Molecular Pathology Core. Antibodies for IHC: PUMA (Santa Cruz Biotechnology), Ki-67, DR5 and cleaved Caspase 3 (Cell Signaling Technology), ATF4 (Abcam).

### Statistical analysis

All results were obtained from triplicate experiments, unless other indicated. Statistical analyses were performed using PRISM4 Software (GraphPad Software, Inc.), and the Student *t* test. Statistical significances were determined by *P* < 0.05.

## Acknowledgements

The authors thank the Fox Chase Cancer Center genomics, Flow cytometry cores and bioinformatics facility for help with cell cycle analysis, real-time PCR experiments and RNA-Seq data analysis. This work has been presented in part at the 2017, 2018, and 2020 AACR meetings. W.S.E-D. is an American Cancer Society Research Professor and is supported by the Mencoff Family Professorship at Brown University.

## Author Contributions

X.T., and W.S.E-D. conceptualized the project and all experiments that were performed. X.T. was involved with the technical performance of all experiments. X.T., and W.S.E-D. were involved in all of the data analysis and discussion of the results. N.A. performed proteomic experiments and analyzed proteomic dataset. S.Z. assisted with early compound testing. A. Lulla assisted with some Q-RT-PCR experiments and *in vivo* studies. A. Lev assisted with animal experiments. P.A. assisted with the CRISPR cloning strategy and experiments. D.T.D. assisted with flow cytometry. All authors were involved in writing and editing of the manuscript. W.S.E-D. was responsible for administrative oversight of the research, securing funding for the project, and overall conduct of the experiments.

## Supplementary Materials

There are 9 supplementary Figures with Legends included. In addition, there are an additional 22 source files that can be made available as spreadsheets/excel files that are relevant to the bioinformatic analyses.

## Supplementary Figure Legends

**Figure S1.**
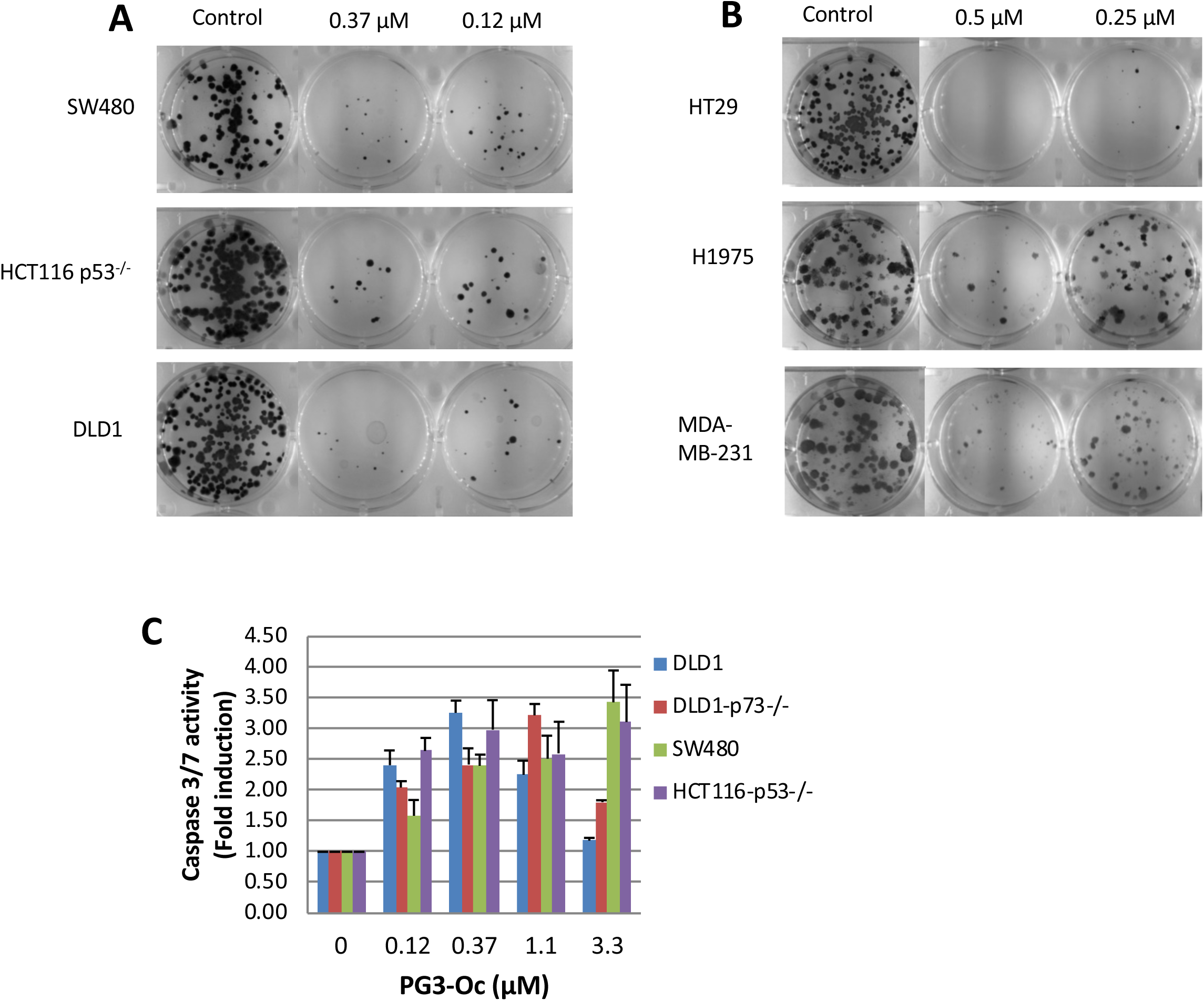
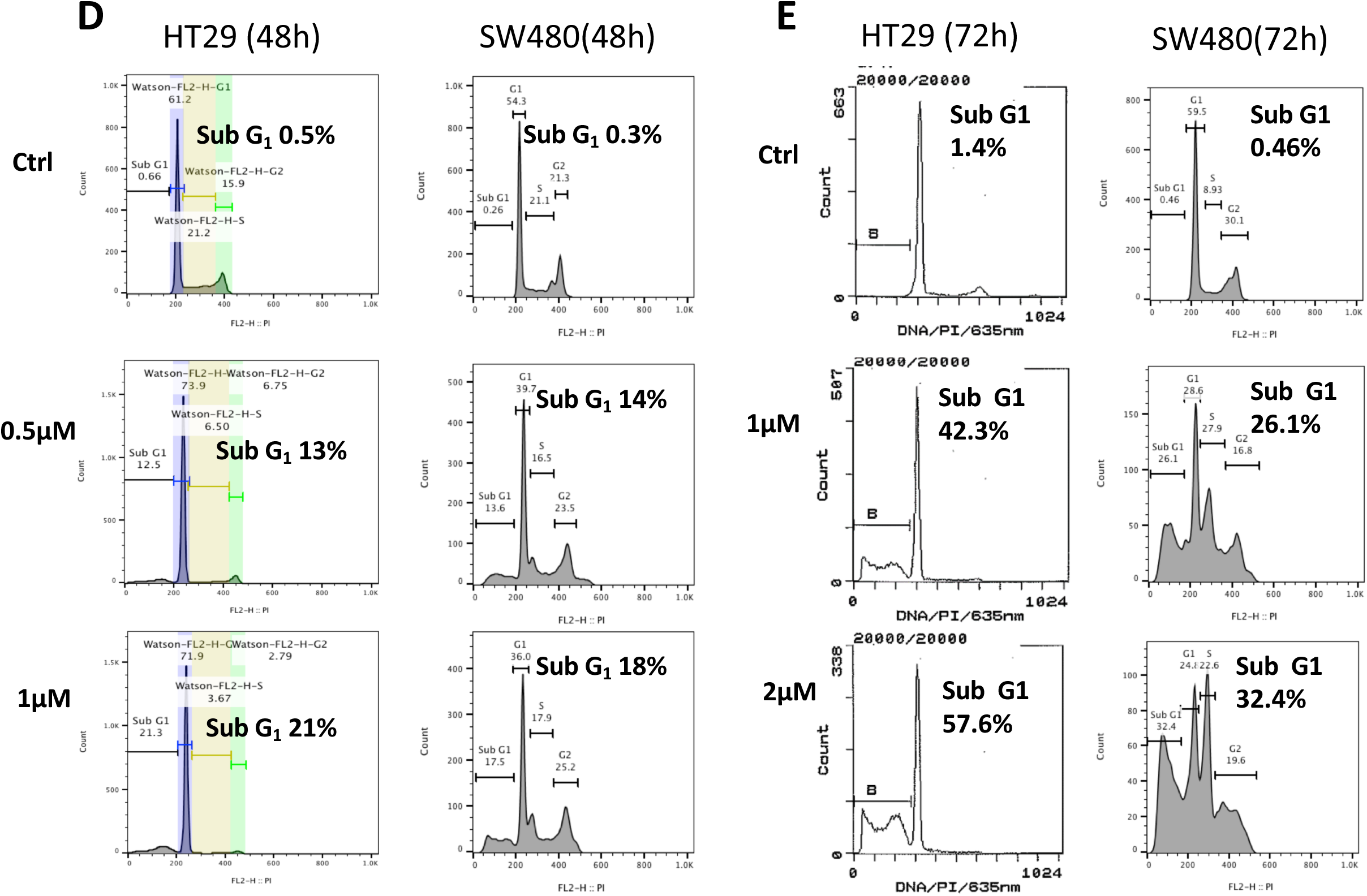
PG3-Oc inhibits cell proliferation and induces apoptosis in mutant p53-expressing cancer cell lines. (**A)** and (**B)** Colony formation assay of p53-mutant and p53-null human cancer cells. Cells were treated with the indicated concentrations of PG3-Oc for 24 hr, and then cultured in drug-free media for 14 days at which time crystal violet staining was performed on the attached cells. **(C)** Caspase 3/7 activity assay. Cells were treated with PG3-Oc at the indicated concentrations for 24 hr. Luciferase activity was imaged by the IVIS Imaging System after treatment. Caspase activity data (performed in triplicate) were normalized to cell numbers and to the DMSO treatment control for each cell line and data analyses were performed using Excel. **(D)** and **(E)** Cell-cycle profiles after PG3-Oc treatment. Apoptosis was analyzed by nuclear PI-staining using flow cytometry. HT29 and SW480 cells were treated with PG3-Oc at the indicated concentrations for 48 hr or 72 hours respectively.

**Figure S2.**
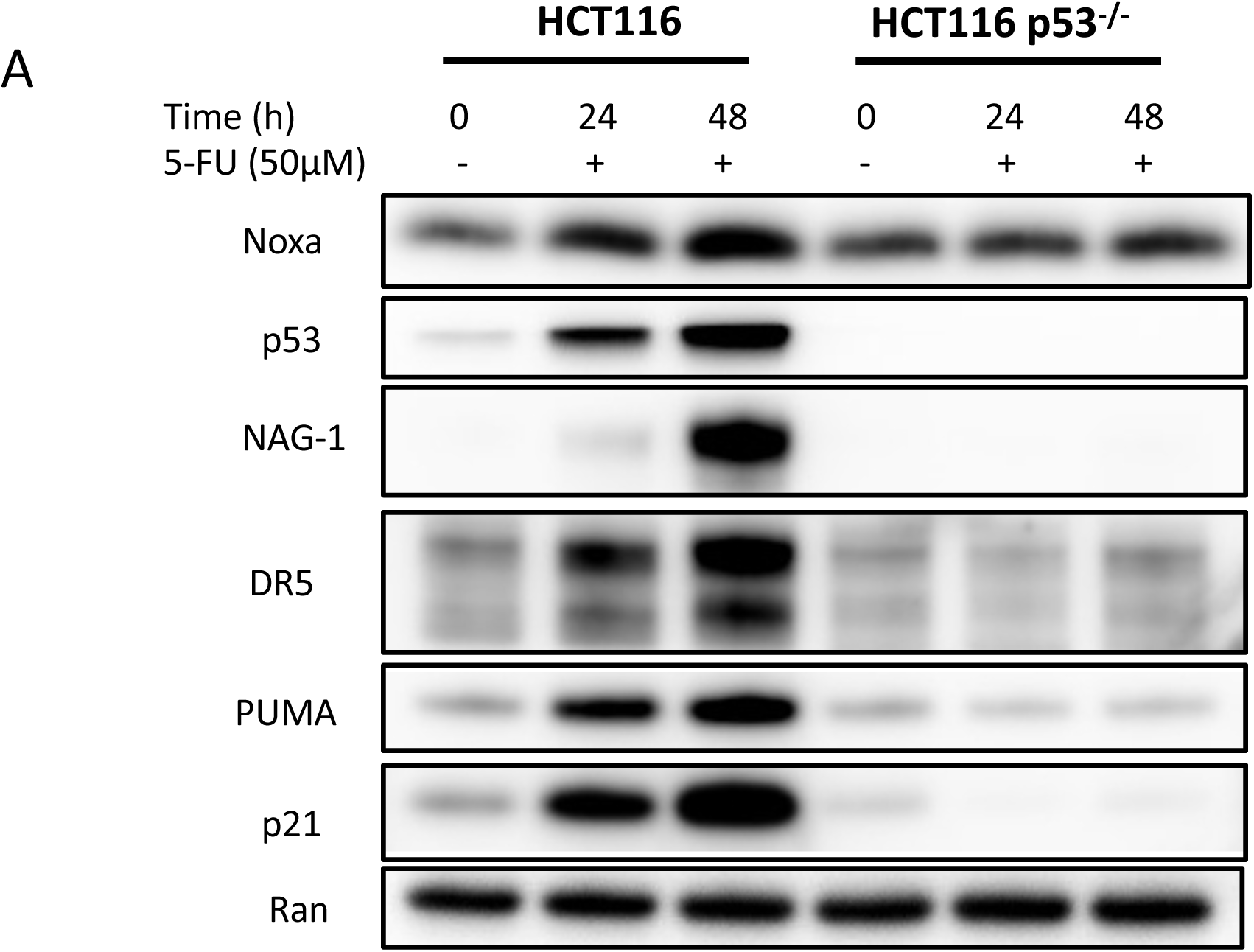
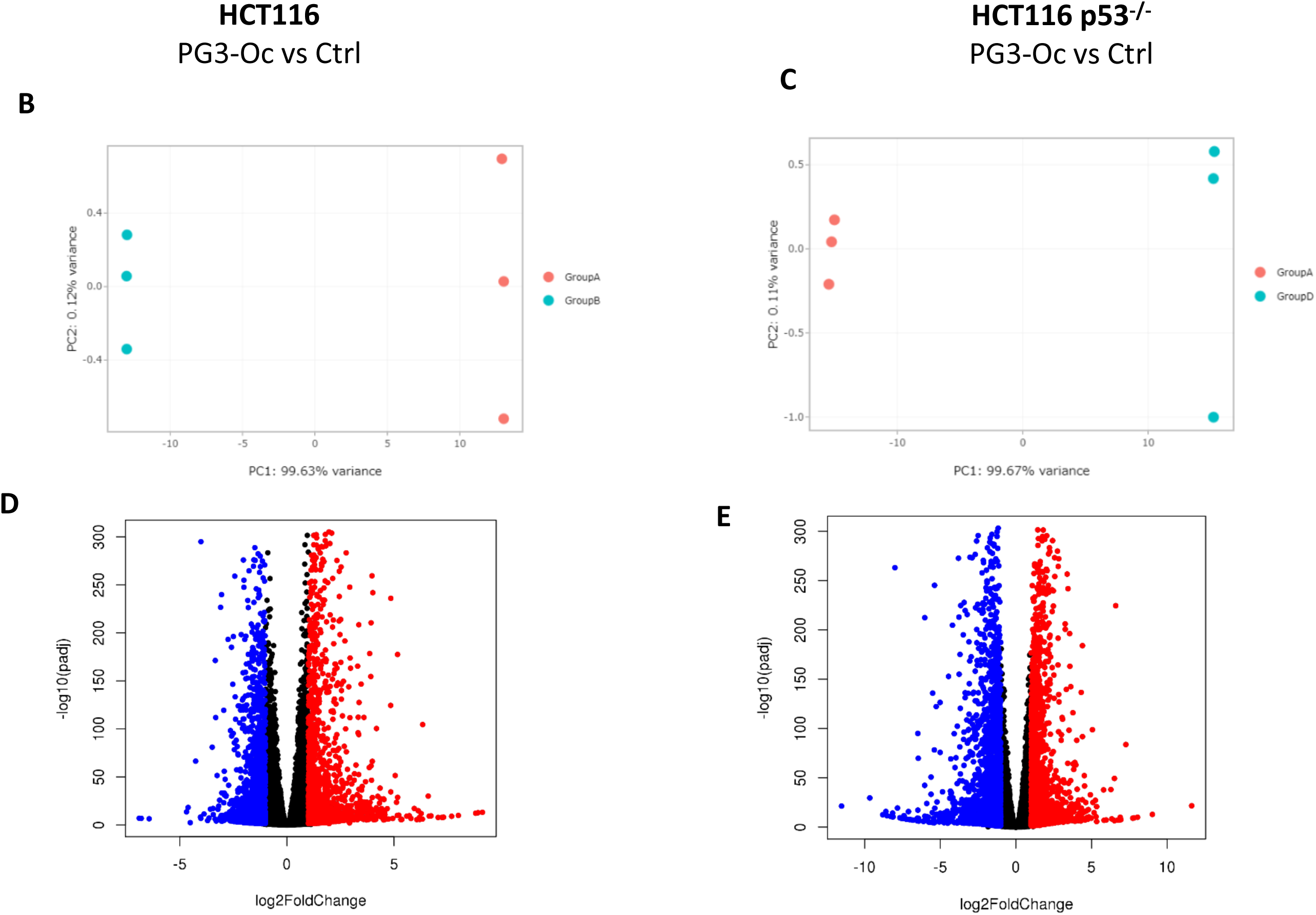
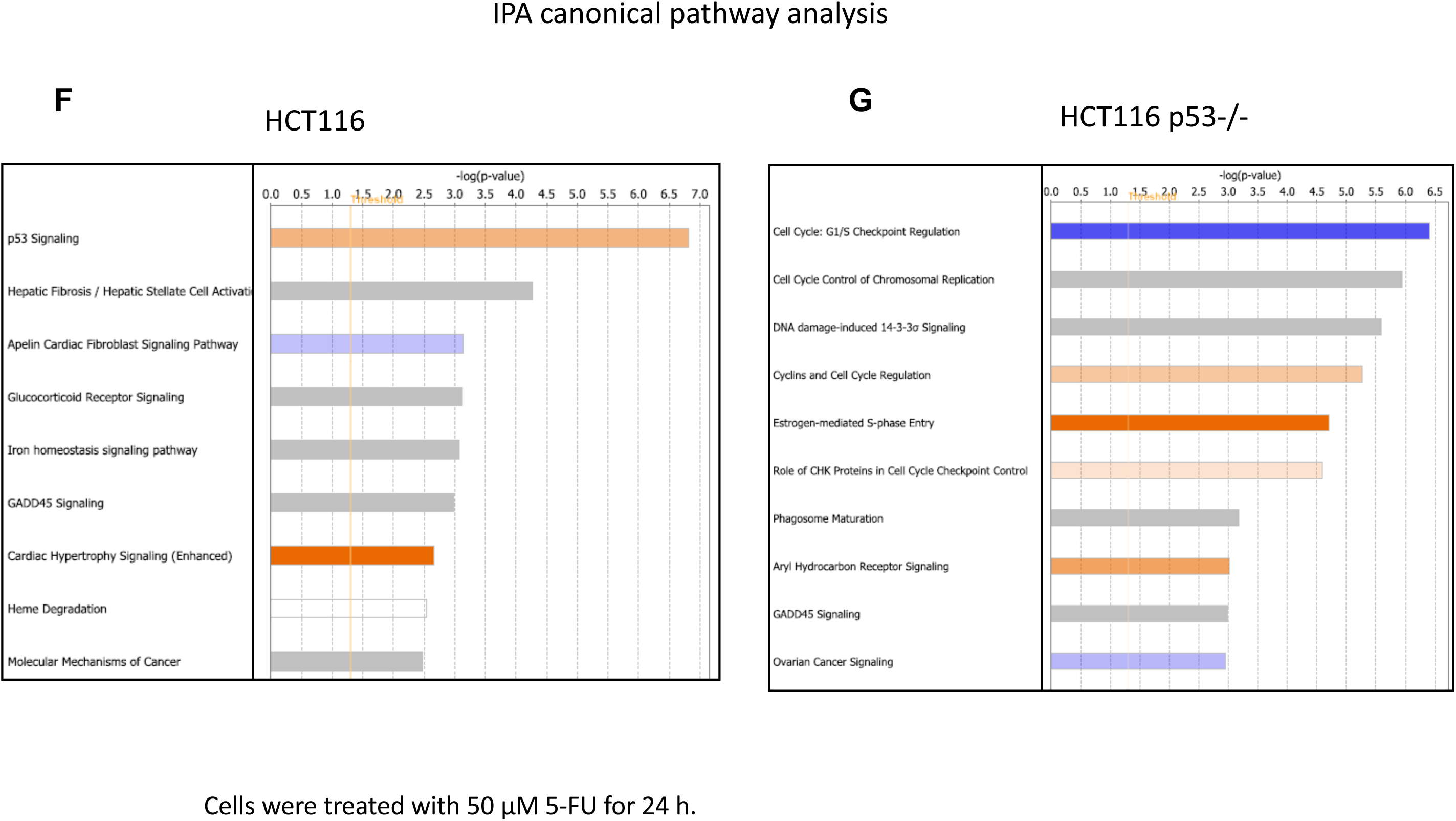
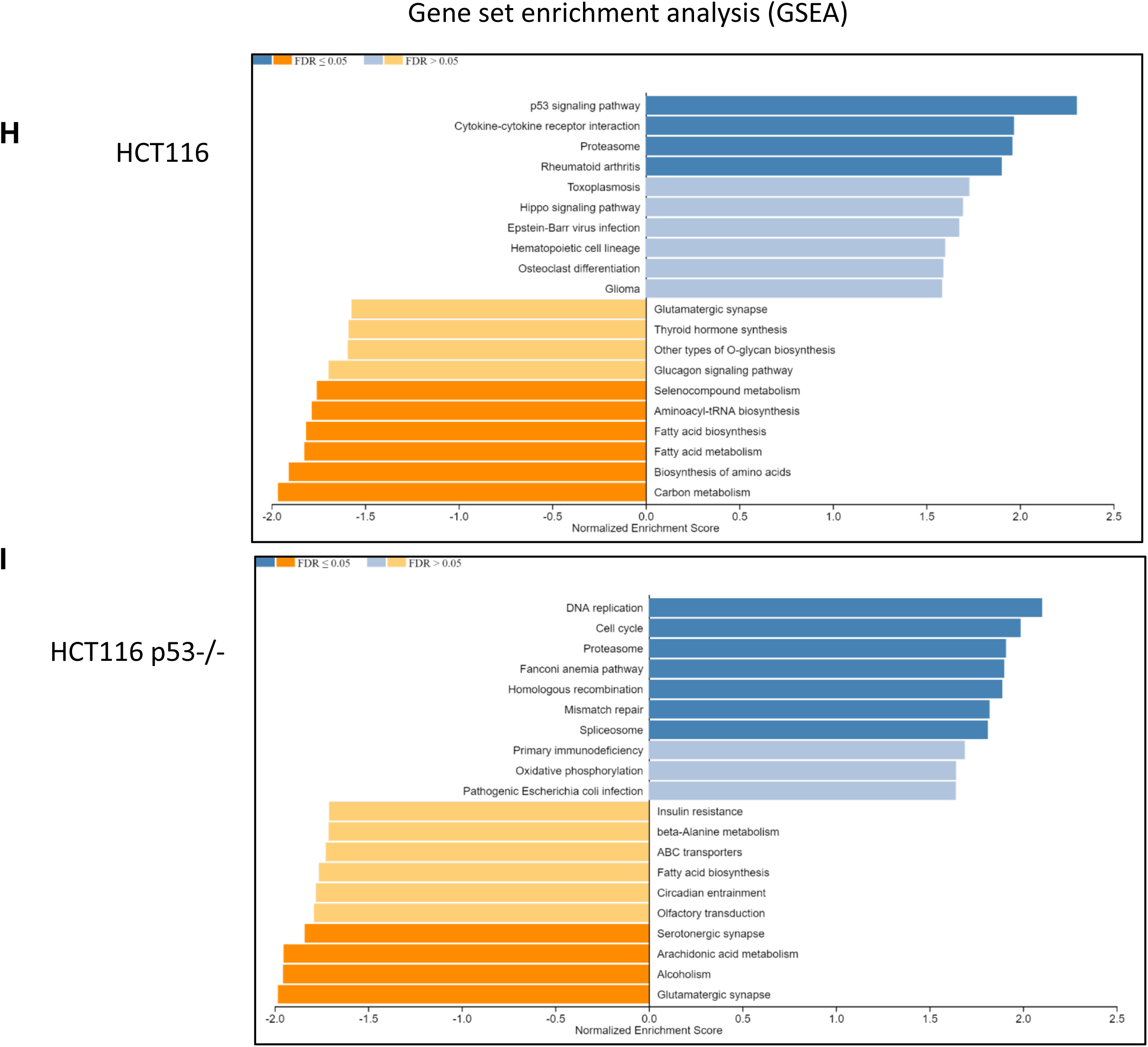
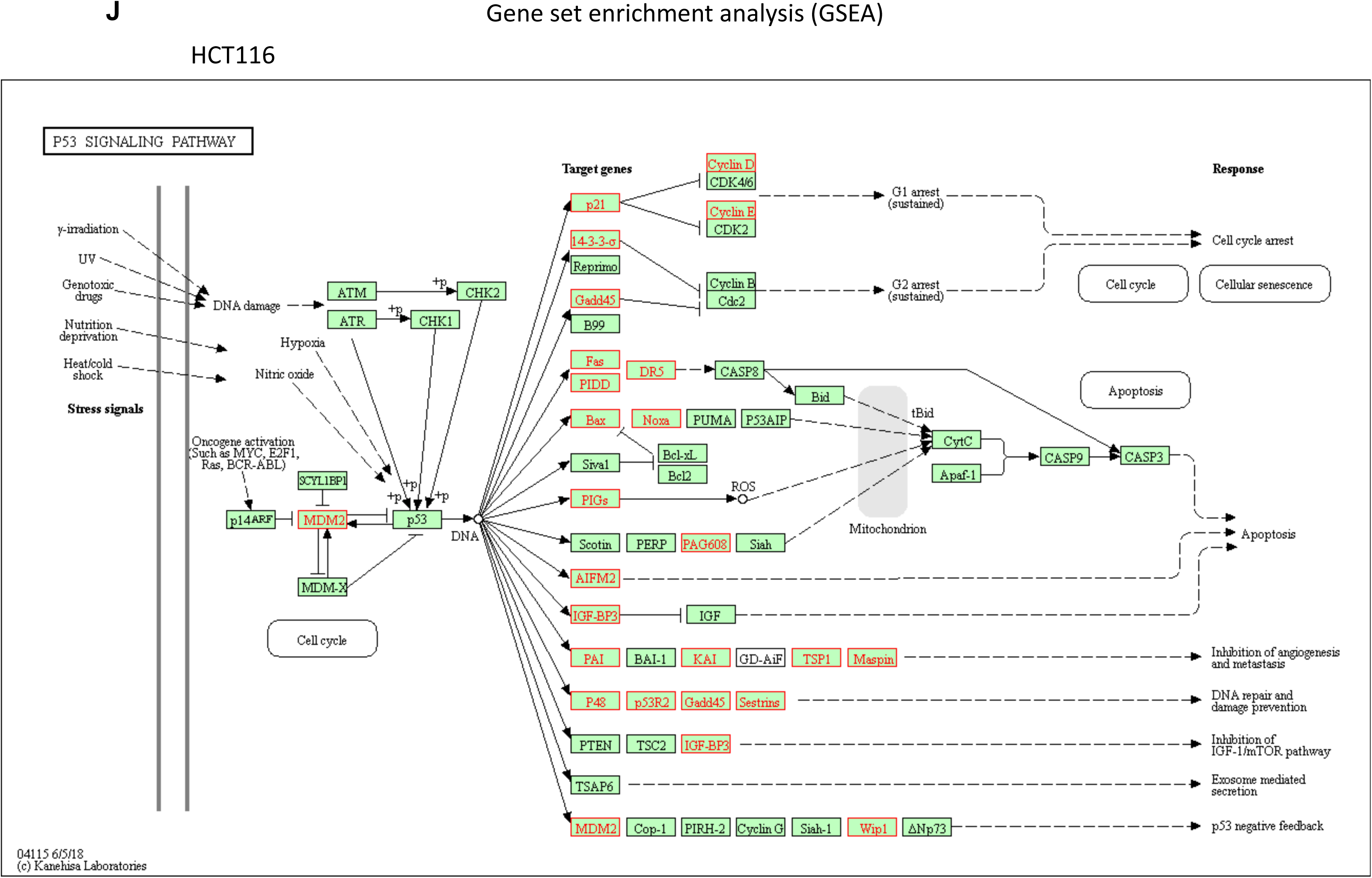
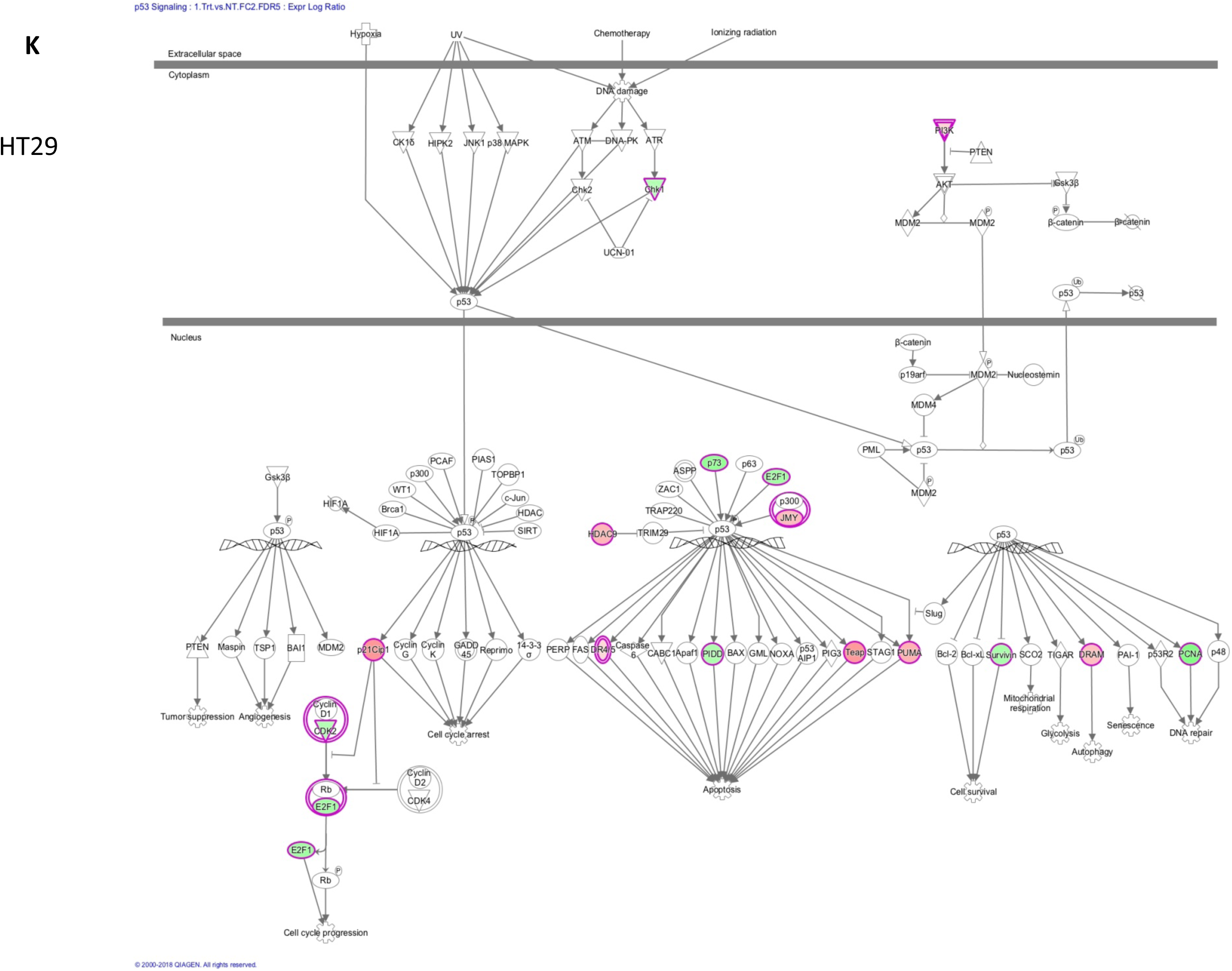
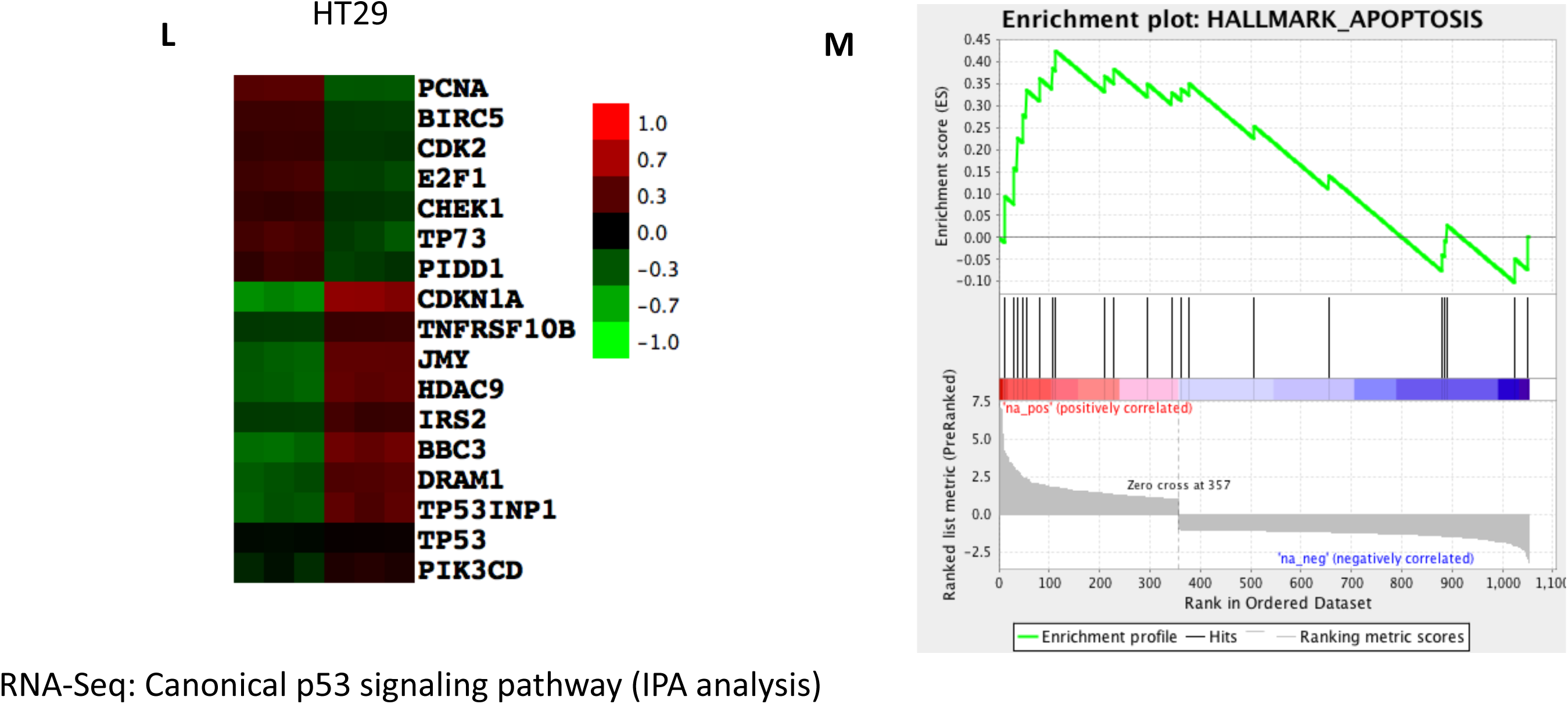
PG3-Oc partially restore p53 pathway. **(A)**. HCT116 and HCT1116 p53^-/-^ cells were treated with 50 µM 5-FU for 24 and 48 hours. Western blot was performed using indicated antibodies. (**B**, **C, D, E, F** and **G**) HCT116 and HCT1116 p53^-/-^ cells were treated with 50 µM 5-FU for 24 hours in triplicate. RNA-Seq and IPA were performed (see Materials & Methods for details). (**H, I** and **J**) GSEA analysis and enriched KEEG p53 signaling pathway. HT29 cells were treated with or without 1 µM PG3-Oc for 24 hours in triplicate, and RNA samples were prepared. RNA-Seq and IPA were performed (see Materials & Methods for details). (**K).** IPA analysis of the canonical p53 signaling pathway. (**L**) A heat-map depicts differential gene expression of genes in the canonical p53 pathway in PG3-Oc treated cells as identified by IPA analysis. **(M)** GSEA plot: Representative gene set from 1867 differential expression genes showing specific responses to the apoptosis pathway.

**Figure S3.**
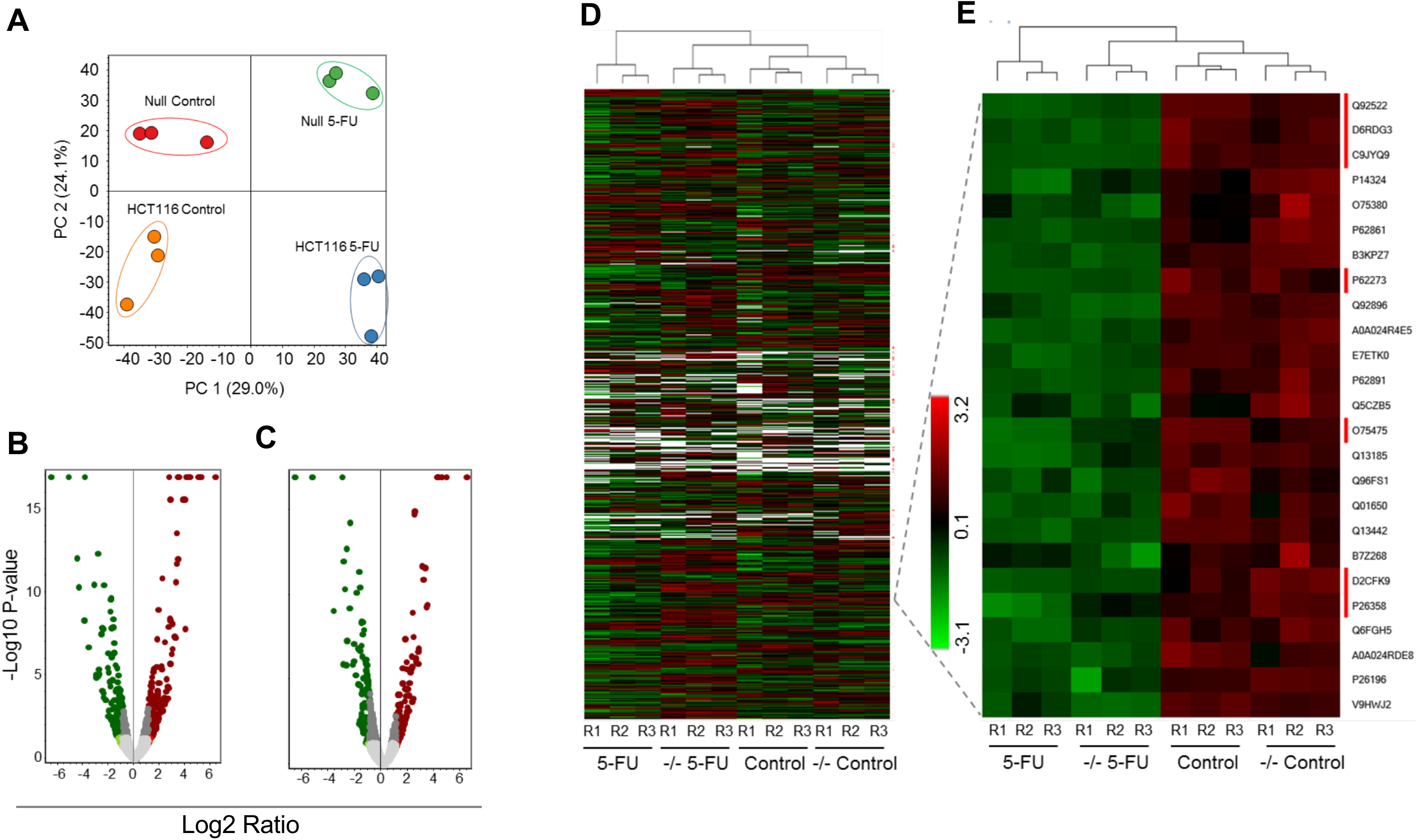
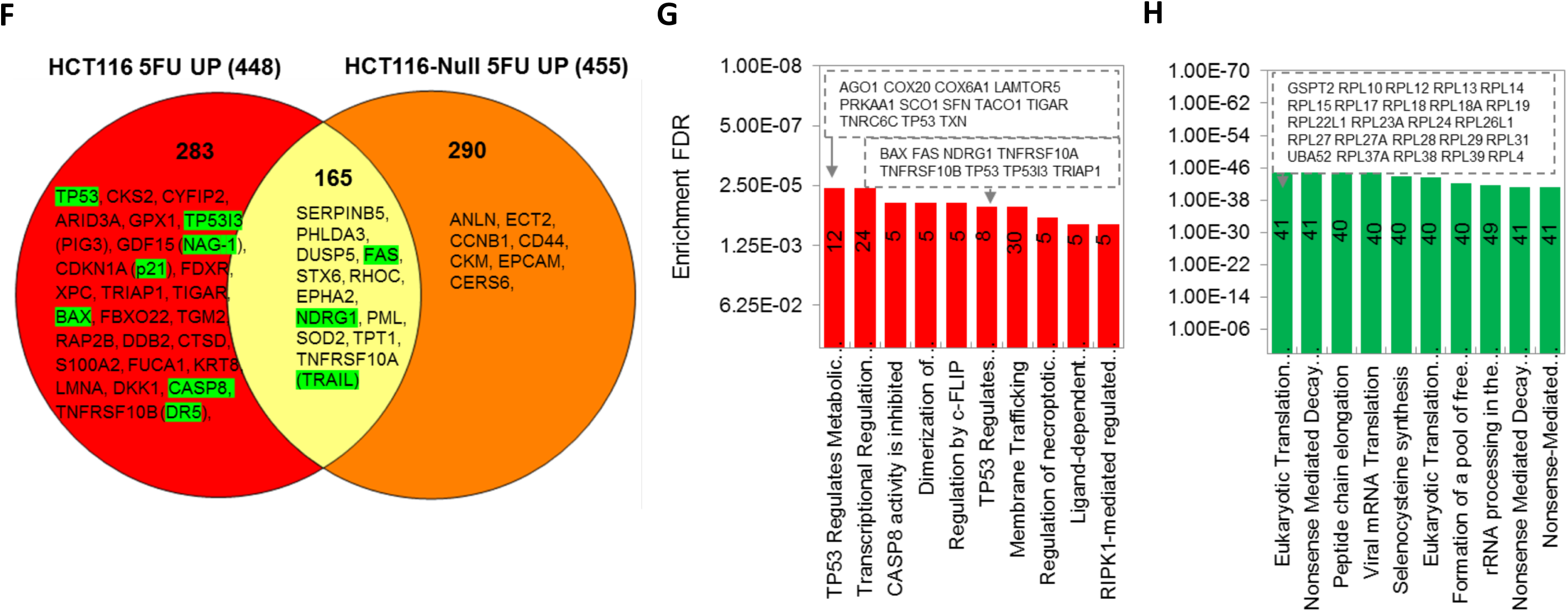
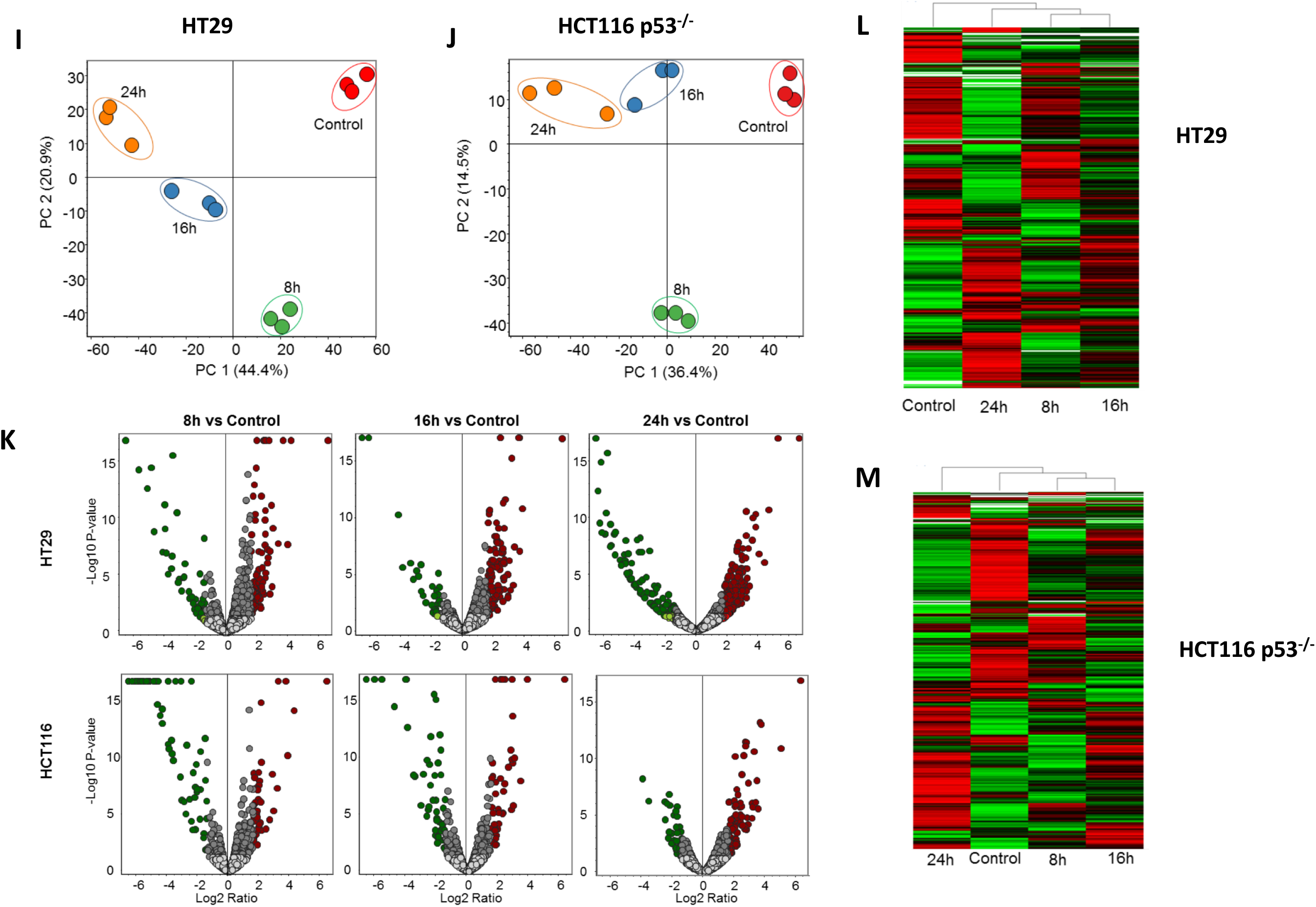
Possible p53 target genes-proteome analysis. Label-free comparative quantitative proteome analysis of HCT116 and HCT116 p53 null cell lines in response to 5-FU treatment. **A**, PCA analysis showed close clustering total normalized protein abundance of the replication in each condition, however each treatment condition is distinct from other groups. **B** and **C**, Volcano plot analysis of the significantly increased (red dot) and decreased (green dot) proteins in HCT116 and HCT116 p53 null cell lines in response to 5-Fu treatment. Gray dots are non-significant (p=0.05) and below the threshold fold change (1.5fold). **D** and **E** show the protein expression patterns of each groups. Development of p53-dependent proteome dataset. **F**, Comparative proteome analysis of up-regulated proteins in HCT116 and HCT116-p53 Null cell lines in response to 5-FU treatment. Venn diagram analysis shows the unique and overlapping up regulated proteins in HCT116 and HCT116-p53 Null cell lines. The green marked proteins are known p53-dependent proteins identified in several publications. **G** and **H**, represent the KEGG gene enrichment analysis of the unique upregulated proteins identified from HCT116 and HCT116-p53 Null cell lines. As expected, p53 signaling proteins are enriched in HCT116 cells that is consistent with that 5-FU is a positive regulator of p53 pathway. Label-free comparative quantitative proteome analysis of HCT116 p53-null and HT29 p53-mutant cell lines in response to PG3-Oc treatment exposed different time points. **I** and **J**, PCA analysis showed close clustering total normalized protein abundance of the replication in each condition, however each treatment condition is distinct from other groups. **K**, Volcano plot analysis of the significantly increased (red dot) and decreased (green dot) proteins in HCT116 p53 null and HT29 p53 mutant cell lines in response to PG3-Oc treatment. Gray dots are non-significant (p=0.05) and below the threshold fold change (1.5 fold). **L** and **M** show the protein expression patterns of HT29 and HCT116 cell lines, respectively.

**Figure S4.**
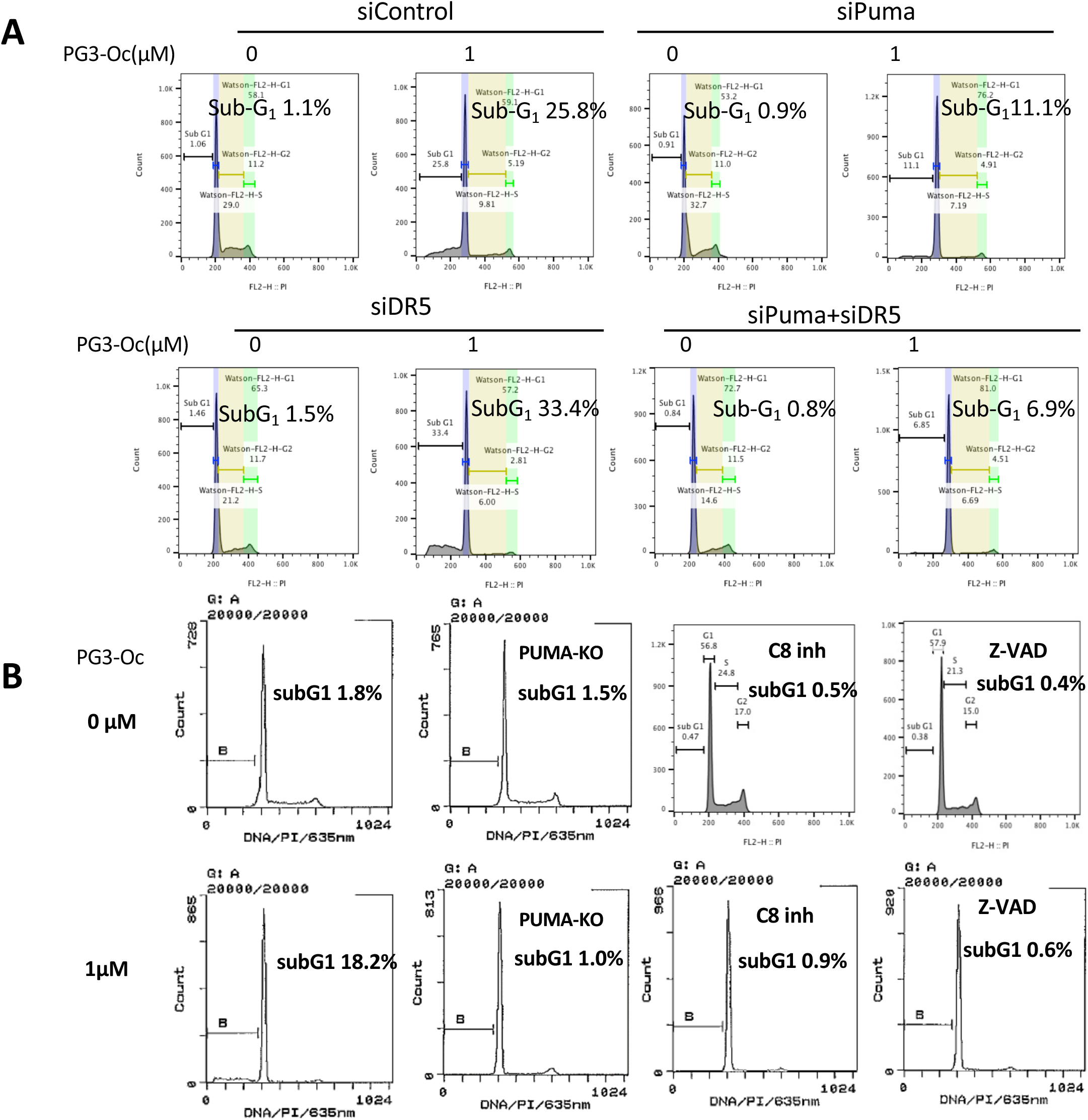
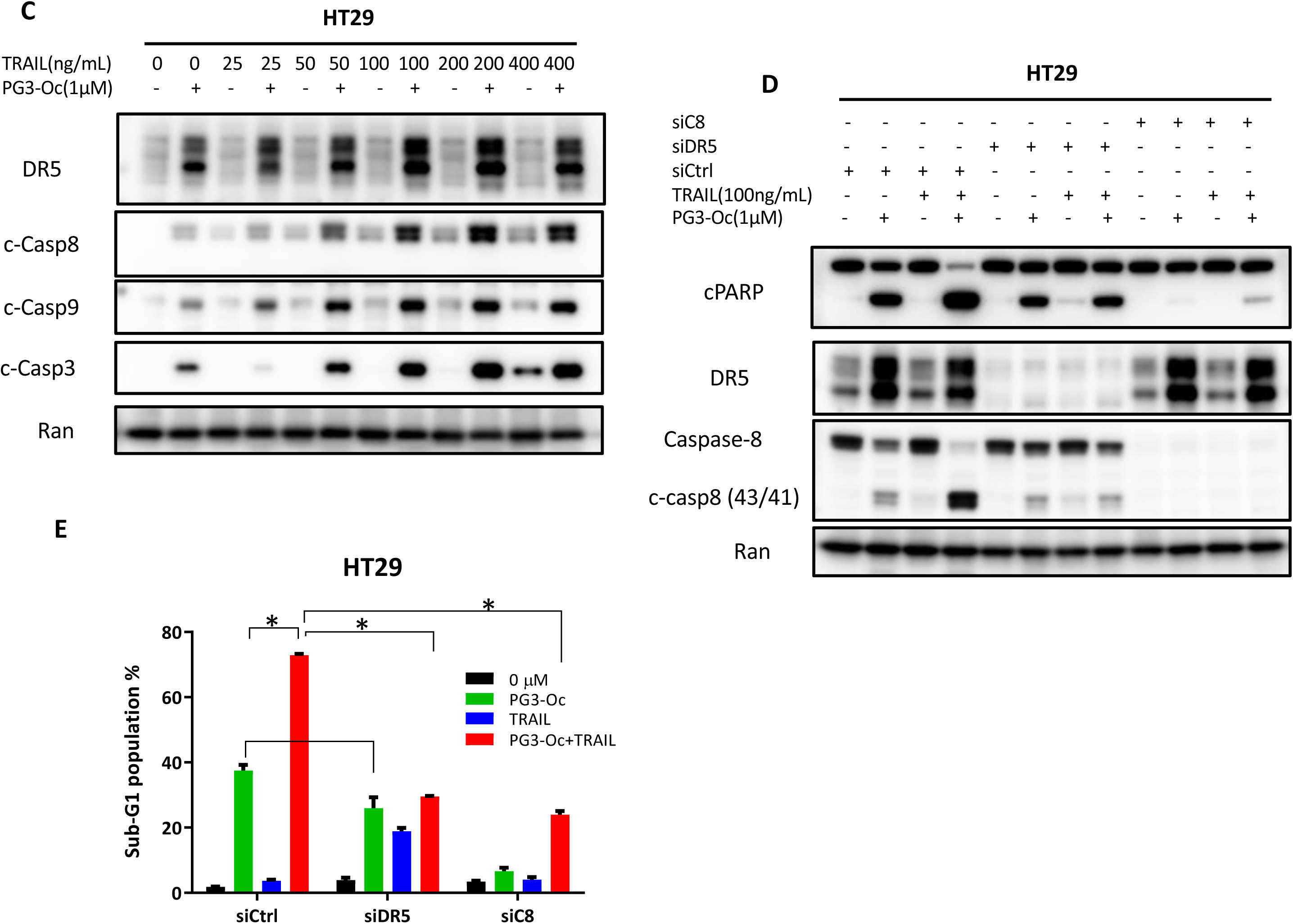
Puma is required for PG3-Oc-mediated apoptosis. **(A)** HT29 cells were transfected with Control, PUMA, DR5 and PUMA/DR5 siRNAs, and after 24 hr of transfection, the cells were treated with 1 μM PG3-Oc for 48 h. Cell death was analyzed by nuclear PI-staining using flow cytometry. (**B)** HT29 and HT29-PUMA-KO cells were treated with PG3-Oc or co-treated with caspase 8 inhibitor (cas8 inh) and pan-caspase (Z-VAD-FMK) inhibitor for 48 hours. Cell death was analyzed by nuclear PI-staining using flow cytometry. (**C)** HT29 cells were pre-treated with 1 μM PG3-Oc for 41h, and then TRAIL was added to the media at the indicated doses and incubated for 5 hours, and western blots were performed using the indicated antibodies. **(D)** HT29 cells were transfected with the indicated siRNAs. After 24 hours, cells were treated with 1 μM PG3-Oc for 41 hours, followed with addition of 100 ng/mL TRAIL for another 5 hours. Western blots were performed using the indicated antibodies. **(E)** HT29 cells were transfected with the indicated siRNAs. After 24 hours, cells were treated with 1 μM PG3-Oc for 41 hours, followed with addition of 100 ng/mL TRAIL for another 5 hours. Sub-G1 population analysis was carried out using flow cytometry.

**Figure S5.**
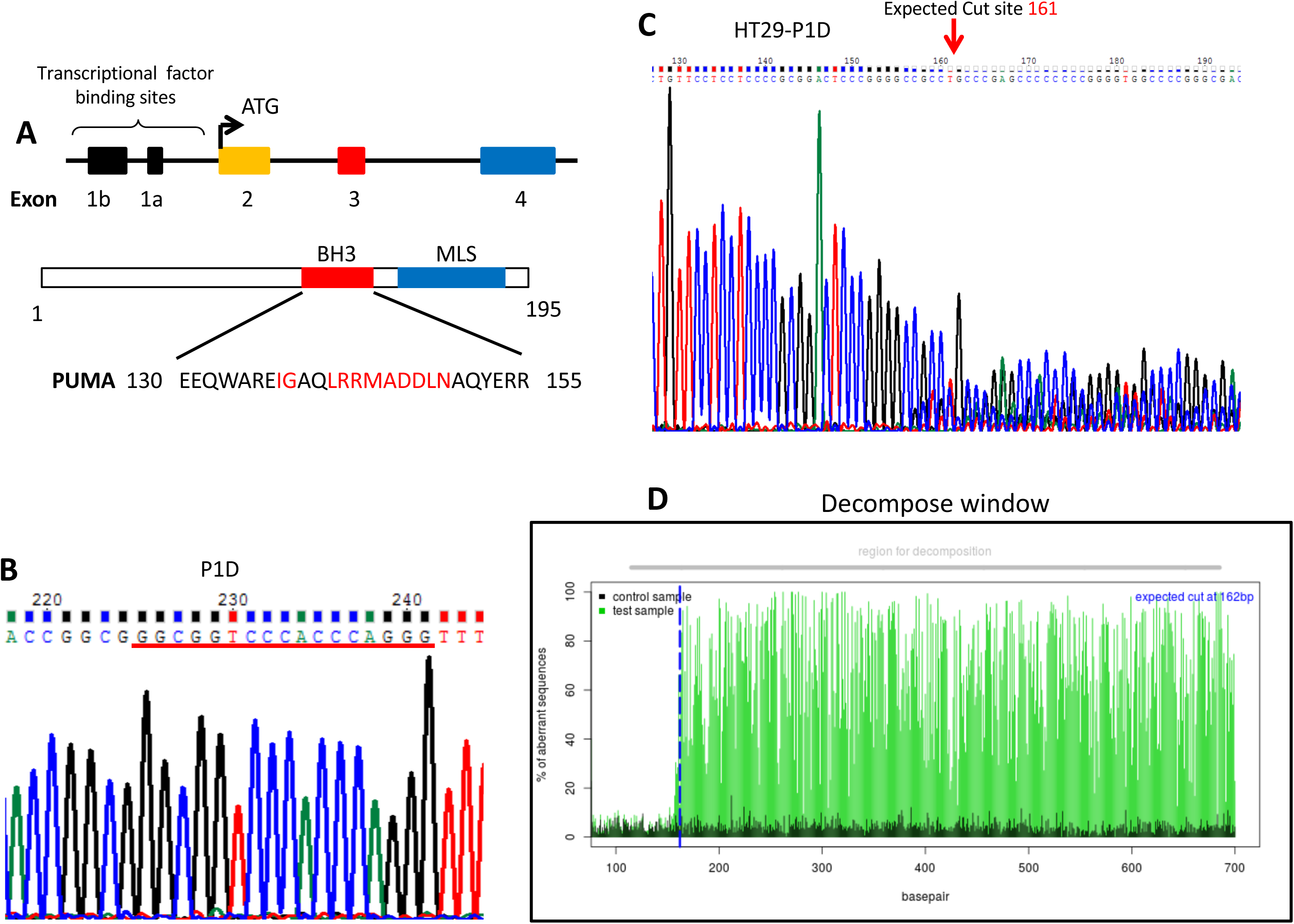
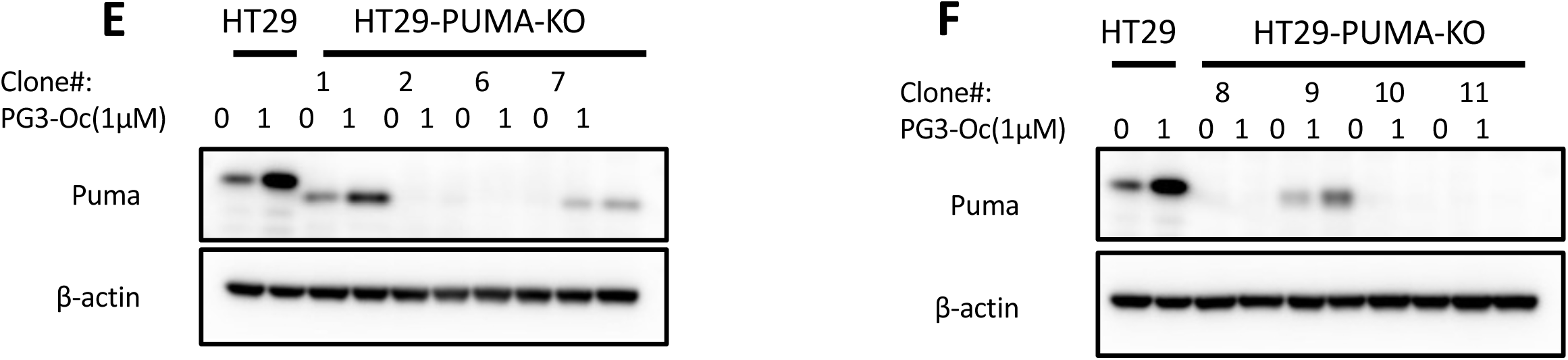
Knock-out of the *PUMA* gene by CRISPR/Cas9 gene editing in colon cancer cells. **A**. The human *PUMA* gene contains three coding exons (exons-2-4) and two non-coding exons (exons 1a and 1b). PUMA protein has two functional domains, the BH3 and C-terminal mitochondria-localization signal (MLS). The red-colored residues are conserved within other pro-apoptotic Bcl-2 family members. **B.** sequencing results of guide 1-containing plasmid P1D. **C.** DNA sequencing results of HT29-P1D, which are pools of lentivirus-infected and puromycin-selected cells. **D.** The decomposition window of TIDE analysis for HT29-P1D. **E** and **F**. Western blot analysis of the expression of PUMA protein from single cell colonies isolated from a pool of HT29-PUMA-KO cells.

**Figure S6.**
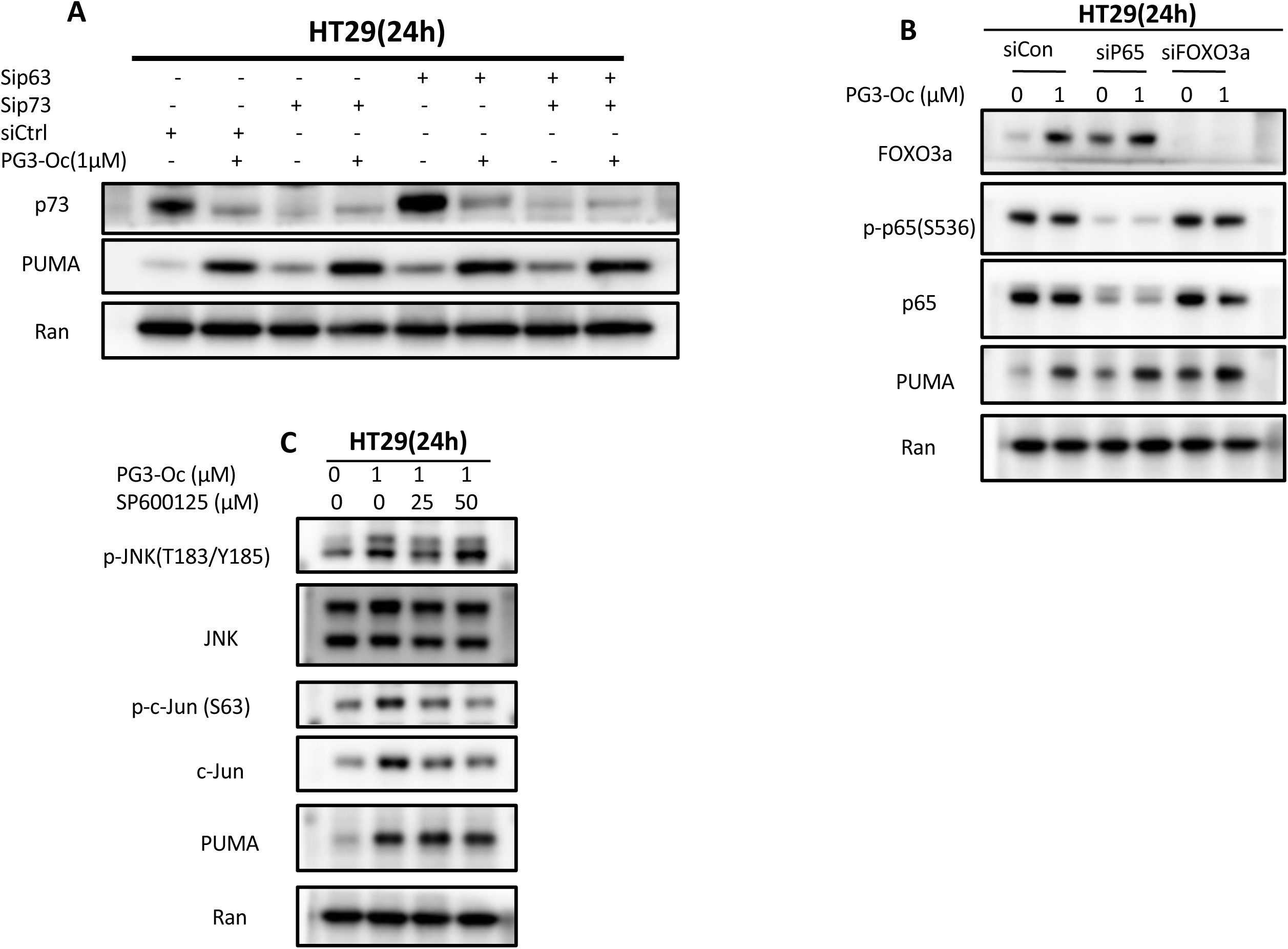
Explore molecular mechanism of PG3-Oc-induced upregulation of PUMA. (**A).** HT29 cells were transfected with p73 and p63 siRNAs for 24h, and then treated with 1 μM PG3-Oc for 24 hr. Western blots were performed using the indicated antibodies. (**B).** HT29 cells were transfected with NF-κB p65 siRNA or FOXO3a siRNAs for 24 hr, and then treated with 1 μM PG3-Oc for 24 hr. Western blots were performed with the indicated antibodies. **C**. HT29 cells were treated with PG3-Oc or co-treated with SP600125 for 24 hours, and then western blots were performed with the indicated antibodies.

**Figure S7.**
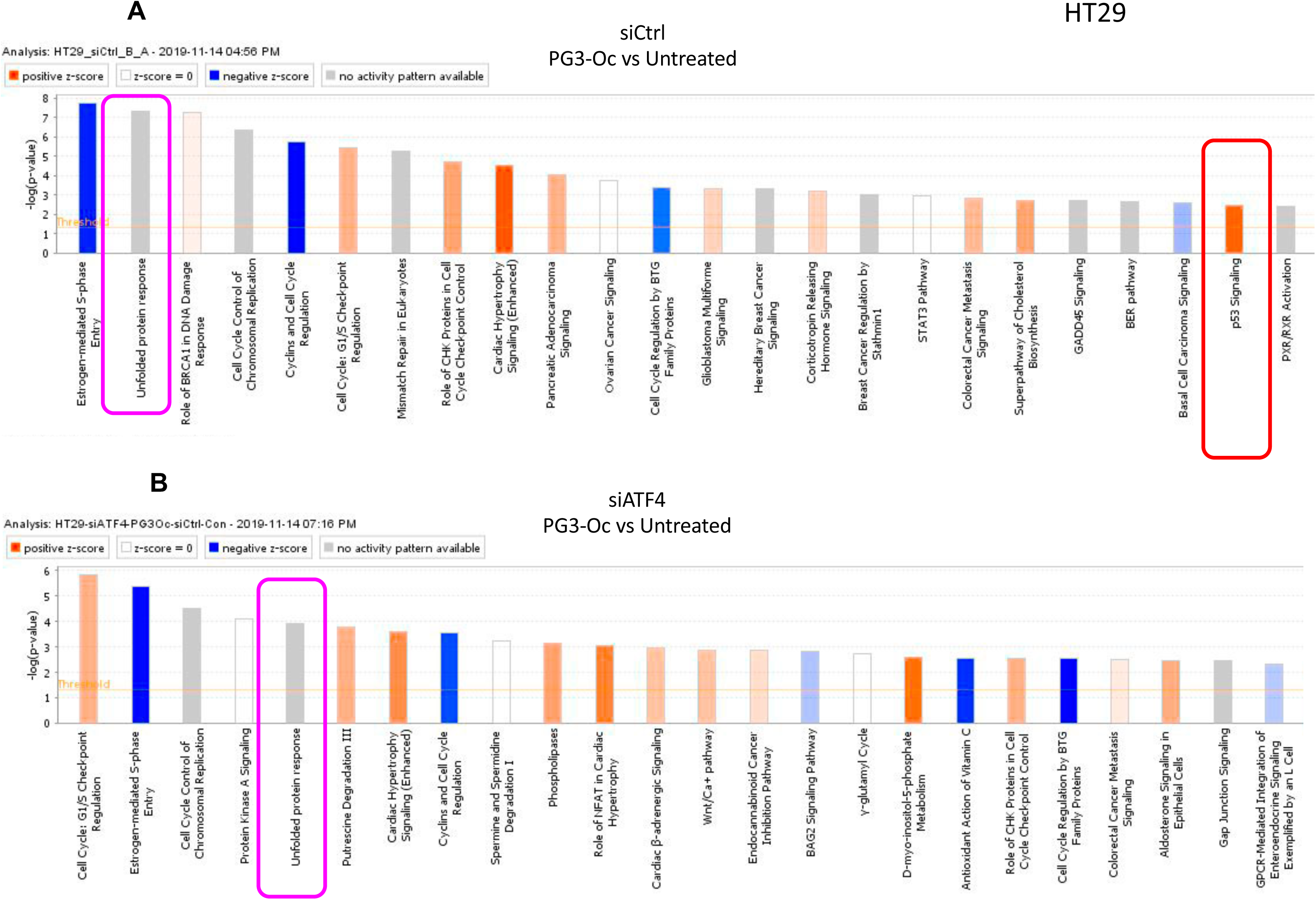
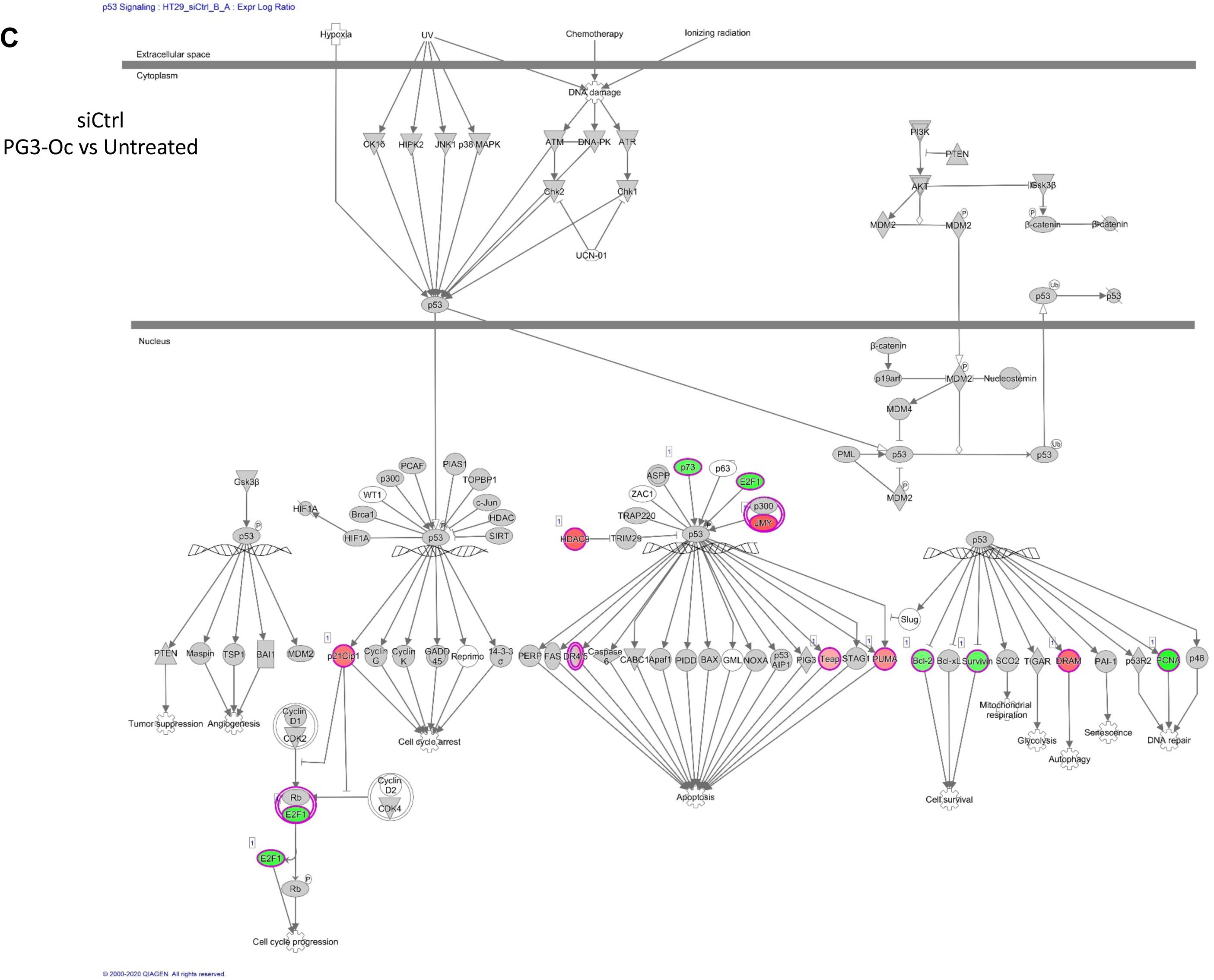
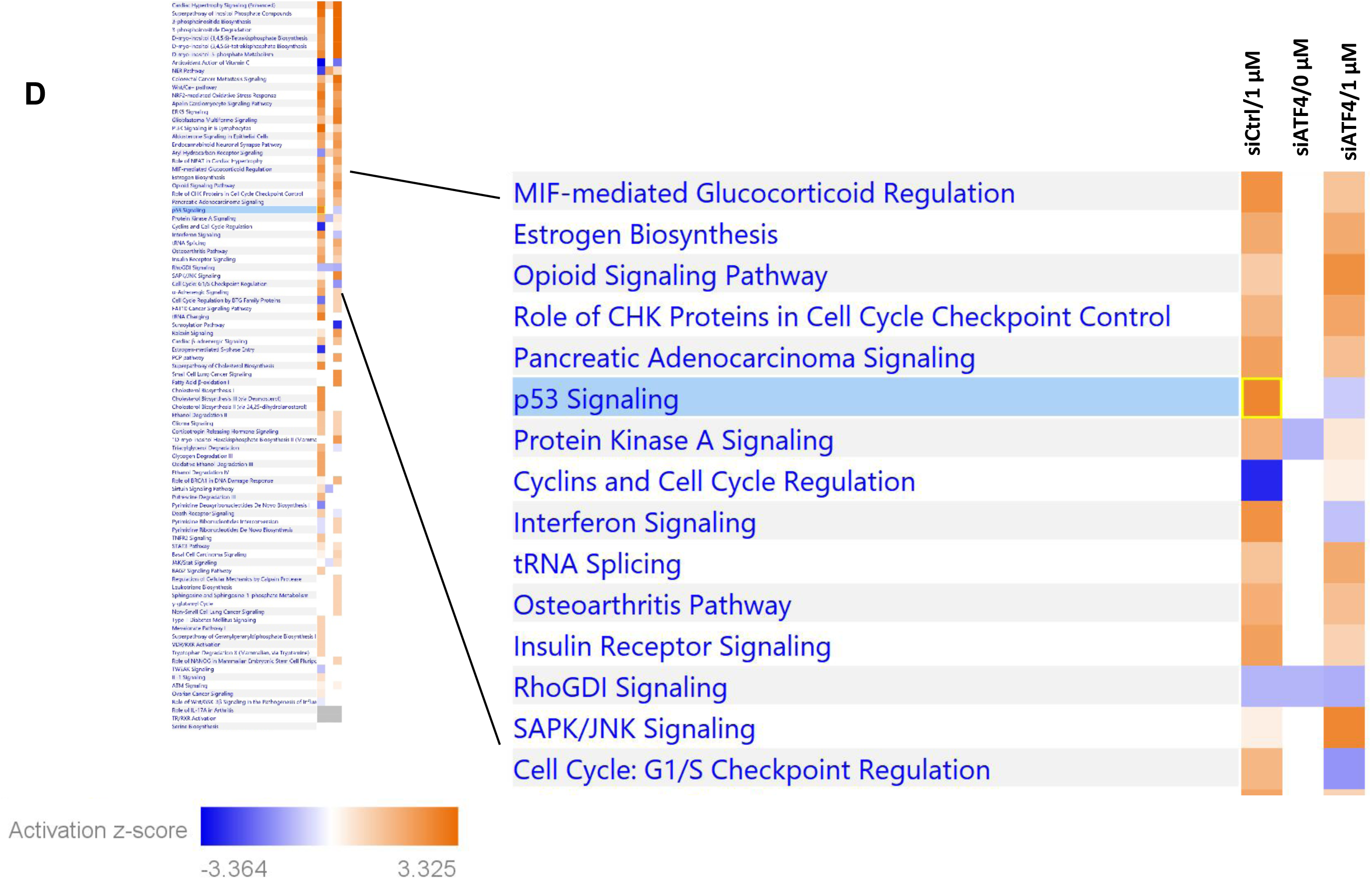
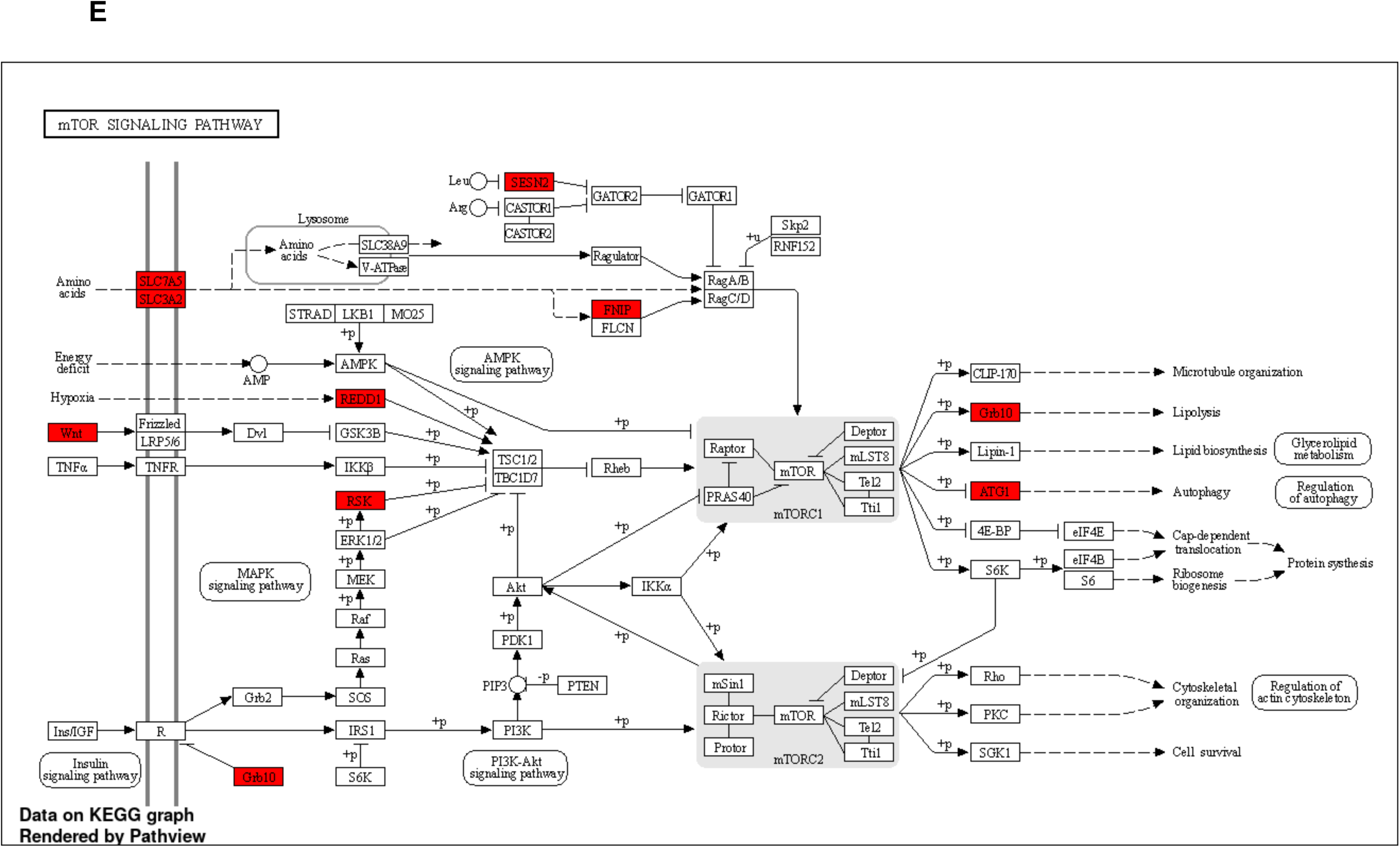
ATF4 is a possible regulator that mediates PG3-Oc induced p53 pathway restoration. HT29 cells were transfected with Control and ATF4 siRNAs, and at 24 hr after transfection, cells were treated with or without 1 µM PG3-Oc for 24 hours in triplicate, and RNA samples were prepared. RNA-Seq, and IPA analysis were performed (see Materials & Methods for details). **(A)** Canonical pathway analysis of siControl samples treated with/without PG3-Oc indicated that p53 pathway is one of hits and activated, **(B)** but not in siATF4 samples treated with/without PG3-Oc. **(C)** Canonical p53 pathway from **(A)**. **(D)** Heatmap of signaling pathways was generated by IPA analysis from three groups of samples (normalized to Control siRNA transfected sample without PG3-Oc treatment). **(E)** KEEG gene enrichment showed unique genes identified in PG3-Oc treated si-Control cell lines in comparison with known p53 gene data-base.

**Figure S8.**
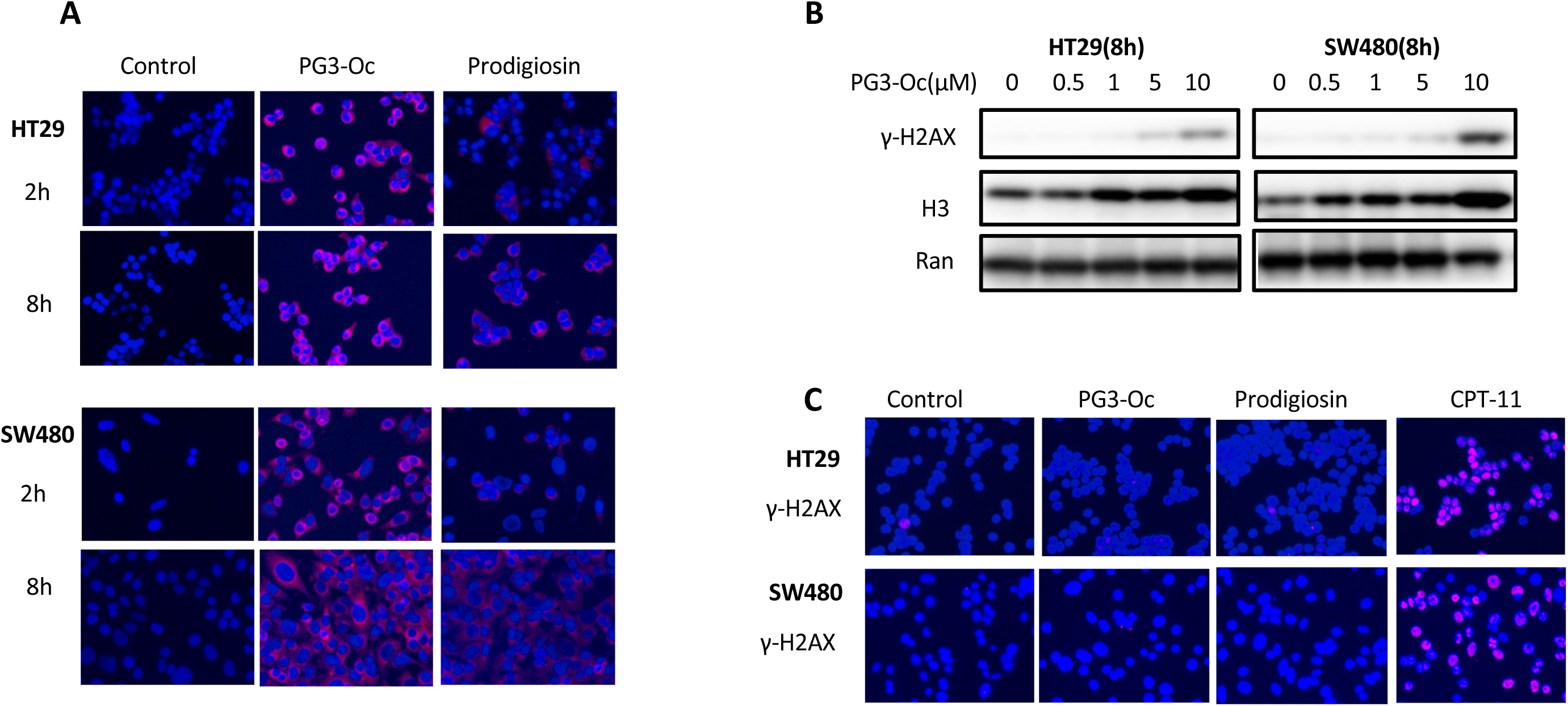
PG3-Oc induces apoptosis without genotoxicity. (**A)** Localization of PG3-Oc in cells. HT29 and SW480 cells were incubated with PG3-Oc (1 μM) for the indicated times. Cells were fixed, counterstained with DAPI, and analyzed with fluorescence microscopy. (**B)** PG3-Oc did not induce DNA damage. HT29 and SW480 cells were treated with PG3-Oc at the indicated concentrations and time. Western blots are shown for γ-H2AX, histone H3 and the loading control Ran. (**C)** Immunofluorescence staining for γ-H2AX foci. HT29 and SW480 cells were treated with PG3-Oc at the indicated concentrations and time, respectively. Cells were fixed, incubated with γ-H2AX antibody and then Cy3-secodnary antibody, counterstained with DAPI, and examined by fluorescent microscope.

**Figure S9.**
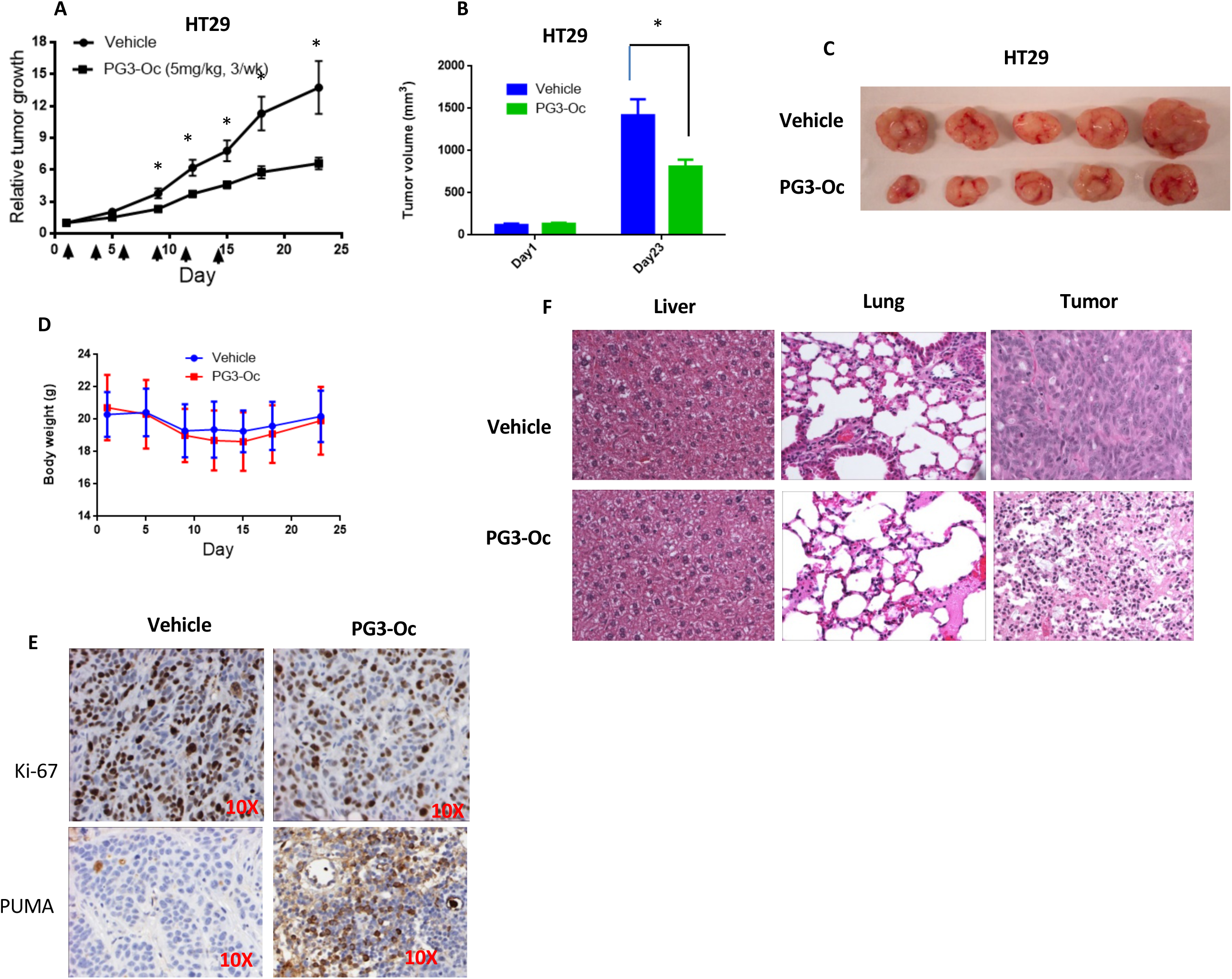
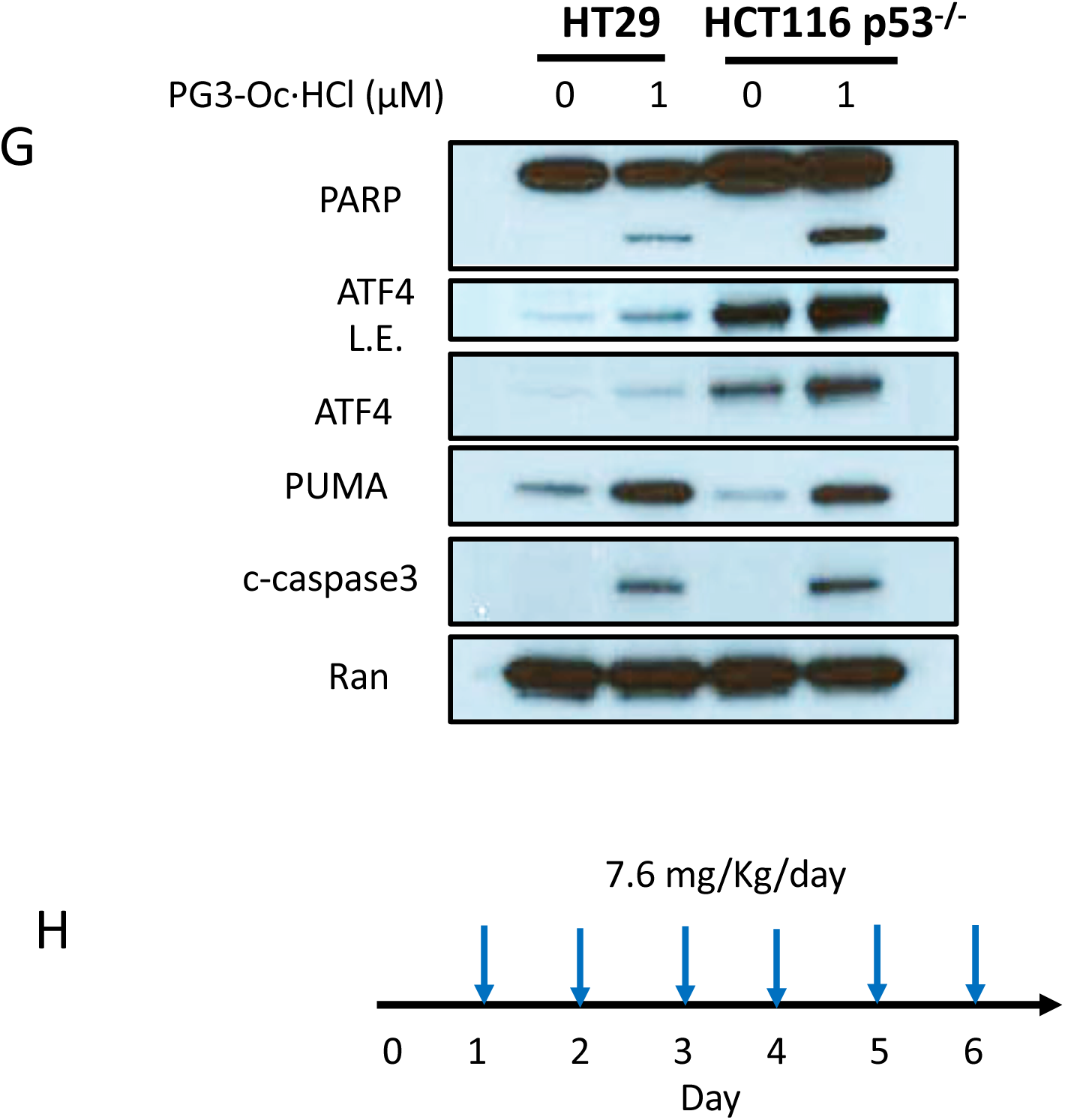
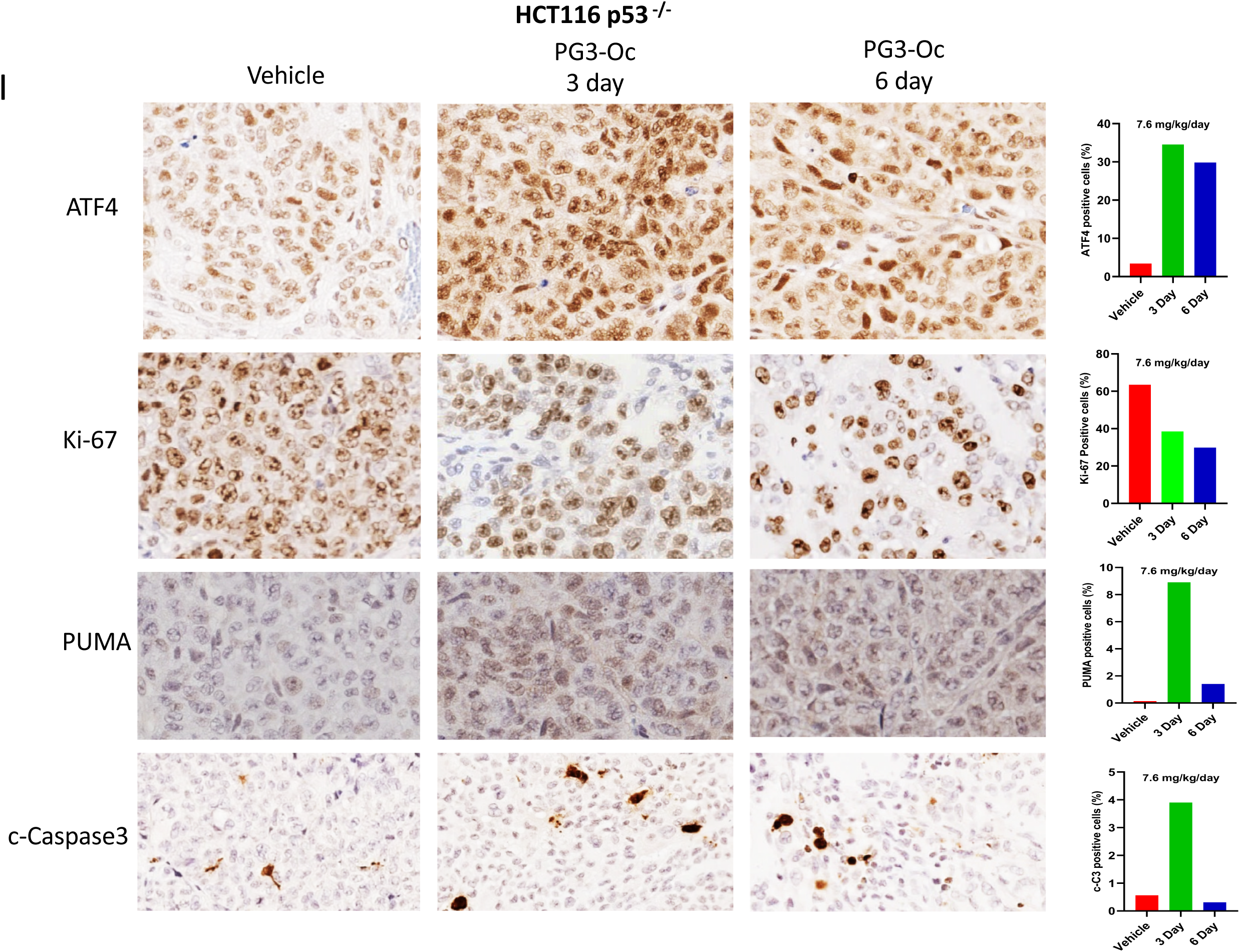
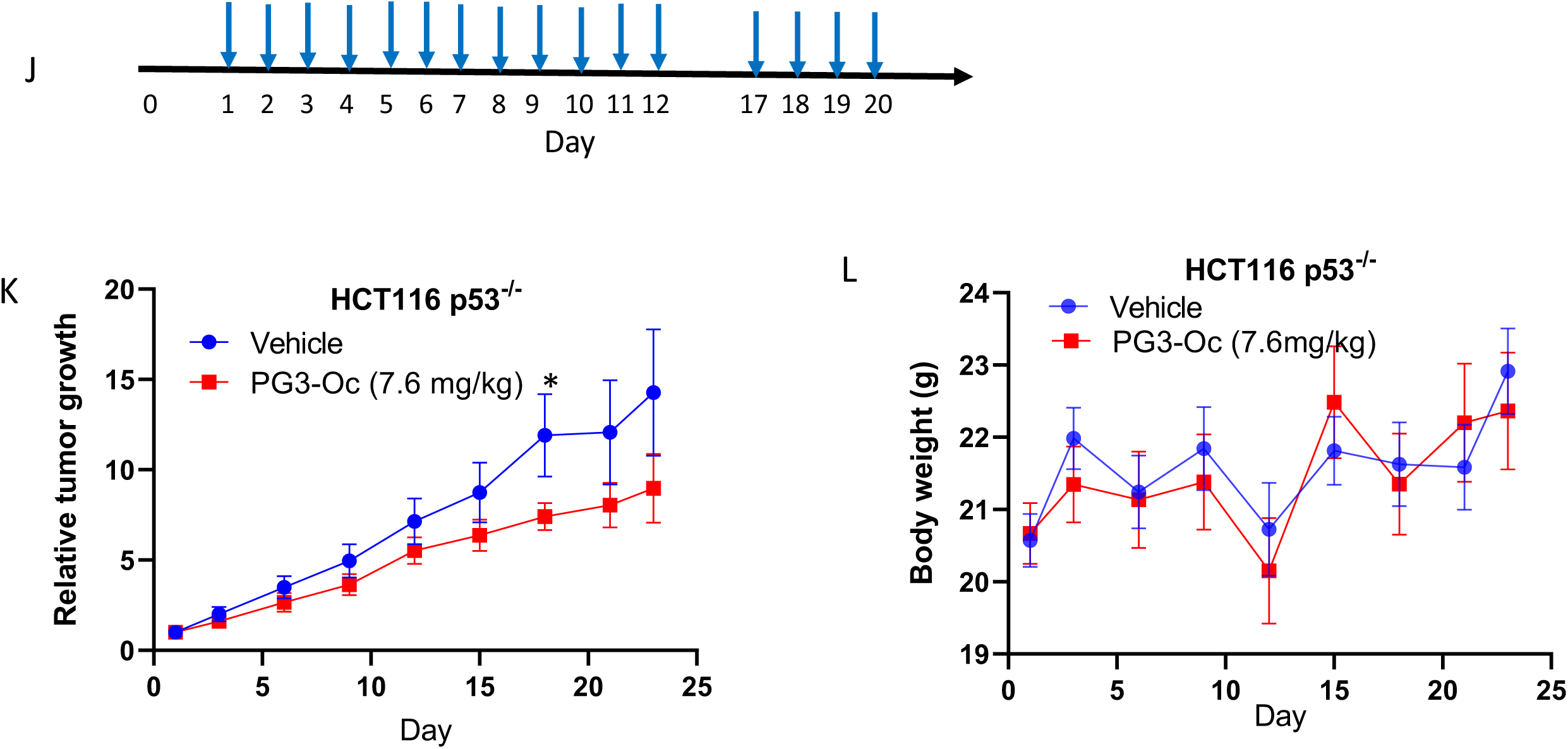
PG3-Oc inhibits tumor growth in a colon cancer xenograft mouse model. Established HT29 xenografts were treated with 5 mg/kg PG3-Oc and vehicle control three times weekly for a total 6 treatments. The arrows show the days of the treatment. **(A)** The relative tumor growth is normalized tumor size to the tumor size of day 1 before the treatment (*, p< 0.05 by an unpaired *t* test). (**B)** The mean tumor volume before and after treatment (*, p< 0.05 by an unpaired *t* test). (**C)** Images of 5 representative tumors from vehicle control and treated groups. (**D)** Body weight changes of nude mice during treatment period (*, p< 0.05 by an unpaired *t* test). **(E)** Ki-67 and PUMA antibody staining of HT29 tumors. (**F**) H&E staining of liver, lung and HT29 tumors. **(G)** HCT116 p53^-/-^ cells were treated with PG3-Oc·HCl 1µM for 48 hours, and wester blot were performed using indicated antibodies. **(H)** Scheme of the treatment of mice. **(I)** IHC and quantification of the tumor xenografts using indicated antibodies. **(J)** Scheme of the treatment of the mice. **(K)** The relative tumor growth is normalized tumor size to the tumor size of day 1 before the treatment (*, p< 0.05 by an unpaired *t* test). (**L)** Body weight changes of nude mice during treatment period (*, p< 0.05 by an unpaired *t* test).

